# Nanopore sequencing with unique molecular identifiers enables accurate mutation analysis and haplotyping in the complex Lipoprotein(a) KIV-2 VNTR

**DOI:** 10.1101/2024.03.01.582741

**Authors:** Stephan Amstler, Gertraud Streiter, Cathrin Pfurtscheller, Lukas Forer, Silvia Di Maio, Hansi Weissensteiner, Bernhard Paulweber, Sebastian Schönherr, Florian Kronenberg, Stefan Coassin

## Abstract

**Background:** Repetitive genome regions, such as variable number of tandem repeats (VNTR) or short tandem repeats (STR), are major constituents of the uncharted dark genome and evade conventional sequencing approaches. The protein-coding *LPA* kringle IV type-2 (KIV-2) VNTR (5.6 kb per unit, 1-40 units per allele) is a medically highly relevant example with a particularly intricate structure, multiple haplotypes, intragenic homologies and an intra-VNTR STR. It is the primary regulator of plasma lipoprotein(a) [Lp(a)] concentrations, an important cardiovascular risk factor. However, despite Lp(a) variance is mostly genetically determined, Lp(a) concentrations vary widely between individuals and ancestries. This VNTR region hides multiple causal variants and functional haplotypes.

**Methods:** We evaluated the performance of amplicon-based nanopore sequencing with unique molecular identifiers (UMI-ONT-Seq) for SNP detection, haplotype mapping, VNTR unit consensus sequence generation and copy number estimation via coverage-corrected haplotypes quantification in the KIV-2 VNTR. We used 15 human samples and low-level mixtures (0.5% to 5%) of KIV-2 plasmids as a validation set. We then applied UMI-ONT-Seq to extract KIV-2 VNTR haplotypes in 48 multi-ancestry 1000-Genome samples and analyzed at scale a poorly characterized STR within the KIV-2 VNTR.

**Results:** UMI-ONT-Seq detected KIV-2 SNPs down to 1% variant level with high sensitivity, specificity and precision (0.977±0.018; 1.000±0.0005; 0.993±0.02) and accurately retrieved the full-length haplotype of each VNTR unit. Human variant levels were highly correlated with next-generation sequencing (R^2^=0.983) without bias across the whole variant level range. Six reads per UMI produced sequences of each KIV-2 unit with Q40-quality. The KIV-2 repeat number determined by coverage-corrected unique haplotype counting was in close agreement with droplet digital PCR (ddPCR), with 70% of the samples falling even within the narrow confidence interval of ddPCR. We then analyzed 62,679 intra-KIV-2 STR sequences and identified ancestry-specific STR patterns. Finally, we characterized the KIV-2 haplotype patterns across multiple ancestries.

**Conclusions:** UMI-ONT-Seq accurately retrieves the SNP haplotype and precisely quantifies the VNTR copy number of each repeat unit of the complex KIV-2 VNTR region across multiple ancestries. This study utilizes the KIV-2 VNTR, presenting a novel and potent tool for comprehensive characterization of medically relevant complex genome regions at scale.

## Introduction

Complex genome regions such as copy number variations (CNVs), variable number of tandem repeats (VNTR), short tandem repeats (STRs), and centromeric and telomeric regions can hide medically highly relevant mutations that shed new light on human phenotypes and diseases[1–6]. The development of long-read sequencing technologies and new bioinformatic tools have greatly improved the resolution of these difficult regions[7–14]. However, some overly complex repeat structures remain challenging. The *LPA* KIV-2 VNTR is a medically highly relevant protein-coding example of such a difficult region[15].

The *LPA* gene encodes the apolipoprotein(a) [apo(a)] and controls most (>90%) of the lipoprotein(a) [Lp(a)] plasma variation[16]. High Lp(a) plasma concentrations are considered a nearly monogenically determined, very frequent, causal, independent and heritable risk factor for atherosclerotic cardiovascular diseases [17–19] that increase cardiovascular risk up to threefold[20, 21]. Elevated Lp(a) concentrations are found in ≈20% of White individuals and even in ≈50% of individuals of African ancestry[17]. Median Lp(a) levels vary tenfold between ancestries[22] and the individual plasma concentrations vary even 1000-fold[16]. The causes of this huge phenotypic variance are not fully understood but likely result from intricate, haplotype-dependent, non-linear interactions between multiple functional *LPA* variants and the KIV-2 VNTR size[15].

The complex structure of the *LPA* gene severely complicates genetic analysis[15]. It comprises ten highly homologous kringle-IV domains (KIV-1 to -10)[23, 24]. Each KIV domain consists of two short exons (≈160 and 182 bp) interspaced by a mostly ≈4 kb intra-kringle intron and a 1.2 kb intron linking it to the next domain[15]. The KIV-2 domain is encoded by a polymorphic VNTR, which introduces 1 to ≈40 KIV-2 units per gene allele (5.6 kb per repeat unit)[23]. This creates an up to >200 kb VNTR region consisting of highly homologous coding repeat units that encompass up to 70 % of the protein[25]. The VNTR copy number explains 30-70 % of Lp(a) variance in a non-linear, ancestry-dependent manner[16]. Individuals carrying at least one low molecular weight (LMW) apo(a) isoform (defined as 10-22 KIV units[16], i.e. 1 to 13 KIV-2 units[15]) present 5 to 10 times higher *median* Lp(a) levels compared to high molecular weight isoforms (>22 KIV; HMW) due to higher protein production rates[17]. However, the *individual* Lp(a) levels within the same isoforms can still vary 10 to 200-fold[15] due to multiple, partially unknown genetic variants that modify the effect of the VNTR[15].

The interactions between the KIV-2 VNTR size and modifier SNPs are complex and multilayered (reviewed in detail in [15]). They are haplotype-dependent and only partially captured by linkage disequilibrium (LD)[15]. Indeed, several functional SNPs, including the two SNPs (4925G>A[25] and 4733G>A[26]). These two variants alone explain remarkable 11.9% of the Lp(a) variance in the general population, are ancestry-specific, are associated with considerably reduced cardiovascular risk and were hidden in the KIV-2 VNTR until recently[25–28]. The background apo(a) isoform size and other variants on the same haplotype can both limit and amplify the effects of any functional variant[15]. Although the KIV-2 VNTR encompasses most of the *LPA* gene region, the full genetic and haplotypic diversity of KIV-2 units and the LD of KIV-2 haplotypes with the haplotypes in the non-repetitive kringles remain largely unknown.

Current short-read deep sequencing approaches confidently identify KIV-2 SNPs[24], but mostly lose the long-range SNP haplotype data. Early cloning studies identified three synonymous KIV-2 haplotypes named KIV-2A, KIV-2B and KIV-2C[29, 30]. These KIV-2 subtypes are defined by the haplotype of three SNPs in KIV-2 exon 1 and differ by about 120 bases[23, 24]. The KIV-2 subtypes have no functional relevance, but their frequencies differ widely between ancestries and correlate with known differences of the Lp(a) phenotype across ancestries[24, 30]. They may thus reflect distinct evolutionary histories of the KIV-2 region and tag novel ancestry-specific functional variants. Further haplotypic effects in the KIV-2 VNTR are unknown and could not be studied so far.

Nanopore sequencing (ONT-Seq; Oxford Nanopore Technologies, ONT) provides novel means to address this knowledge gap. DNA is sequenced by monitoring the sequence-specific current fluctuations generated by single-stranded DNA molecules translocating through protein pores[31, 32]. This generates hundred times longer read lengths than short-read next-generation sequencing (NGS)[14, 32] and provides single molecule resolution. It thus allows retrieving the complete haplotype of each DNA molecule sequenced, even in DNA mixtures[32]. However, at the single molecule level the benefits of nanopore sequencing are limited by its relatively high raw-read error rate (0.7-1% median error rate per read). Especially in highly similar repeat sequences like the KIV-2, errors cannot be polished by sequencing deeper (because the parental repeat of each read is unknown[33]) or by using double-stranded (“duplex”) basecalling (because of erroneous matching of strands originating from different parental molecules).

Coupling of ONT-Seq with unique molecular identifiers (UMI-ONT-Seq) allows lowering the ONT-Seq error rate considerably[33, 34]. UMIs are oligonucleotide libraries that randomly tag each template molecule with a unique identifier (Figure 1). The tagged library is amplified by PCR to generate multiple copies of each UMI-tagged template molecule and full-length sequenced[34]. The reads are clustered according to terminal UMI combination, which reflects their original template, and a consensus sequence is generated for each UMI cluster. This removes PCR and sequencing errors[34] (Figure 1), while preserving the complete SNP haplotype of each input molecule. In highly repetitive and homologous regions such as the *LPA* KIV-2 repeats, this finally provides highly accurate consensus sequences of each repeat unit.

**Figure 1:**
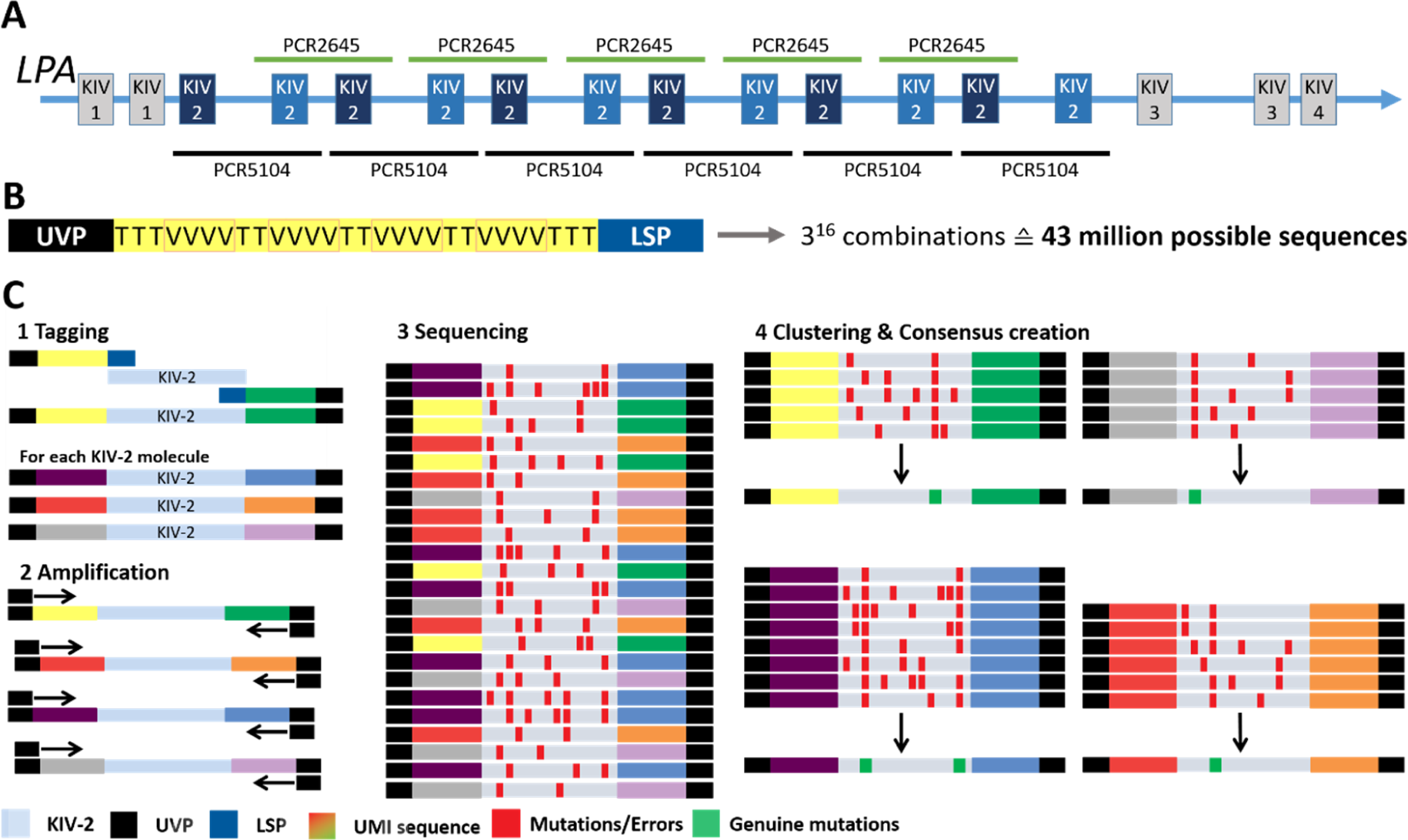
Technical aspects: *LPA* structure, amplicon location and UMI design. **Panel A:** Partial *LPA* gene structure (second exon of KIV-4, KIV-5 to KIV-10 and the C-terminal protease domain omitted) and amplicon location. **Panel B:** UMI-ONT-Seq primer design, including a universal primer (UVP) binding site for amplification of the tagged KIV-2 repeats, the UMI sequence and a locus-specific primer sequence (LSP) (e.g. KIV-2 specific). The UMI sequence leads to about 43 million possible unique tagging sequences. **Panel C:** The LSP primer site is used to tag specifically each KIV-2 repeat in a sample with a unique UMI sequence (1). . Subsequent amplification with a universal primer (UVP) (2) and sequencing (3) causes random errors that cannot be differentiated from genuine low-level variants (red boxes in 3). The UMI-sequences are used to cluster all sequences originating from one KIV-2 repeat unit and create molecule-wise consensus sequences. This removes errors that occur only in subset of sequences in each UMI cluster, but retains genuine variants that occur in a majority of the sequences in within each UMI cluster.

We describe a comprehensive assessment of UMI-ONT-Seq for SNP detection, SNP haplotyping, generation of consensus sequences for each VNTR unit and copy number determination by coverage-corrected quantification of the unique haplotypes, using the *LPA* KIV-2 VNTR region as an example for a medically highly relevant complex VNTR region. We created a scalable freely available UMI-ONT-Seq Nextflow analysis pipeline that can be generalized to also any other UMI-ONT-Seq experiment (https://github.com/genepi/umi-pipeline-nf) and demonstrate *LPA* KIV-2 haplotyping by UMI-ONT-Seq in 48 multi-ancestry samples from the 1000 Genomes[35] (1000G) Project.

## Methods

### KIV-2 amplicons and reference sequence

All KIV-2 VNTR units were amplified using two amplicons that amplify all KIV-2 units as an overlapping amplicon mixture[24, 25, 30] (Figure 1A). The PCR5104 amplicon spans one KIV-2 unit and portions of the inter-kringle intron. The PCR2645 amplicon spans one inter-KIV-2 intron with the flanking exons and parts of the intra-kringle intron. Experimental conditions are given in Supplementary table 1 and Supplementary table 2. All positions and fragment lengths in this manuscript are based on the reference sequences for a single KIV-2 unit used in [24].

### Recombinant standards

Generation of the recombinant KIV-2A and KIV-2B standards has been described before[24, 25]. The PCR5104 plasmids differ by 87 positions from each other (including four insertions/deletions [indels]; excluding a microsatellite region in the intra-KIV-2 intron of LPA5104)[24]. The PCR2645 plasmids differ at 120 positions to each other. The plasmids also differ from the respective reference sequences at 28 (PCR5104 KIV-2A), 85 (PCR5104 KIV-2B), 8 (PCR2645 KIV-2A) and 116 (PCR2645 KIV-2B; 113 being in the first intron) positions. Mimicking the in-vivo situation, where the KIV-2B represents always the minor component in the KIV-2 VNTR[24, 30], we generated five mixtures of KIV-2B plasmids in KIV-2A background, ranging from 5% to 0.5% KIV-2B. All variants present on the same plasmid represent one haplotype and should be thus detected at the same level (Supplementary figure 1). Mixture accuracy was validated by ddPCR (Biorad QX200) (Supplementary table 3).

### Human samples and KIV-2 ddPCR

Sixteen unrelated samples from the healthy-working population SAPHIR (Salzburg Atherosclerosis Prevention Program in subjects at High Individual Risk; sample codes EUR in figures and tables)[36] with KIV-2 SNP data from ultra-deep NGS from Coassin and Schönherr et al, 2019[24] were used to evaluate UMI-ONT-Seq performance in human samples using amplicon PCR5104. The KIV-2 repeat number was quantified by ddPCR (Supplementary Methods, Supplementary table 4). Mean confidence interval (CI) width of ddPCR KIV-2 quantification was as low as 4.81±2.34 KIV-2 copies. One sample was excluded due to technical failure. 48 multi-ancestry samples (Yoruba [YRI], Dai Chinese [CDX], Japanese [JPT], Punjabi [PJL]; 12 samples per group) were obtained from the Coriell 1000G samples repository. The 1000G samples are available at the Coriell Sample Repository (Camden, NJ, USA). The SAPHIR study was approved by the ethical committee of the Land Salzburg and all participants provided an informed consent.

### KIV-2 UMI-ONT-Seq principle

UMI-ONT-Seq follows the approach developed by Karst et al[34] using the oligo design from ONT technical note CPU_9107_v109_revC_09Oct2020 (generating 3^16^ diverse oligos; Figure 1B). The UMI primers consist of a 3’ locus-specific primer (LSP)[24], the UMI and a 5’ universal priming site (UVP)(Figure 1B). All PCRs were performed with the ThermoFisher Platinum SuperFi II ultra-high fidelity polymerase (accuracy >100-fold over Taq[37, 38]). About 2 ng gDNA (i.e. 50,000 KIV-2 template copies under assumption of maximum 80 KIV-2 genomic repeats; observation from our database with >13,000 apo(a) Western blots[39] and from [40–42]) were tagged with two UMI-PCR cycles (Figure 1C1) followed by New England Biolabs ExoCIP treatment. The tagged templates were then amplified in two rounds of PCR with UVP primers (early and late UVP-PCR) to produce multiple copies of each tagged molecule (Figure 1C2). A 0.9x SPRI beads purification between the two rounds of UVP-PCR was used to reduce background (Supplementary Methods). After full-length nanopore sequencing (Figure 1C3), reads are clustered based on the terminal UMI pairs and a consensus sequence is created for every UMI cluster with a defined minimal read number (UMI cluster size; Figure 1C4). Each consensus sequence represents the sequence of a single genomic KIV-2 VNTR repeat unit.

### Sequencing and basecalling

All samples were full-length sequenced on a MinION 1B system (ONT, Oxford, UK) with either the earlier R9 chemistry (SQK-LSK109 with native barcoding kits NBD-104 and NBD-114 and R9.4.1/MIN106D flow cells) or the newer V14 chemistry (SQK-NBD114 and R10.4.1/MIN114 flow cells). About 100,000 on-target, full-length reads were generated per sample (Supplementary table 5 to Supplementary table 9). This was sufficient to recall all expected variants in preliminary experiments with plasmid standards (both unmixed plasmids and mixtures at 1% and 5%) and two human samples. Basecalling was done using guppy v6.3.8 with either “high accuracy” (HAC; for R9 and V14) or “super accuracy” (SUP; for V14) settings. The 15 human PCR5104 SUP samples were additionally duplex basecalled (SUPDUP; for V14) using ONT duplex tools v0.2.20[43]. The 1000G dataset was basecalled with dorado v.0.5.1[44] and SUP algorithm.

### UMI-ONT-Seq Nextflow analysis pipeline

Initially, our analysis pipeline corresponded to the proof-of-principle UMI analysis pipeline published online by ONT[45] (which follows the pipeline steps of Karst et al[34]), but was migrated to the Nextflow framework with several optimizations to improve performance and parallelization. The pipeline steps are described in Supplementary figure 2 and in the Supplementary Methods. In brief, all reads of each barcode are merged, filtered for length (>1000 bp) and mean per-base quality (>Q9) and aligned to the target reference sequence. Only primary alignments with more than 95% overlap are retained to remove chimeric amplicons. The reads are clustered according to the terminal UMI sequences using vsearch[46] and only UMI clusters with ≥20 reads were retained for consensus sequence generation (as recommended by Karst et al.[34]). Each cluster is then polished using Medaka v1.7.0. The UMI extraction, clustering and reference alignment steps are repeated for the polished consensus sequences to generate the final consensus sequences (clustering step 2).

Extensive analysis of the migrated pipeline revealed inaccurate clustering by vsearch (see Results section for details). In both clustering steps, vsearch clustered distant UMI combinations and separated identical UMI combination into different clusters. We therefore modified the clustering strategy of the pipeline. Looser clustering parameters (80 % sequence identity) in clustering step 1 prevent separation of identical UMI sequences. To prevent mixing of distant UMIs into one cluster, all clusters containing more than 12 (R9 HAC), 10 (V14 HAC), 8 (V14 SUP) reads were then split by taking the first UMI sequence of the cluster file and clustering it with all remaining UMI sequences in the same file that show ≤2 bp edit distance (UMI collision probability <0.01 %). The remaining UMIs were treated as a new cluster and clustered iteratively. The minimal UMI cluster size required for consensus creation was derived based on plateauing of the consensus quality at these values (see Results). Subsequently, in clustering step 2 stringent clustering parameters (>99 % sequence identity) were used to remove PCR duplicates without mixing distinct UMI clusters.

Our analysis pipeline and test data is available at https://github.com/genepi/umi-pipeline-nf under GNU General Public License v3.0. For reproducibility, we provide also the migrated ONT pipeline at https://github.com/genepi/umi-pipeline-nf/tree/default_clustering_strategy.

### UMI-ONT-Seq residual error rate estimation

Sequencing the unmixed plasmids provides an intuitive way to estimate the residual error rate of the UMI-ONT-Seq as any variation in the consensus sequences can be considered an error. The error rate can be quantified by either averaging the Phred (Q-) scores of each consensus sequence (“consensus sequence Q-Score”) and or dataset-wide (“dataset Q-Score”). The latter was introduced because the fragment length limits the maximum achievable consensus-sequence Q-Score and because perfect consensus sequences produce infinite Q-Score. The dataset Q-Score was defined as *Error rate = n_differences_/n_sequences_ × length_sequences_* with the Q-score being *Q = -10 × log_10_(error rate)*[47].

### Variant calling

Variants in the UMI consensus sequences were called using a modified version of Mutserve[48] v2.0.0-rc13[49] that does not require bidirectional confirmation. The minimum KIV-2 mutation level that needs to be detected is 1.25% (1 in ≈80 KIV-2 units)[25]. To allow for random fluctuations and sequence-context effects, variants >0.85% variant level were considered true. Variant KIV-2 calling without UMIs was done by omitting the UMI clustering. This resembles ultra-deep KIV-2 NGS sequencing as done in [24, 25]. All reads were aligned to the reference sequence for one repeat unit and conventional low-level variant calling was performed using default Mutserve v2.0.0-rc15[49] and ClairS-TO[50]. Extraction of the intronic KIV-2 STR sequences and variants for each KIV-2 unit is described in the Supplementary methods.

### Variant truth set

Ultra-deep targeted KIV-2 NGS data obtained previously for all SAPHIR samples[24, 25] was used as truth set for UMI-ONT-Seq evaluation. The 1000G variant truth set was generated using a KIV-2 NGS variant calling pipeline on the 1000G 30X whole-genome (WGS) sequencing data[51] (with minor adaptions for WGS data). All UMI-ONT-Seq datasets were benchmarked in terms of sensitivity (true positive rate), specificity (true negative rate), precision (positive predictive value, i.e. the proportion of genuine variants among all variants found) and F1 score (harmonic mean of sensitivity and precision). For the polymorphic intronic short tandem repeat (STR) in PCR5104 (position 2472-2506) no NGS reference data was available and it thus was analyzed separately.

### KIV-2 units haplotype extraction

Haplotypes were extracted using a three-step algorithm (available at https://github.com/AmstlerStephan/haplotyping-KIV2-nf). 1) Extraction of uniquely occurring haplotypes including all positions that were called as variants in any of the samples (unique haplotypes). 2) Noise polishing and removal of “unlikely” haplotypes (merged haplotypes). 3) Estimating the repeat number per haplotype by coverage correction (coverage-corrected haplotypes).

The unique haplotypes were determined by extracting from the consensus sequences of each sample the base present at any polymorphic position in the dataset, as found by mutserve variant calling. At positions per sample with a variant frequency <0.85 % only the major variant was used in the haplotype. Next, only the uniquely occurring haplotypes per sample were retained, including their occurrence in the consensus sequences (unique haplotypes).

To obtain the merged haplotypes, the residual noise was reduced by clustering identical haplotypes and assigning each haplotype cluster below the threshold *n_sequences_* x 0.0085 to the haplotype cluster with the smallest edit distance, but not more than a maximum edit distance of 1. Next, assuming unbiased and random tagging of KIV-2 repeats in our samples and unbiased PCR amplification, we applied a binomial distribution to determine the minimal occurrence required for a haplotype to be considered genuine. The binomial distribution formula in this context is expressed as:

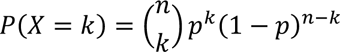

Here, *n* represents the total number of haplotypes observed after the UMI sequencing, *p* is the probability of a haplotype to occur and *k* the minimal occurrence threshold for a haplotype to be considered genuine. The probability is set to 0 (stringent filtering) to calculate the sample-specific minimal occurrence threshold (*k*) for any given haplotype. Haplotypes falling below the determined minimal occurrence threshold *k* have 0 probability to be genuine (being e.g. generated by the residual noise in the UMI sequencing) and were therefore excluded from the analysis.

For all remaining haplotypes, the number of occurrences is adjusted for potential identical KIV-2 repeats by dividing the occurrence of each haplotype by the median occurrence (sample-wise) and rounding that to the next integer. This gives the coverage-corrected number of KIV-2 repeats (coverage-corrected haplotypes).

### Data processing

Data processing and visualization was done in R 4.3.1 with ggplot2 v3.3.6 or in bash scripts (available at https://github.com/AmstlerStephan/UMI-ONT-Seq_Analysis). Reading and manipulation of BAM and FASTQ/A files was done using BioStrings v2.68.1. R-squared values were calculated using the linear model (lm) function. Bland-Altman plots were generated using BlandAltmanLeh v0.3.1. The coverage-corrected haplotypes were used to generate unrooted KIV-2 phylograms using MAFFT v7.520[52] with global iterative refinement (G-INS-i) and modified UPGMA guide trees. Visualization was done with ggtree v.3.8.2[53].

## Results

We performed a comprehensive evaluation of the performance of UMI-ONT-Seq for variant calling, haplotyping, generation of consensus sequences for each VNTR unit and VNTR copy number determination in the complex *LPA* KIV-2 VNTR[15]. We generated 28 sequencing libraries encompassing both PCR products of two unmixed plasmid standards differing by 87 (PCR5104) and 120 (PCR2645) bases, five plasmid mixtures (ddPCR-validated, Supplementary table 3) and two sequencing chemistries (R9, V14). Moreover, the KIV-2-spanning PCR5104 amplicon was sequenced on 15 human validation gDNA samples and finally used to call mutations, map haplotypes, and quantify the genomic KIV-2 units in 48 1000G samples from 4 different populations.

### UMI-ONT-Seq recapitulates expected mutation levels and KIV-2B haplotype fractions in plasmid mixtures

The switch from R9 to V14 chemistry, as well as basecalling V14 data with SUP or SUPDUP instead of HAC had, as expected, a large impact on sequencing accuracy (Supplementary table 5). For technical performance values across all experiments see Supplementary table 6 to Supplementary table 9. Variant calling performance of UMI-ONT-Seq was excellent in all plasmid mixtures down to 1% for both amplicons (Figure 2A, for R9 data see Supplementary figure 3A). Notably, no performance difference was seen for V14 data between HAC and the computationally much more expensive SUP basecalling. Specificity was 99.9 to 100% in all samples (Supplementary table 10), but a residual background noise at 0.2% to 0.5% variant level was observed in all plasmid mixtures and sequencing chemistries (Figure 2B, Supplementary figure 3B). Therefore, we introduced a cut-off of 0.85% for all further experiments, which includes all bona-fide KIV-2 variants while allowing some technical variation.

**Figure 2:**
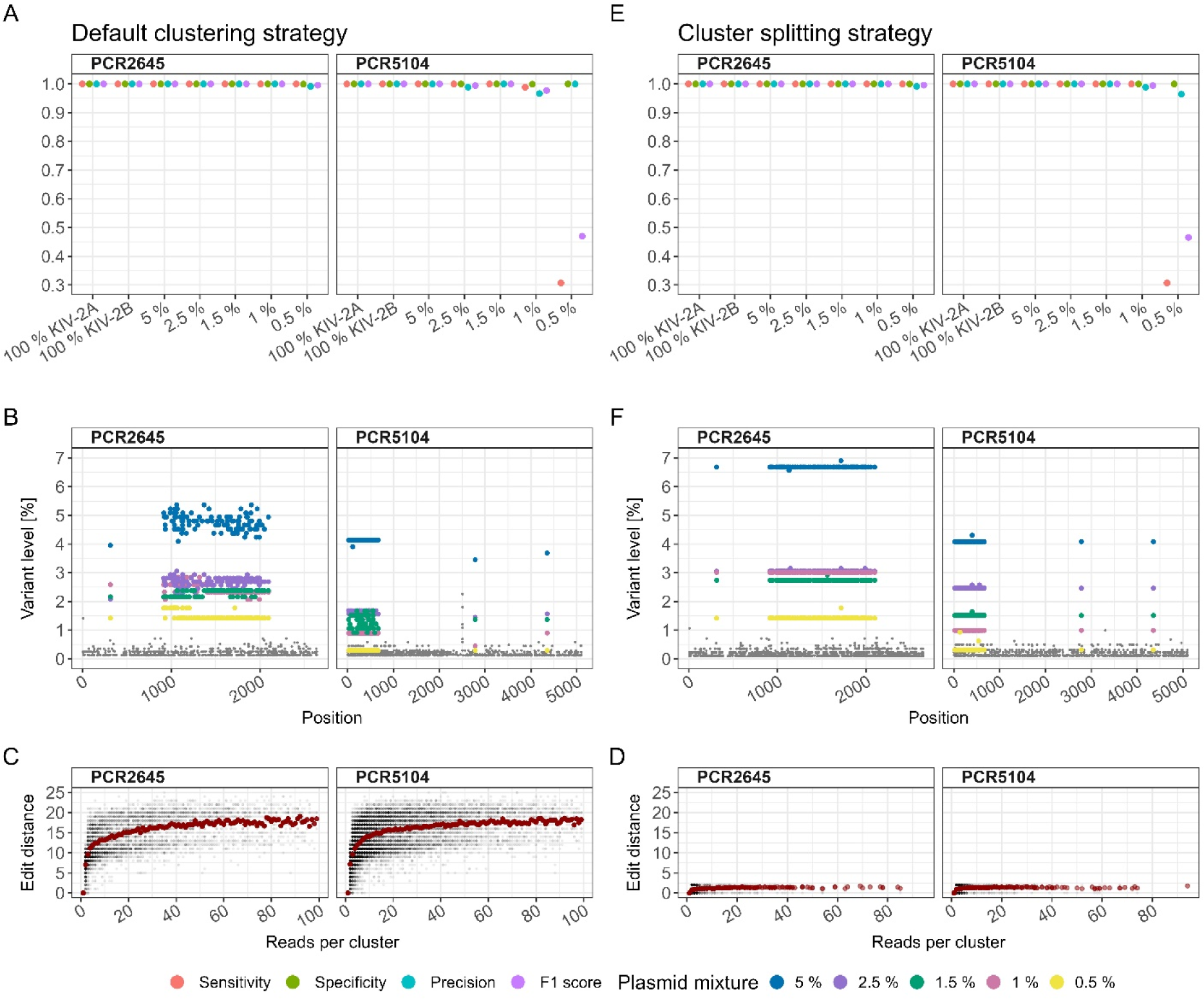
Variant detection in plasmid mixtures with the ONT UMI analysis pipeline using the default clustering strategy and the cluster splitting strategy for the V14 chemistry and HAC basecalling. **Panels A to C** show the results for the default clustering strategy, **panels D-F** show the corresponding results for cluster-splitting strategy. **Panels A and E**: Performance measures for the default clustering strategy in recalling variants of the two unmixed plasmids and plasmid mixtures from 5 to 0.5 % KIV-2B in KIV-2A background of two fragments (PCR2645, PCR5104). **Panels B and F:** variant levels for the plasmid mixtures from 0.5 – 5 % across every position of both fragments. Grey points: Low-level residual noise. Variation of detected variant levels of up to ±1 % in absolute values was observed (blue points, 5 % mixture). **Panels C and D**: Edit distance of the UMI-sequences for different UMI cluster sizes. Grey: single clusters. Red: Average cluster size. Using the default clustering strategy admixing of UMI-sequences up to an edit distance of 25 within one cluster (i.e. the sequences did not originate from the same KIV-2 repeat) led to considerable variance in the observed variant levels (panel C). Using the cluster splitting strategy with maximal edit distance 2 between the UMI-sequences (D) reduced the variant level noise considerably (E, colored points) for both fragments and all mixture levels.

Since the variants present on the two plasmids represent distinct haplotypes, we expected all mutations from the same plasmid to occur at the same level. We found that all mutations from the same plasmid were, indeed, close to the expected level on average, but showed considerable per-position noise when analyzed with the ONT UMI analysis pipeline using the default clustering strategy (Figure 2B). Systematic analysis of the UMI clusters revealed that inaccurate clustering of the UMI sequences by vsearch resulted in partially heterogeneous UMI clusters (Figure 2C, Supplementary table 11). If the UMI clusters contained a mixture of sequences (e.g. KIV-2A and KIV-2B sequences), the cluster-polishing step produced noisy mutation levels (Figure 2B, Supplementary table 12).

Implementation of the cluster splitting strategy reduced the edit distance in the UMI clusters considerably (Figure 2D). This had no impact on variant calling performance in the V14 chemistry (Figure 2E, Supplementary table 13) and most importantly, mutations originating from the same plasmid now showed virtually no residual variation and matched the expected values very accurately (Figure 2F). The number of plasmid standards with no variant level noise increased from 4/14 to 12/14 samples in HAC and SUP. The median variant level noise was reduced by 3.1-fold and 2.3-fold for V14 HAC and SUP (R9 HAC: 0.8-fold, Supplementary table 14). This indicates that our cluster splitting strategy allows accurate recalling of the haplotype of each read (Supplementary table 15). All further results are based on the UMI-ONT-Seq analysis pipeline using the cluster splitting strategy.

### UMI-ONT-Seq produces highly accurate KIV-2 consensus sequences at ≥6 reads per UMI cluster

We investigated the relationship between the UMI cluster size and consensus sequence qualities using the unmixed plasmids (KIV-2A and KIV-2B), where it can be assumed that any difference between the consensus sequences represents a PCR or sequencing error.

Up to 10 reads per UMI cluster, the dataset Q-score increased rapidly in the V14 data, reaching Q40 already at 6-10 reads per cluster (Figure 3A, Supplementary table 16; 14 reads for R9 data, Supplementary figure 4A). The consensus sequence Q-score increased in a similar manner as the dataset Q-score, reaching the maximal quality after 6 reads per cluster for the V14 chemistry (14 for R9 HAC; Supplementary figure 5A). At 6 reads per cluster already 96.8 % (HAC) and 98.3 % (SUP) of all consensus sequences showed no more than 2 errors, and 58.1 % (HAC) and 62.5 % (SUP) were even error-free (Figure 3B; R9: 79.6 % and 33.5 %; Supplementary table 17 and Supplementary figure 5B).

**Figure 3:**
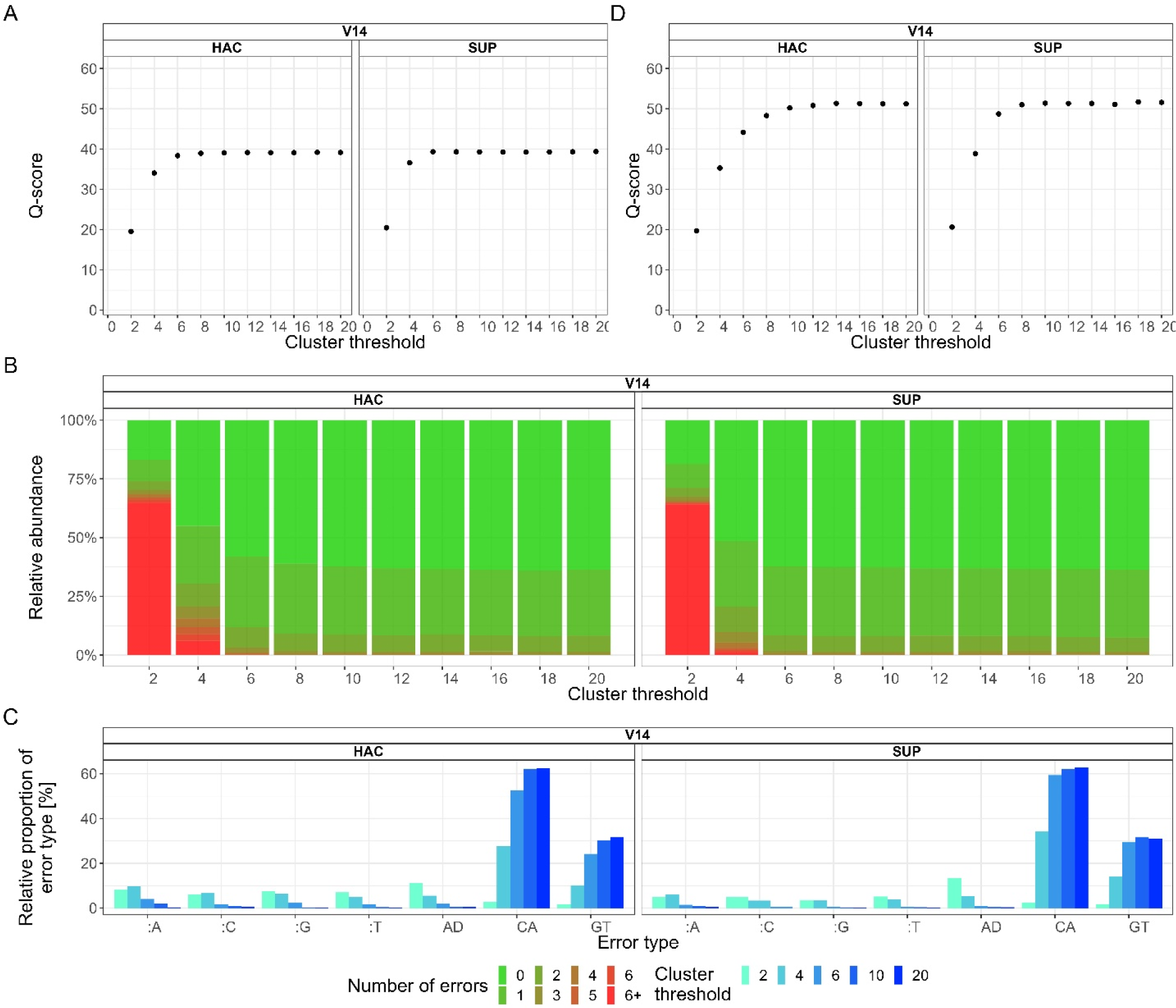
Impact of the minimal cluster size on consensus sequence quality and error-profile. **Panel A:** Dependency of the dataset Q-score from the minimal cluster size. V14 HAC and V14 SUP dataset Q-score increases rapidly being close to Q40 already at cluster size 6 to 10. **Panel B:** Percentage of perfect reads depending on the minimal cluster size threshold. At Q40 dataset Q-score 62 % of all consensus sequences are error-free and 98.5 % of all cluster had no more than 2 errors for both V14 conditions. **Panel C:** Error type frequency for V14 HAC and SUP. C to A and G to T transversions were the most common errors. “:N” denotes insertions D denotes deletions. **Panel D:** Dataset Q-score at different cluster thresholds after filtering for the sequencing chemistry-specific errors. The dataset Q-score of V14 consensus sequences reaches Q50 already at a cluster size of 6 to 10.

While indels were the most prominent error type in older R9 chemistry (Supplementary figure 4B), both V14 conditions were primarily characterized by C to A (62.40 % V14 HAC; 63.60 % V14 SUP) and G to T (31.80 % V14 HAC; 31.30 % V14 SUP) transversions (Figure 3C, Supplementary table 18). Disregarding these systematic errors, which were all below the 0.85 % mutation level, further improved the consensus sequences qualities for both V14 kits considerably (Figure 3D). Both basecalling algorithms again reached dataset Q-score Q40 at 6 reads per cluster and even Q50 at 10 and 8 reads (V14 HAC: 92 errors in 10 Mb; V14 SUP: 94 errors in 11.9 Mb; Supplementary table 19). Already at cluster size 6, 89.9% to 95.5% of all consensus sequences were error-free (Supplementary figure 5; Supplementary table 16).

### Accurate variant calling of variants within the KIV-2 VNTR in human samples

We evaluated the performance of UMI-ONT-Seq on the KIV-2 PCR5104 fragment, which encompasses about 92% of the KIV-2 VNTR region, in 15 human validation samples with KIV-2 batch NGS sequencing data available[24]. Sensitivity, specificity, precision and F1 score were mostly very close or equal to 100% (Figure 4A; see Supplementary table 20 for single sample values and Supplementary figure 6 for R9 HAC). Most importantly, specificity (mean± SD) was 1.0±0.001 for all conditions, demonstrating a very low false-positive rate of UMI-ONT-Seq despite its relatively high raw-read error rate (Supplementary table 21). Also in human samples V14 data performed generally better than R9, while V14 SUP provided marginal benefit over V14 HAC (Figure 4A). Importantly, despite providing considerably higher raw read accuracy (median read quality ≈Q28; Supplementary table 5), UMI-ONT-Seq with duplex basecalling (SUPDUP) leads to very low sensitivity and F1 score (0.481±0.298; 0.578±0.221; Figure 4B; Supplementary table 21). This was actually expected due to the specific technical implementation of duplex basecalling and is addressed in the Discussion section.

**Figure 4:**
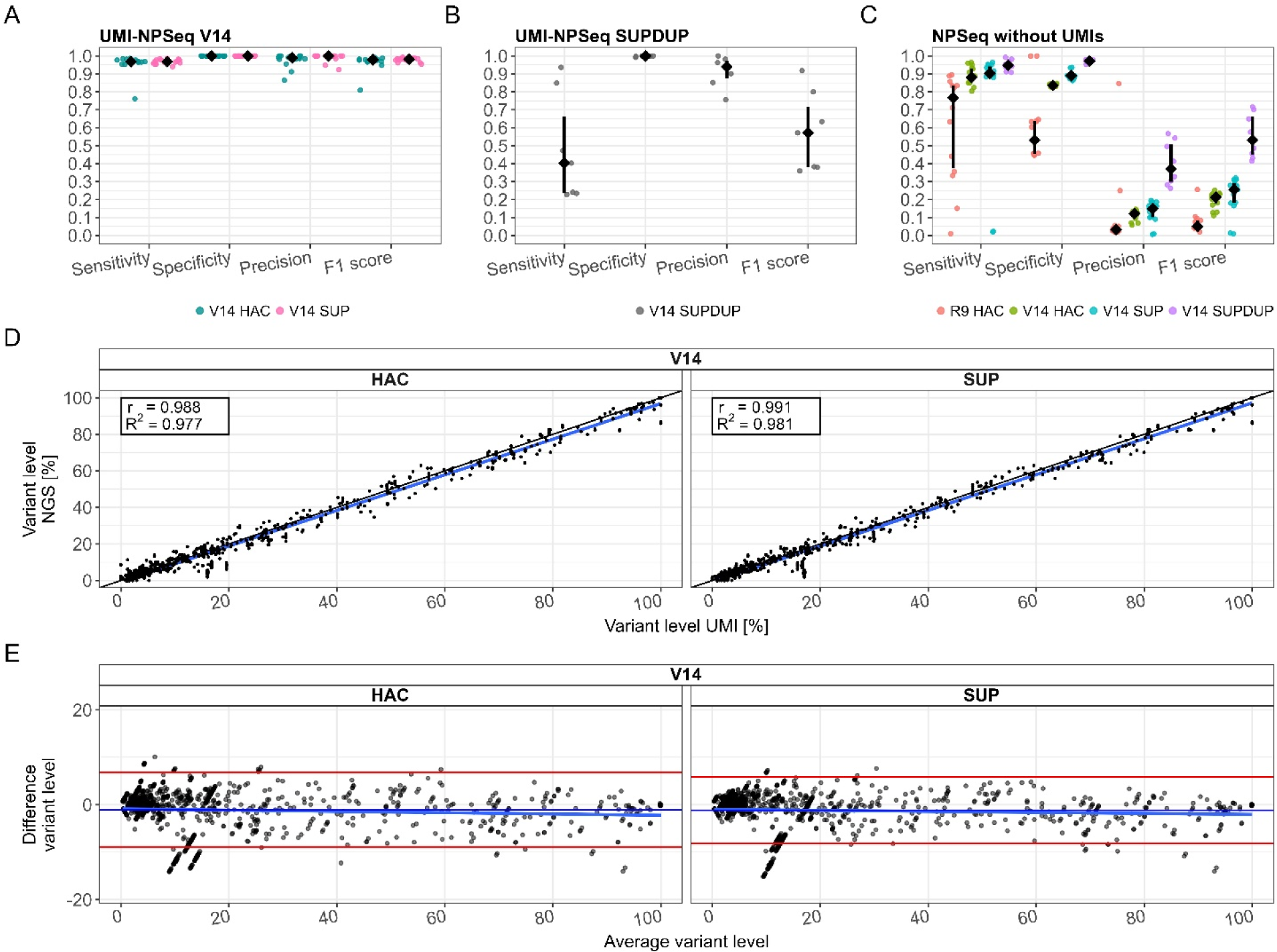
Performance measures compared to ultra-deep NGS sequencing of 15 human DNA samples (SAPHIR) for the V14 chemistry (HAC and SUP basecalling). **Panel A**: Performance of UMI-ONT-Seq for the 15 human DNA samples. Black points are median values. Colored points are the single samples. We observed high agreement between the ultra-deep NGS sequencing and UMI-ONT-Seq for most samples, leading to median performance values above 95 % and slightly higher performance values for SUP basecalling. **Panel B**: Performance values for duplex basecalling (SUPDUP). Black points and lines are median values and interquartile range. Grey points are the single samples. Despite increased raw read quality there was a significant drop in sensitivity when using SUPDUP basecalling (see Discussion for explanation). **Panel C**: Performance values for ONT-Seq (without UMIs) for different chemistries (R9, V14) and basecalling algorithms (HAC, SUP, SUPDUP) (black points and lines are median values and interquartile ranges; colored points are the single samples). Sensitivity increased with increasing raw read quality up to median values of 95 % for SUPDUP basecalling, but precision and F1 score were consistently low due to by high number of false positives. **Panel D:** Correlation of variant levels for each mutation of all 15 DNA samples (black points) of UMI-ONT-Seq for V14 HAC and SUP basecalling compared to ultra-deep NGS sequencing. We observed nearly 100 % correlation (r and R2 > 0.977) for both conditions, with no bias across the variant level range (E).

To evaluate the advantage of the UMI-ONT-Seq we called the KIV-2 variants on the same ONT-Seq data without UMI clustering (simulating a conventional KIV-2 deep sequencing approach[24]). The performance of the variant calling without UMIs increased continuously from R9 HAC to V14 HAC to V14 SUP to V14 SUPDUP (Figure 4C, Supplementary table 22), but precision and F1 score were considerably below UMI-ONT-Seq in all conditions. For V14 SUPDUP without UMIs, sensitivity and specificity reached 0.950±0.031 and 0.971±0.008, but precision and F1 score were only 0.399±0.122 and 0.554±0.123. UMI-ONT-Seq also considerably exceeded the performance of the nanopore-specific low-level variant caller ClairS-TO (Supplementary figure 8).

Importantly, UMI-ONT-Seq not only discriminated variants in the KIV-2 very efficiently, but also recapitulated very closely the individual variant levels measured previously by deep NGS[24] (R^2^: 0.977 and 0.981 for V14 HAC and SUP, Figure 4D, Supplementary figure 7). With V14 conditions, no bias was observed across the whole mutation level range (Figure 4E; Supplementary figure 9 for R9). Conversely, both the R9 chemistry and especially the UMI-free conditions showed noticeable bias and less correlation, which was exacerbated by a high number of false-negatives in the UMI-free analysis (Supplementary figure 7, Supplementary figure 9).

### UMI-ONT-Seq preserves the haplotype of each KIV-2 repeat unit and allows precise KIV-2 copy number quantification

We used 15 human DNA samples and the two unmixed plasmids of PCR5104 to develop an algorithm (for V14 SUP) to estimate the KIV-2 copy number in human genomic DNA. The number of unique haplotypes showed a high correlation with the KIV-2 copy number measured by ddPCR (r=0.84, R^2^=0.709, Supplementary figure 10A and B), but overestimated the true number of repeats (mean ± SD, 9.1±8.2, Supplementary table 23). Merging unlikely haplotypes (see Methods) reduced the deviation to only -3.6±3.9 repeats (Supplementary table 23) and increased the correlation with ddPCR considerably (r=0.96, R^2^=0.92, Supplementary figure 10C), but in turn led to an underestimation of the KIV-2 copy number, especially for the high KIV-2 numbers (Supplementary figure 10D).

We thus hypothesized that truly identical KIV-2 repeats may occur more often than assumed and may bias the KIV-2 copy number estimation (Supplementary figure 11). Indeed, in line with this hypothesis, coverage-correction of the occurrence of each haplotype finally led to nearly perfect correlation between the predicted and the expected KIV-2 copy number (r: 0.975, R^2^: 0.95) and reduced the mean deviation to only 0.4 ± 2.9 repeats (Figure 5A, Supplementary table 23). Both unmixed plasmids showed only one single haplotype and for 12 of 15 samples the estimated KIV-2 copy numbers were even within the narrow confidence interval (CI) of ddPCR (Figure 5B).

**Figure 5:**
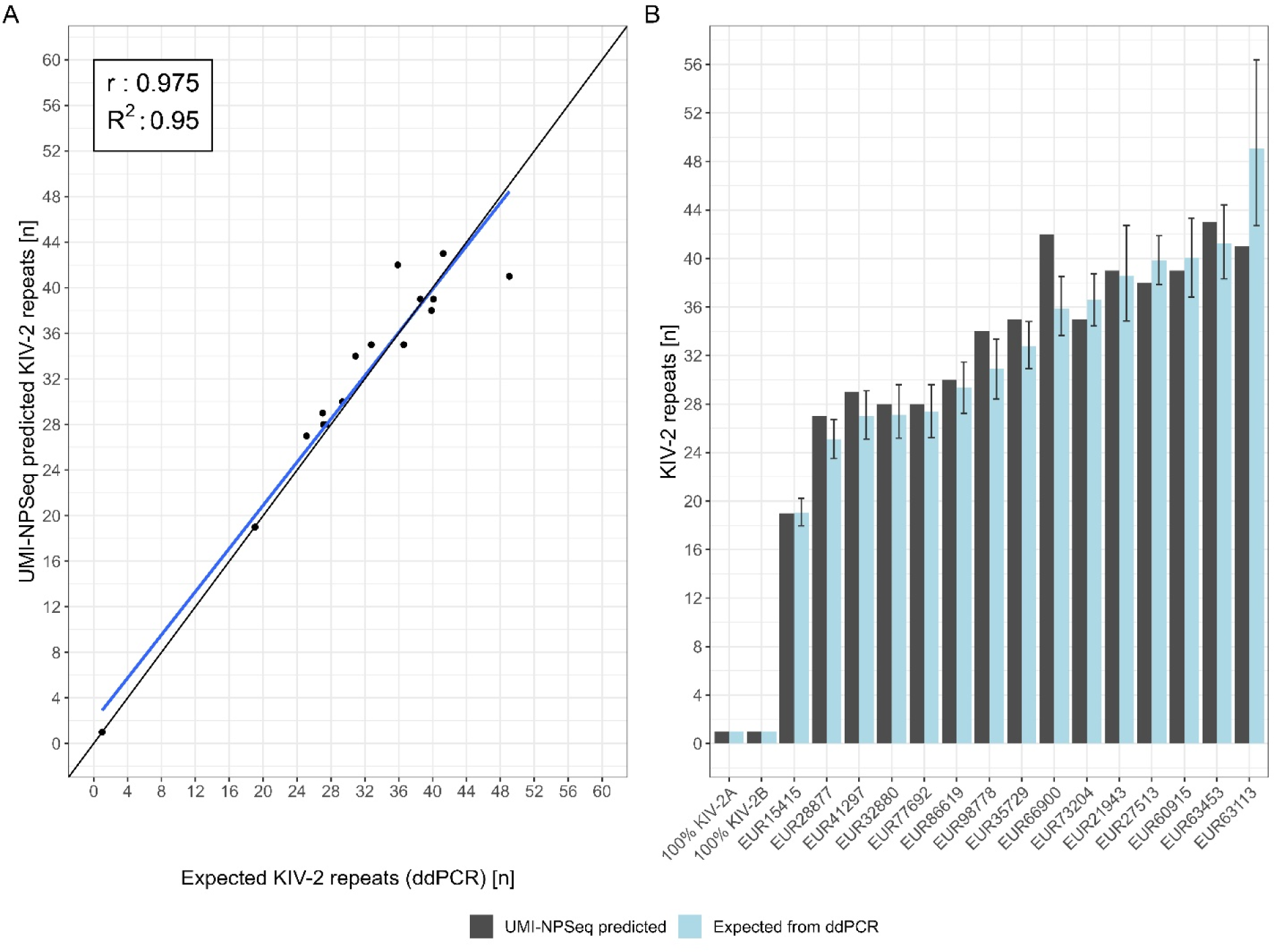
Correlation between the expected and UMI-ONT-Seq predicted number of KIV-2 repeat units. **Panel A:** Correlation of the coverage-corrected UMI-ONT-Seq haplotypes with the number of KIV-2 repeats expected from ddPCR. Coverage-corrected UMI-ONT-Seq haplotypes allow precise determination of the true number of KIV-2 repeats. **Panel B:** The barplots report the predicted versus measured number of KIV-2 repeats per sample including the confidence interval of the ddPCR quantification. 75%-80% of the samples are within the 95 % confidence interval of the ddPCR. EUR: European samples from Austria (SAPHIR study).

### KIV-2 repeat number quantification and haplotype diversity in 48 samples from 1000G

Despite the Lp(a) trait shows marked differences across ancestries, genetic Lp(a) research in the last decades was focused mainly on individuals of European ancestry. Little is known about genetic variability in non-European samples, especially within the KIV-2 region. To explore the diversity of the KIV-2 VNTR across ancestries we performed KIV-2 UMI-ONT-Seq on 48 randomly selected gDNA samples from four non-European 1000G populations (Yoruba [YRI], Dai Chinese [CDX], Japanese [JPT], Punjabi [PJL]; 12 samples per group).

The results reflect previously suggested differences between ancestries, such as the different frequencies of KIV-2B and KIV-2C units[24, 30]. Interestingly, KIV-2C repeats occurred at least once in 50 % of the East Asian samples (CDX and JPT) but were completely absent in the African samples (Supplementary table 24).

For the 35 samples that contain KIV-2B and KIV-2C repeats variant calls from UMI-ONT-Seq and NGS closely agreed (mean±SD; sensitivity: 99.5±0.7 %, specificity:100±0.1 %, precision: 98.3±2.2 %, F1 score: 98.9±1.3 %; Figure 6A; Supplementary table 25, Supplementary table 26 for per-sample values). Specificity and precision were equally high also for the 13 samples containing only KIV-2A repeat units, but sensitivity was considerably lower in these samples (59.2±4.7 %, Figure 6A and Supplementary table 25). This is caused by known issues of NGS-based variant calling in this complex region[24]. The non-repetitive KIV-3 apo(a) domain presents the same exonic sequence as KIV-2B units within the KIV-2 VNTR. When using WGS data from samples that do not contain KIV-2B units, reads from the KIV-3 domain are mistakenly aligned to KIV-2, resembling KIV-2B-specific variants and thus causing false-positive variant calls[24]. The KIV-2 specific amplification in UMI-ONT-Seq alleviates the wrong alignments. Exclusion of KIV-2B specific variants increased sensitivity for all 48 samples to 98.7±1.8 % (Supplementary table 27). As for the SAPHIR validation dataset, NGS and UMI-ONT-Seq variant levels were highly correlated also in 1000G dataset (r: 0.992, R^2^: 0.983, Figure 6B). We found no systematic bias and only deviations in the aforementioned KIV-2B specific mutations. Remarkably, KIV-2B specific mutations were detected at the same variant level, as expected from one haplotype (Supplementary figure 12).

**Figure 6:**
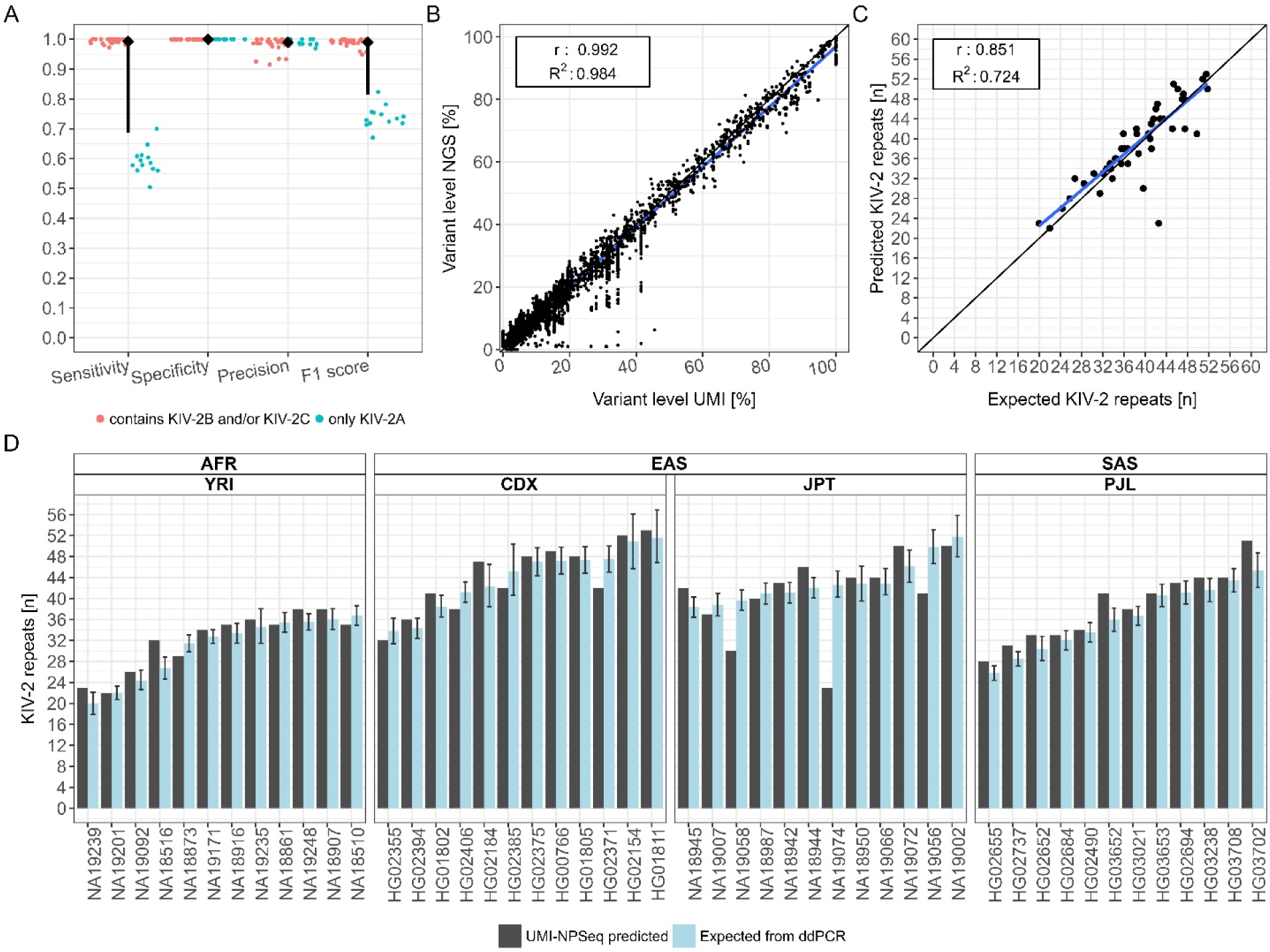
UMI-ONT-Seq analysis of 48 samples from four populations of the 1000 Genomes Project. Panel A: Performance of UMI-ONT-Seq compared to KIV-2 specific variant calling of high-coverage whole genome sequencing (WGS; ADD VNTR pipeline) for the 48 1000G samples. While sensitivity and specificity are close to perfect (mean ± SD: 0.993±0.01, 0.996±0.005), precision and respectively F1 score deviate from the WGS data. All mutations that were additionally found by the UMI-ONT-Seq were previously classified as KIV-2B specific, intronic variants, which are reported to be difficult to call in WGS data[24]. Panel B: Correlation of UMI-ONT-Seq variant levels versus WGS variant levels (n = 5968 variants). Found variant levels of both methods are highly correlated (r: 0.992, R2: 0.983). Deviations were observed only for KIV-2B specific variants. Panels C: Correlation of ddPCR measured versus UMI-ONT-Seq predicted number of KIV-2. We observed a high correlation between UMI-ONT-Seq quantified and true number of KIV-2 repeats (r: 0.851, R2: 0.724). Panel D: Comparison of UMI-ONT-Seq with ddPCR quantification. UMI-ONT-Seq accurately predicts the number of KIV-2 repeats. The mean (± SD) deviation between UMI-ONT-Seq and ddPCR was only -0.295 ± 4.26 repeats. For 32 of 48 samples even within 95 % confidence interval of ddPCR.

We observed a similar step-wise improvement for the KIV-2 repeat number estimations between the different quantification strategies as in the SAPHIR validation set (Supplementary table 28 and Supplementary figure 13). Correlation with ddPCR data for the 48 1000G samples was R^2^=0.724, which raised to R^2^=0.903 after exclusion of two outliers (Figure 6C). The mean difference was as low as 0.295 ± 4.259 repeats and 32 of 45 samples were even within the narrow CI of the ddPCR (Supplementary table 28 and Figure 6D).

In summary, UMI-ONT-Seq provides a new, very accurate, ancestry-independent method for variant calling, haplotype extraction and determination of KIV-2 repeat number for large as well as short alleles, which well exceeds the technical limit of the commonly used KIV-2 VNTR qPCR method[54–57].

### Exploration of the KIV-2 intronic short tandem repeat

Leveraging the high accuracy of UMI-ONT-Seq, we explored the poorly described CA short tandem repeat (STR) in each KIV-2 intron. This STR shows pronounced variability between the KIV-2 units and thus resembles somatic STR variation, which is a further class of hard-to-resolve genetic variation. The KIV-2 STR had been characterized before only once in just 2 individuals[58].

We extracted the STR region from the high-quality UMI consensus sequences of all 63 gDNA samples (SAPHIR and 1000G) and the two unmixed plasmids, resulting in 62,679 single STR regions. In close agreement with the prior work, we observed STR lengths between 8 to 22 repeats (12-18 in Rosby et al[58]; Supplementary figure 14; Supplementary table 29). We observed degeneration of the STR region at mainly positions 7 and 15 from to either GA or AA in 5,685 (≈9.1 %) STR sequences (Supplementary table 30). Both the degeneration patterns and the STR diversity showed potential population-specificity (Supplementary table 30 and Supplementary table 31). This provides a further example of the capabilities and broad applicability of UMI-ONT-Seq to analyze complex genetic variation.

### Haplotype diversity in 63 multi-ancestry samples

Finally, we used the extracted haplotypes to analyze the diversity in the KIV-2 repeats in 63 samples from 1000G and SAPHIR. We were readily able to classify the KIV-2 haplotypes into the commonly known KIV-2A, B and C subtypes (representative samples in Figure 7; all samples in Supplementary figure 14). We observed two novel clusters within KIV-2A, which were defined by three positions (35, 3103 and 4358; Supplementary figure 15; had been proposed previously as new haplotypes also in [24]). Interestingly, these positions were invariant in all KIV-2B and C repeats across all 63 samples. Analysis of the KIV-2B and C repeats revealed 5 positions, which define the three clusters of KIV-2B and C (positions 50, 2409, 5037, 5045 and 5052; Supplementary figure 16). Interestingly, KIV-2C repeats did not built a distinct cluster (Supplementary figure 17).

**Figure 7:**
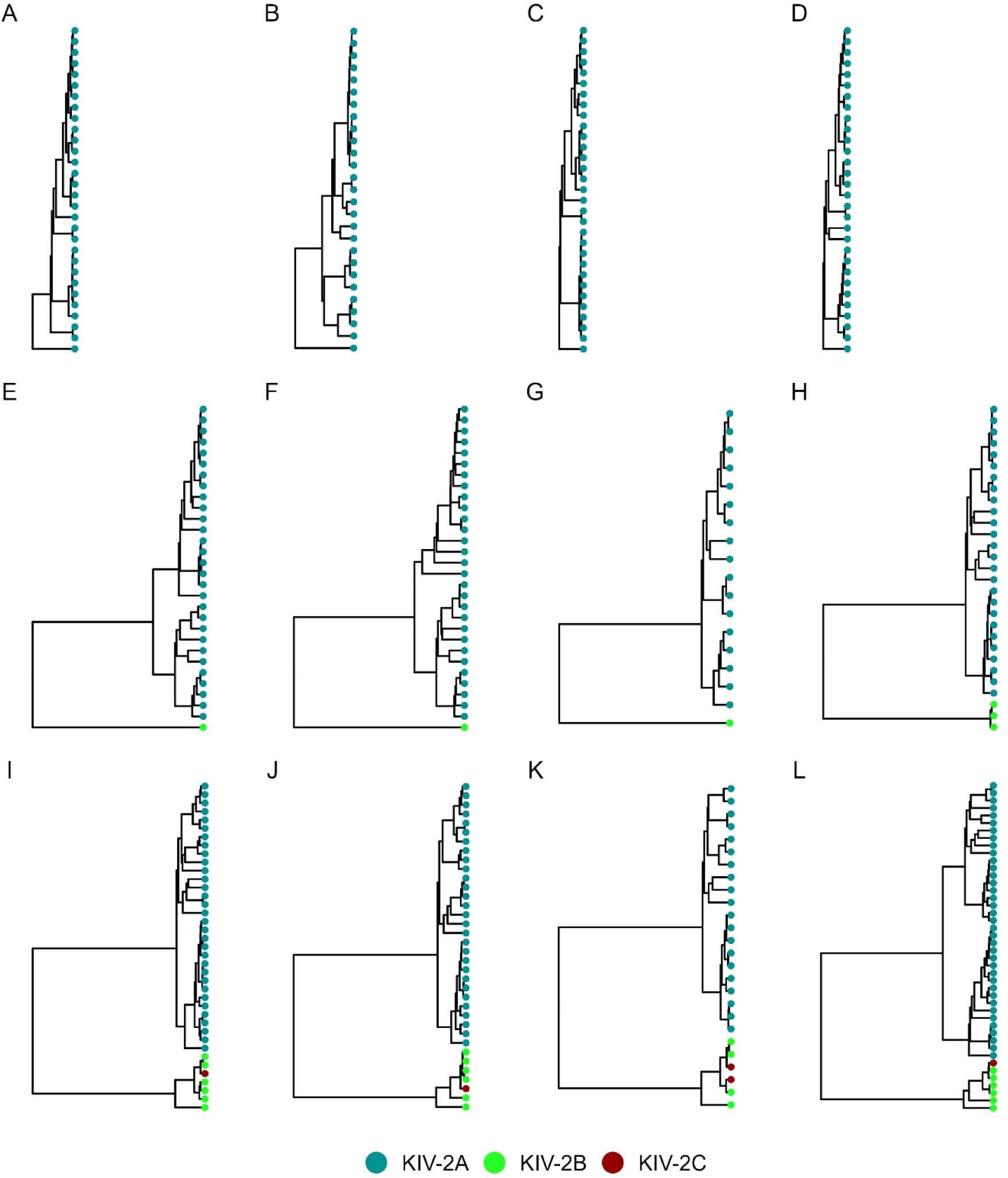
KIV-2 subtype specific haplotype diversity of 12 representative human gDNA samples in the present study (1000G, SAPHIR). Splitting the samples by their containing KIV-2 subtypes, revealed 2 major clusters containing either only KIV-2A (**panels A-D**, KIV-2A repeat haplotypes marked in blue) or the phylogenetically distant KIV-2B (**panels E-H**, KIV-2B repeat haplotypes marked in green) and C (**panels I-L**, KIV-2C repeat haplotypes marked in red) subtypes. While the KIV-2B and C cluster contain very similar sequences, with low diversity, the KIV-2A cluster has large internal differentiation (see Supplementary figure 14 for all samples).

Given the unique per-repeat resolution of UMI-ONT-Seq we finally chose to characterize in detail the haplotypes of two frequent, clinically relevant KIV-2 SNPs (4925G>A[25] and 4733G>A[26]). Either one of the two SNPs was found in 7 samples (6 EUR, 1 PJL). Both variants were exclusive to KIV-2A repeats. Remarkably, three otherwise unrelated samples showed exactly the same sequence for the complete 5.1 kb long repeat unit carrying the 4733G>A variant and two unrelated samples showed exactly the same sequence of the 4925G>A variant carrying repeat unit (Supplementary figure 18). The background haplotypes of these variants were clearly located one of the two different KIV-2A clusters (Supplementary figure 18). This highlights the potential of UMI-ONT-Seq to preserve highly accurate, full-length and per-repeat haplotype information of the KIV-2 VNTR and gaining profound insights in the per-unit haplotype structures within this complex VNTR.

## Discussion

Large, highly similar VNTR repeat units are a major constituent of the “dark genome”[2]. Nanopore sequencing has recently achieved well beyond 99% variant calling accuracy for germline mutations[59, 60], but identification and direct phasing of differences in highly homologous regions (especially VNTRs) remains challenging. The medically highly relevant *LPA* KIV-2 VNTR[17] is among the most complex coding VNTRs in the human genome[15, 27] and still poses issues in VNTR variant calling[1, 11, 61], making it a challenging and interesting model. We extensively evaluated the performance of amplicon-based UMI-ONT-Seq for SNP detection, direct SNP phasing and determination of VNTR repeat numbers.

In all experiments, the UMI-ONT-Seq with V14 chemistry showed nearly perfect variant calling performance, extraordinary correlation with NGS data and no systematic bias across the variant level range. The effects of raw read quality on variant calling were marginal between V14 HAC and V14 SUP and affected only a few positions in the PCR5104 amplicon (Supplementary table 10). UMI-based variant calling clearly outperformed UMI-free KIV-2 variant calling, which suffered from low precision and low sensitivity, depending on the variant caller used.

The low performance of using duplex called UMI reads despite nearly Q30 raw read quality may appear counterintuitive, but is intrinsic to the duplex calling process. Duplex basecalling performs consensus basecalling on all strands that enter the same pore within 20 seconds, show similar length and have ≥60% homology between the last, respectively first 250 bp. With amplicons showing >90% similarity like the KIV-2 amplicons[24] this leads to extensive false strand pairings.

Using the V14 chemistry, dataset-quality achieved ≈Q40 already with 6-8 reads per UMI cluster, providing highly accurate sequences of each KIV-2 repeat unit. We identified a V14-specific error profile, which was systematic and limited the consensus quality. Indeed when disregarding these errors, 95.5 % of all UMI-ONT-Seq consensus sequences were error-free at cluster size 6 to 8. These characteristic transversion errors might represent a current limitation of UMI-ONT-Seq, especially for very low-level (≤0.5 %) mutation detection. However, even without error profile correction, one third of the consensus sequences were error-free and 99.7 % of the sequences had no more than three errors. Noteworthy, the observation that the residual errors are systematic makes them well addressable bioinformatically and/or by modified Medaka training sets.

Overall, the performance of V14 was significantly better than previous reports using R9 chemistry across all experiments. Karst et al reported that 15 to 25 reads per UMI cluster were required to achieve Q40 dataset error rates with R9 chemistry[34], which is in agreement with our observations (Supplementary figure 4). Conversely, the low minimal cluster size required by V14 for Q40 quality now puts UMI-ONT-Seq close to the performance of UMI-tagged Pacific Bioscience circular consensus sequencing (PacBio CCS), which requires three reads per UMI cluster for Q40[34] but comes with 100 times higher equipment investments compared to ONT MinION systems. Of note, PacBio CCS data without UMIs showed chimeras up to 3% mutation level, thus requiring UMIs for VNTR sequencing also with PacBio data despite the higher intrinsic quality of PacBio CCS. Given the relatively low cluster size required, UMI-ONT-Seq allows multiplexing ≈50 gDNA samples per MinION flow cell, with costs of roughly 25€/sample on MinION systems and likely <10 €/sample on PromethION systems, which corresponds to ≈0.06/0.025 cents per consensus sequence (assuming roughly 40,000 consensus sequences on average in our setup).

All observations confidently supported that UMI-ONT-Seq provides nearly error-free consensus sequences that allow direct experimental haplotyping of KIV-2 units down to ≤1 % fractional representation. The high consensus sequence quality allowed direct extraction of the repeat structure of an STR located in the intron of the KIV-2 VNTR, which is a challenging task as it resembles somatic STR mosaicism. Given the growing interest in somatic STR mosaicism[62], UMI-ONT-Seq may provide a new efficient way to generate high-quality reference data. We also employed UMI-ONT-Seq to characterize phylogenetic subclusters and two frequent disease-relevant SNPs hidden in the KIV-2 VNTR (KIV-2 4925G>A and 4733G>A). We could readily define subcluster-specific mutations and observed high sequence homology (up to 100 %) of 4925G>A and 4733G>A carrying repeats (possibly indicating either rather recent mutation events or a conservation of these Lp(a)-reducing haplotypes). UMI-ONT-Seq was readily able to resolve the full–length haplotype of these variants, revealing exclusive location with two different KIV-2A subclusters, which confirms previous suggestions that these two most strongly Lp(a)-lowering variants are indeed located in trans on different Lp(a) alleles[26]. Given the high per-repeat resolution of UMI-ONT-Seq we hypothesized that the genomic VNTR repeat number could be accurately determined by coverage-corrected quantification of the unique haplotypes, because truly identical KIV-2 repeats that would arise from e.g. recent expansions would be detectable by higher normalized coverage. We show in 63 samples that UMI-ONT-Seq indeed well exceeds the technical limit of the largely used[63] KIV-2 qPCR assays and provides repeat number estimates that are comparable to high resolution ddPCR, which is among the most precise technologies for copy number quantification and is able resolve ≈1.1-1.2-fold differences (compared to ≈2-fold in qPCR)[64, 65].

Of note, a recent preprint by Behera et al reports a commercially available algorithm to determine the KIV-2 copy number using NGS data[11]. The authors determined the KIV-2 copy number of all 1000G samples and were able to phase the copy number to the two LPA alleles in 47% of the samples, providing the allelic KIV-2 copy number. The KIV-2 copy number reported by Behera et al matches accurately our ddPCR data and, more importantly, the KIV-2 copy number derived by UMI-ONT-Seq matches accurately the data of Behera et al. in 46 of 48 samples (R^2^: 0.965 for these 46 samples, Supplementary table 32). Conversely, current NGS-based approaches do not provide full-length sequences and haplotypes of each KIV-2 unit. The two approaches are thus complementary and can be used together to investigate the KIV-2 genetics at scale.

In general, we observed a considerable improvement of copy number prediction after correcting for haplotype coverage. This was most pronounced in the larger alleles, which may suggest that especially some larger KIV-2 alleles consist of multiple identical units. These may have originated from relatively recent repeat expansion and only slow divergence of the KIV-2 VNTR. Very little is known about the frequency and mechanisms of KIV-2 expansions[23]. Boerwinkle et al report generation of one new allele in 376 meioses[66], but no other reports are available. UMI-ONT-Seq allows for the first time to study the mutational and evolutionary mechanisms of this complex VNTR with single nucleotide, respectively single haplotype resolution at scale.

Given its direct portability to other VNTRs with similar structure, UMI-ONT-Seq provides a novel instrument with general applicability beyond *LPA.* It complements and expands similar approaches like circularization-based concatemeric consensus sequencing (R2C2)[67] or linked-read sequencing. R2C2 with UMIs recently reported up to Q50 quality for 550 and 1200 bp long amplicons using cluster sizes 12-17[68], but might be limited to smaller amplicons, as the coverage on concatemerized targets is also a function of the target length. Conversely, linked-reads can present technical difficulties when analyzing highly similar amplicons[33]. Further use cases for UMI-ONT-Seq may include mapping of epistatic protein mutations in in-vitro evolution experiments and deep mutational scans, monitoring of intra-host disease evolution, immune repertoire mapping, mapping of large inserts for massive parallel reporter assays, generation of reference sequences for complex regions and any other applications requiring precise long consensus sequences and/or SNP phasing at clonal resolution down to 1 %[33, 61, 69, 70].

## Conclusion

Using the *LPA* KIV-2 VNTR as a model of a highly complex, challenging and medically-relevant VNTR, we demonstrate the capability of amplicon-based UMI-ONT-Seq to accurately detect mutations, determine the full-length SNP haplotype of each VNTR unit and determine the VNTR copy number using coverage-corrected haplotypes in recombinant standards, human validation samples and multi-ancestry samples from 1000G. This provides a new, straightforward approach to map variation in such challenging regions. The use of an amplicon-based approach circumvents costly and laborious high molecular weight DNA WGS and provides an efficient method to generate reference data for complex regions with clonal resolution at scale.

## List of abbreviations

CNV: Copy number variation
VNTR: Variable number tandem repeat
STR: Short tandem repeat
Apo(a): Apolipoprotein(a)
Lp(a): Lipoprotein(a)
KIV: kringle IV
KIV-2: KIV type-2
HMW: High molecular weight
LMW: Low molecular weight
LD: Linkage disequilibrium
ONT: Oxford Nanopore Technologies
UMI: Unique molecular identifiers
HAC: High accuracy
SUP: Super accuracy
SUPDUP: Super accuracy with duplex
NGS: Next generation sequencing
UMI-ONT-Seq: Nanopore sequencing with unique molecular identifiers
SAPHIR: Salzburg Atherosclerosis Prevention Program in subjects at High Individual Risk
1000G: 1000 Genome Project
ddPCR: Digital droplet PCR
CI: Confidence interval
LSP: Locus specific primer
UVP: Universal primer
WGS: Whole-genome sequencing
YRI: Yoruba
CDX: Day Chinese
JPT: Japanese
PJL: Punjabi
EUR: European

## Resources

The UMI-ONT-Seq analysis pipeline is available at https://github.com/genepi/umi-pipeline-nf. All code used for data analysis is available at https://github.com/AmstlerStephan/UMI-ONT-Seq_Analysis and https://github.com/AmstlerStephan/haplotyping-KIV2-nf.

## Author contributions

SA performed the experiments, conceived and implemented the modifications to the original ONT pipeline, conceived and implemented the haplotype extraction procedure, analyzed and interpreted all data and wrote the manuscript. GS established the original ONT UMI amplification protocol. CP generated all ddPCR data and supported the project. LF, HW and SS provided computational support and scientific input. SS created the data analysis tools for KIV-2 variant calling in WGS NGS data and generated the 1000G NGS reference data. SDM contributed to development of the KIV-2 variant calling in WGS NGS data and generated the 1000G NGS reference data. BP is responsible for the conduct of the SAPHIR study. FK provided resources, performed Western blot analyses of SAPHIR and gave scientific input to the project. SC conceived the project, reviewed all data, acquired funding, wrote and reviewed the manuscript and supervised the work. All authors read and approved the final manuscript.

