## Supplementary materials for "Nanopore sequencing with unique molecular identifiers enables accurate mutation analysis and haplotyping in the complex Lipoprotein(a) KIV-2 VNTR"

**This PDF file includes:**

Supplementary methods
Supplementary figures 1 - 18
Supplementary tables 1 - 32

### Supplementary methods

#### Human samples for validation

Samples were drawn from the healthy-working population SAPHIR (Salzburg Atherosclerosis Prevention Program in subjects at High Individual Risk)[1]. SAPHIR is an observational study conducted in the years 1999-2002 involving healthy unrelated subjects (645 females from age 39 to 67 and 1093 males from age 39 to 66). Study participants were recruited by health screening programs in companies in area of Salzburg, Austria. For KIV-2 quantification by ddPCR 40 ng template DNA was digested with 2 u EcoRI for 1 h at 37°C and genomic KIV-2 number was assessed on a Biorad QX-200 ddPCR system using the KIV-2 quantification primer/probe set from Lanktree et al[2] (Supplementary table 4). Three ddPCR replicates per sample were merged for precise KIV-2 counting[3].

#### Extraction of the KIV-2 intronic CA STR

STR sequences were extracted from the consensus sequences using a custom python script (available at https://github.com/AmstlerStephan/UMI-NPSeq_Analysis). The STR region was defined as reference sequence position 2472-2506. After extraction of this region the actual start and end of the STR was defined by the first and last occurrence of a CA repeat. Like the process of extracting haplotypes, we employed a sample-specific threshold to eliminate highly improbable STR sequences. We determined this threshold based on a binomial distribution with a 0% likelihood for an STR sequence to occur. Any STR sequences occurring below this threshold were excluded from the analysis.

#### Detailed PCR Protocols

To create the UMI-tagged amplicon library, we adapted and optimized an existing protocol, which is provided by ONT (protocol ID: cpu_9107_v109_reve_09oct2020).

Samples were diluted to contain approximately 50,000 target molecules in 4 µl (1 KIV 2 repeat ≙ 1 target), under maximum case scenario of 80 KIV 2 repeats for the genomic DNA (gDNA) sample (up to 40 copies per allele [15]) s. Average molecular weight of one base-pair was assumed to be 660 g/mol for double stranded DNA. Genomic DNA samples were normalized to a concentration of 20 ng/µL and then diluted to 50,000 targets by mixing them 1:36 with nuclease free water. Plasmid mixtures, with a concentration of 6.5 ng/µL were diluted to 50,000 targets with a dilution series in four steps (1:16 per step, 1:65536 in total) to have sufficient volume for accurate pipetting. The PCR to generate the amplicon library is separated in three parts, which are a tagging PCR followed by two amplification PCRs ("Early PCR" and "Late PCR"). A detailed PCR protocol can be found in Supplementary table 2.

After tagging, the UMI-primers and untagged molecules are digested using ExoCip (New England Biolabs [NEB]; Ipswich, MA, USA; E1050L), mixing 3 µL ExoCip A, 3 µL ExoCip B and 15 µL of the PCR product. The protocol was run at 4 min digestion (37°C) and 4 min inactivation (80°C).

After the "Early-PCR" the amplicons were purified by adding 45 µL AMPure XP beads (Beckman Coulter; Indianapolis, Indiana USA; 19836400) and 10 min incubation on a Hula mixer. The beads were pelleted on a magnet and washed twice with 200 µL 70 % ethanol. After spinning down, re-pelleting and removal of the residual ethanol, the DNA was eluted in 21 µL nuclease free water with 5 min incubation.

After the "Late PCR" the amplicons were again purified with AMPure XP beads as described before but eluted in 30 µL nuclease free water.

#### Quality control of the UMI-amplicons

To ensure the quality and target specificity of the UMI amplicons the concentration of the all samples was measured using a Qubit 3 (Thermo Scientific; Waltham, MA USA; Q33238) fluorometer with the DNA high sensitivity kit (Thermo Scientific; Waltham, MA USA; Q33231). Additionally, 3 µL of the PCR product of the "Late PCR" (prior the final clean up) were loaded onto an 1% agarose gel. As size reference an 1.5 µL of GeneRuler 1 kb ladder (Thermo Scientific; Waltham, MA USA; SM0311) was used.

#### Ligation sequencing amplicons – Native Barcoding Kit V14 and R9

The ONT protocols for both chemistries were followed (R9: SQK-LSK109, EXP-NBD104 and EXP NBD114, protocol version: NBA_9093_v109_revK_12Nov2019; V14: SQK NBD114.24, protocol version: NBA_9168_v114_revD_15Sep2022). We used 250 fmol for V14 and 200 fmol for R9 as input for the end prep of the DNA. After library preparation the final sequencing mix that is loaded onto the flow cell should contain was adjusted to contain 20 fmol of the library.

### Supplementary figures

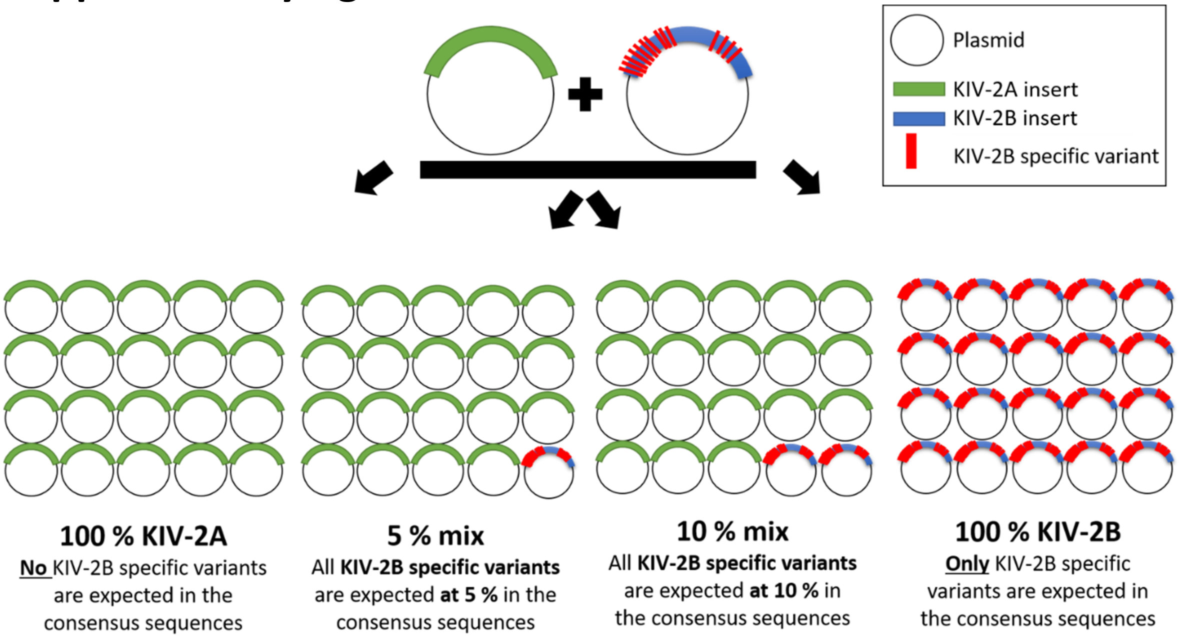

**Supplementary figure 1 Principle of the plasmid mixtures containing KIV-2A and KIV-2B inserts in different ratios that were used for benchmarking UMI-ONT-Seq.**

The plasmids were created using KIV-2 specific amplicons spanning either the inter-kringle intron (2645 insert; amplicon PCR2645) or the intra-kringle intron (5104 bp insert; amplicon PCR5104) of either KIV-2 subtype A (KIV-2A, green) or KIV-2 subtype B (KIV-2B, blue). Generation of the plasmids has been detailed in [4]. KIV-2A and KIV-2B plasmids were mixed at different ratios (100:0 to 95:5 and 0:100), with the KIV-2B comprising the minor fraction to resemble the in-vivo situation. Each KIV-2B insert differs from the reference at 85 (PCR5104) and 116 (PCR2645) positions (red lines). These differences are present all on the same molecule, representing thus one haplotype. Therefore, all KIV-2B positions shall occur at the same level in the consensus sequences. The figure exemplifies mixtures for unmixed (100 %) KIV-2A, unmixed (100%) KIV-2B, 5 % KIV-2B mixture and 10 % KIV-2B mixture.

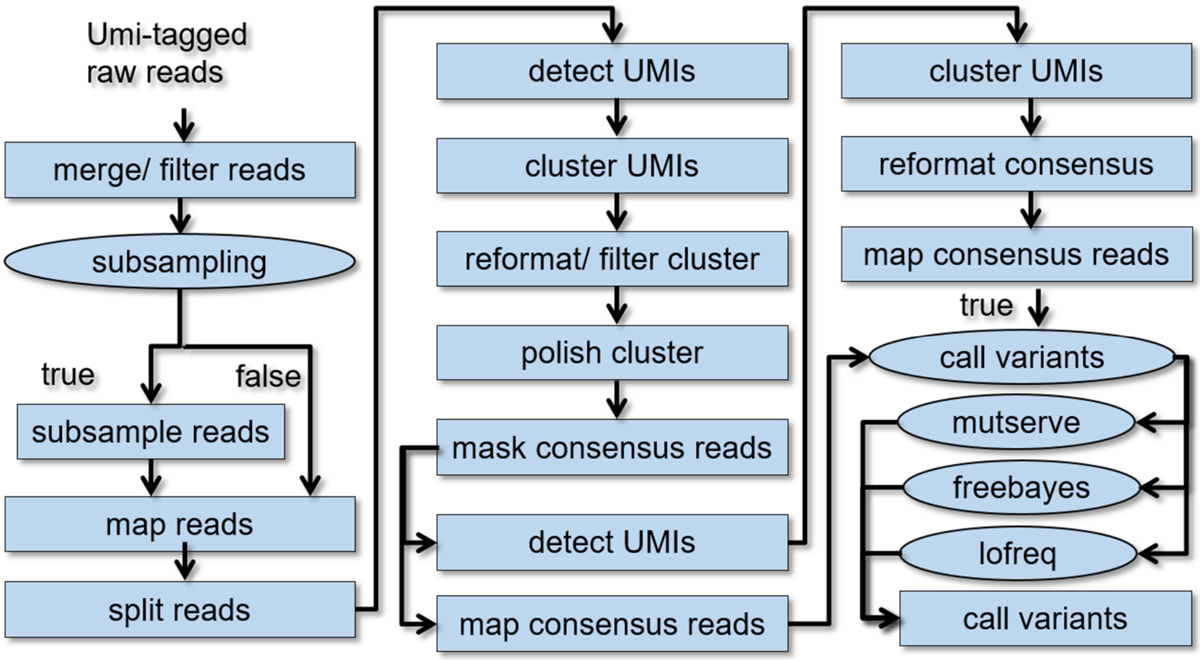

**Supplementary figure 2 Flow chart of the UMI-ONT-Seq analysis pipeline.**

The analysis workflow 1. merges and filters all FASTQ files per sample, 2. Performs an optional read subsampling step, 3. Maps the reads against the reference sequences, 4. Splits the aligned reads to keep only full-length reads with UMI-tags on both sides, 5. Extracts the UMI-sequences in the reads, 6. Clusters them according to the UMI-sequences, 7. Filters the UMI-clusters to keep only reads above a minimal cluster size threshold and 8. polishes each cluster of UMI-sequences to obtain one consensus sequence per cluster. Detection (9), clustering and consensus read creation is repeated and afterwards the consensus sequences are mapped against the reference for variant calling with mutserve[5] (alternatively also Freebayes[6] (v1.3.2) or Lofreq[7] (v2.1.5) can be invoked).

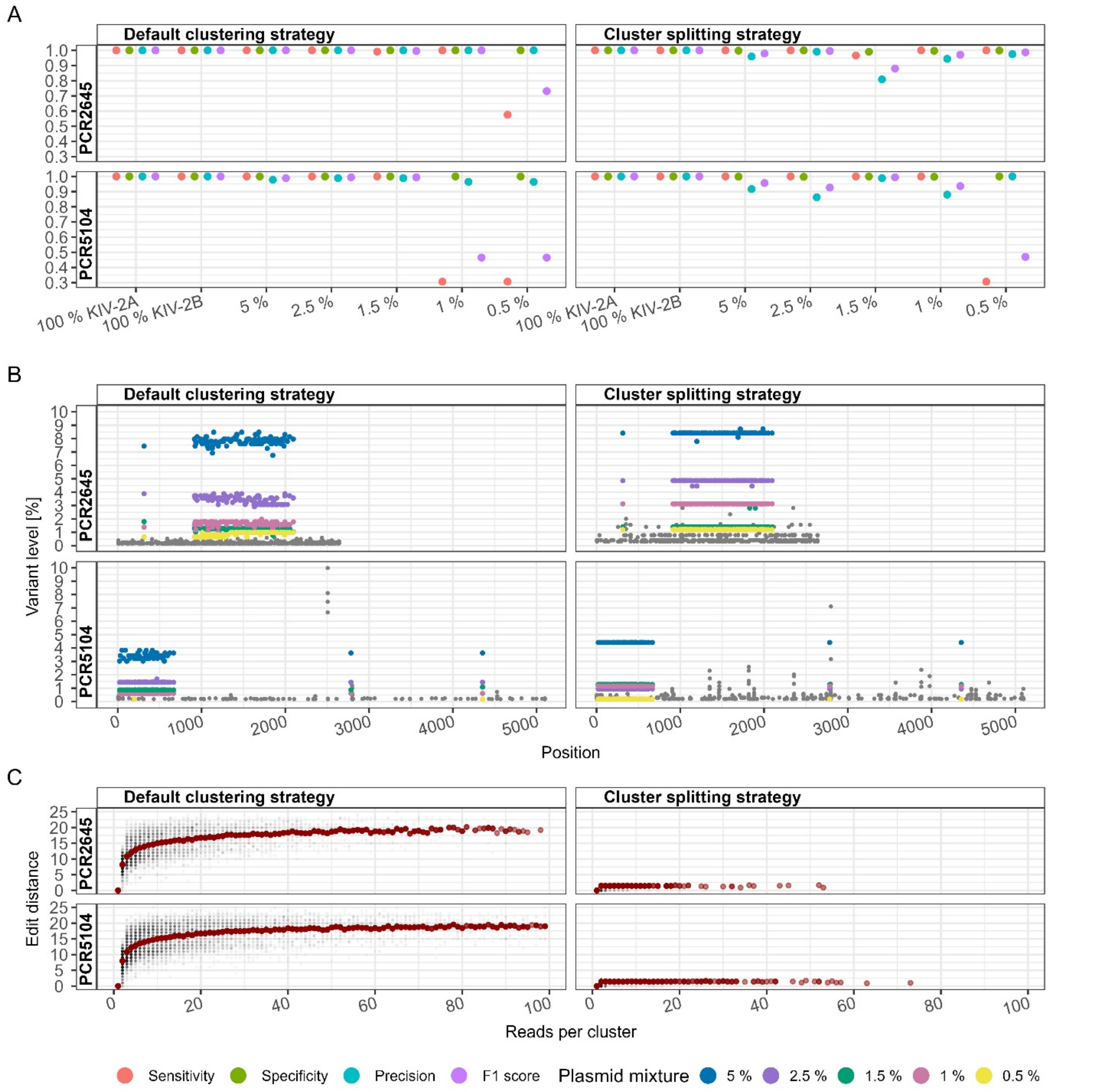

**Supplementary figure 3 Variant detection in plasmid mixtures with the UMI-ONT-Seq analysis pipeline using default clustering (left column) or cluster splitting strategy (right column) for the R9 chemistry and HAC basecalling.**

**Panel A:** Variant recall performance measures for the UMI-ONT-Seq analysis pipeline for different plasmid mixtures (KIV-2B in KIV-2A background, 0.5 – 100 %) and both amplicons (PCR2645, PCR5104). Down to 0.5 % percent variant level performance values were close to 100 % for all conditions. **Panel B:** detected variant levels of the UMI-ONT-Seq analysis pipeline using default clustering or cluster splitting strategy for the plasmid mixtures from 0.5 – 5 % across every position of both fragments. Low-level residual noise (grey points) led to a lower detection limit of ~0.85 % variant level. Variance of detected variant levels of up to ±1 % (blue points, 5 % mixture) was observed for the default clustering strategy. **Panel C** shows the edit distance of the UMI-sequences for different UMI cluster sizes for two approaches. Using default clustering strategy, UMI-sequences with an edit distance up to 25 were found in a single cluster. Using the cluster splitting strategy reduced the maximal edit distance per cluster to ≤ 2.

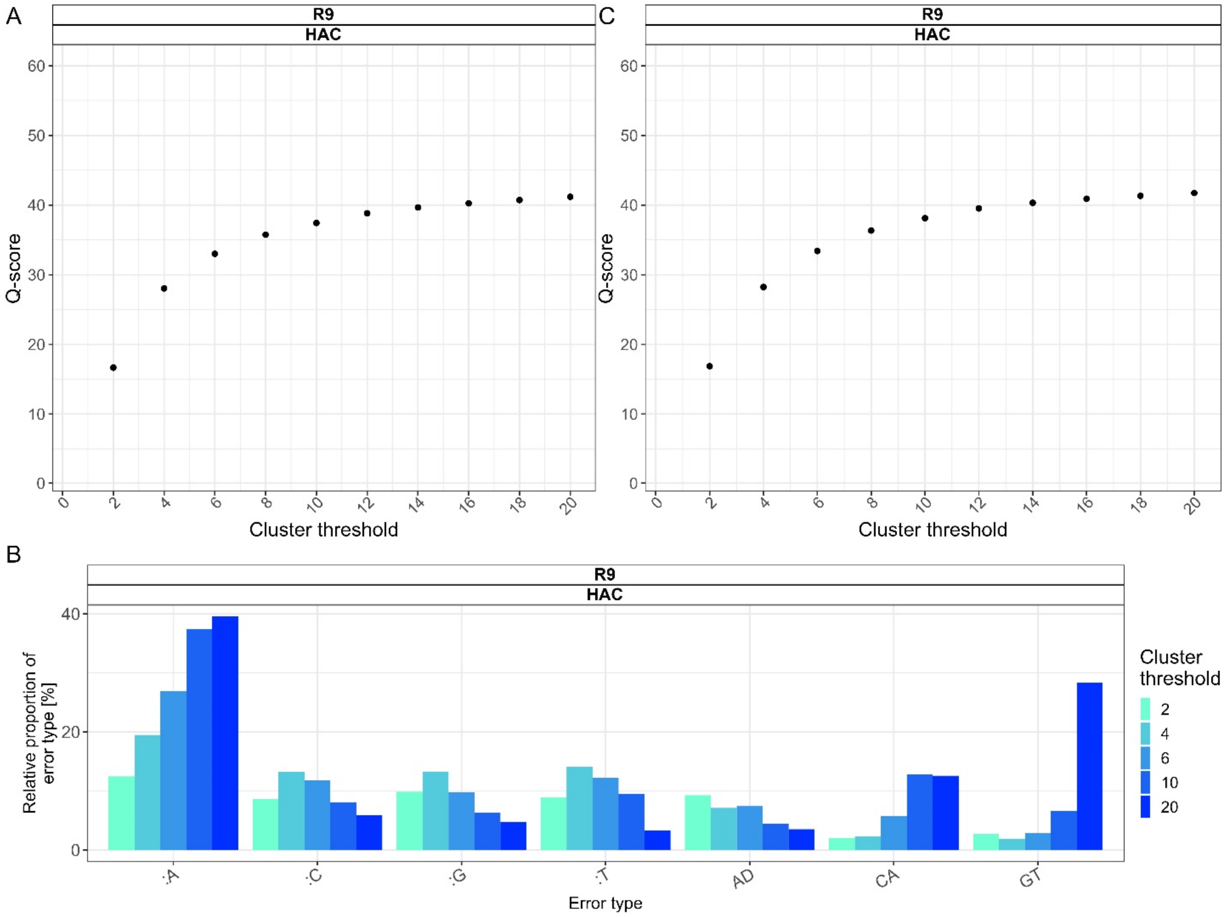

**Supplementary figure 4 Dataset Q-score estimation using KIV-2A and KIV-2B containing plasmids for the R9 chemistry and HAC basecalling algorithm.**

**A)** Dataset Q-score of the consensus sequences. **B)** Characteristic errors for the R9 HAC condition. Like ONT-Seq without UMIs, insertion are the main characteristic error (40%) also in R9 UMI-ONT-Seq at all cluster size thresholds. Panel **C)** Dataset Q-score per cluster size threshold after correction for low-level transversions (G to T and C to A). Correction for these errors caused marginal increase of the dataset Q-Score.

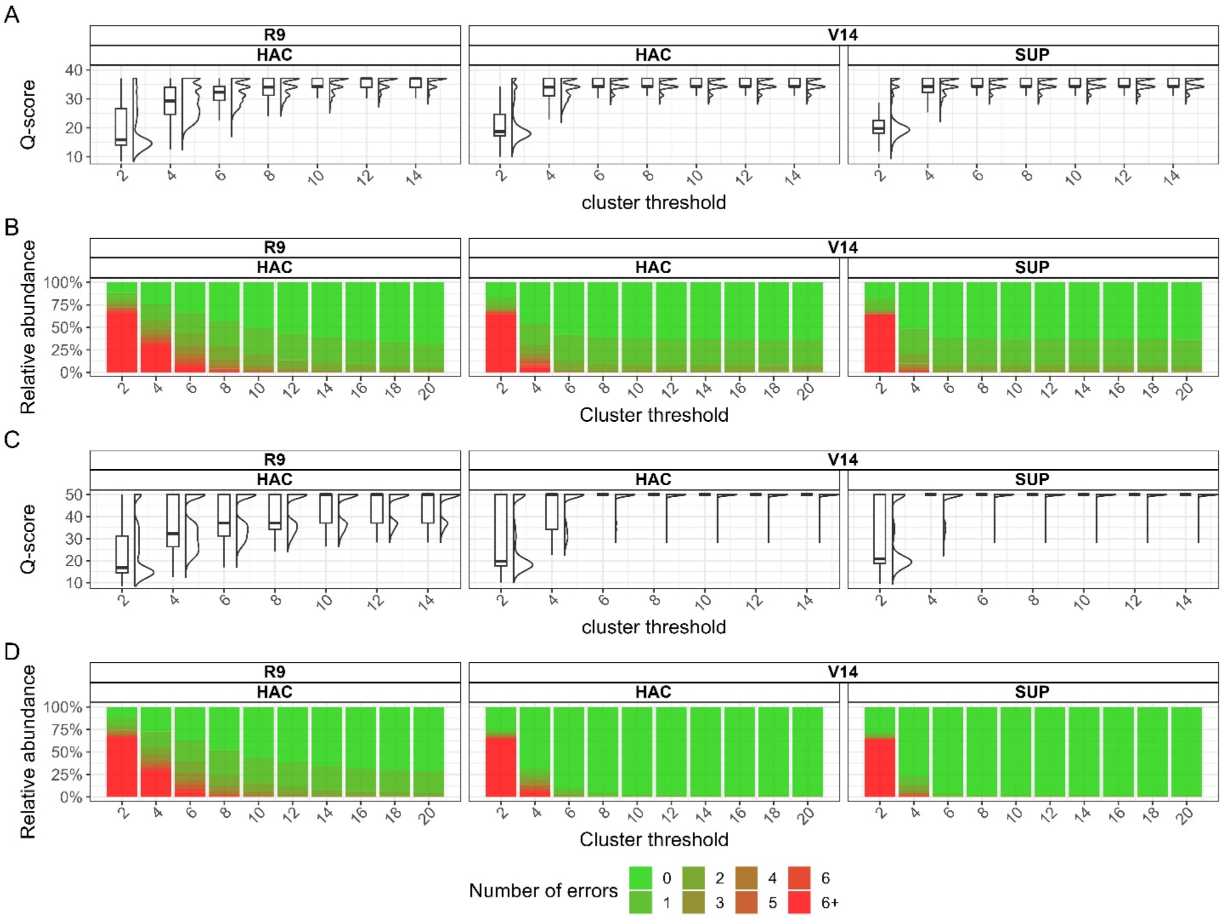

**Supplementary figure 5 Distribution of the consensus sequence Q-Score depending on the minimal cluster threshold for all sequencing conditions of the 4 unmixed KIV-2A or KIV-2B plasmids (both PCR2645 and PCR5104).**

**Panel A:** Consensus sequence Q-score for each chemistry and sequencing condition depending on the minimal cluster size threshold chosen for the analysis. While for the R9 chemistry the median Q-score increased steadily up to a minimal cluster size of 12, the V14 chemistries median Q-score increased rapidly, reaching the maximal per-read Q-score already at 6 reads per cluster (limited by the fragment length). Noteworthy, panel A includes only imperfect reads, since perfect reads would have an infinite Q-score. Therefore, **panel B** shows the percentage of perfect reads depending on the minimal cluster size threshold. The same trend as for the median Q-score is observed also here. The percentage of perfect reads increases steadily. At 12 reads per cluster 48 %, 58 % and 62.5 % of all consensus sequences were perfect for R9 HAC, V14 HAC and V14 SUP respectively.

Correcting for characteristic errors (GT and CA conversions, see Results for details), increased median Q-score **(C)** and percentage of perfect reads **(D)** per minimal cluster threshold drastically. At 6 reads per cluster almost 100 % of the reads of both V14 conditions were perfect.

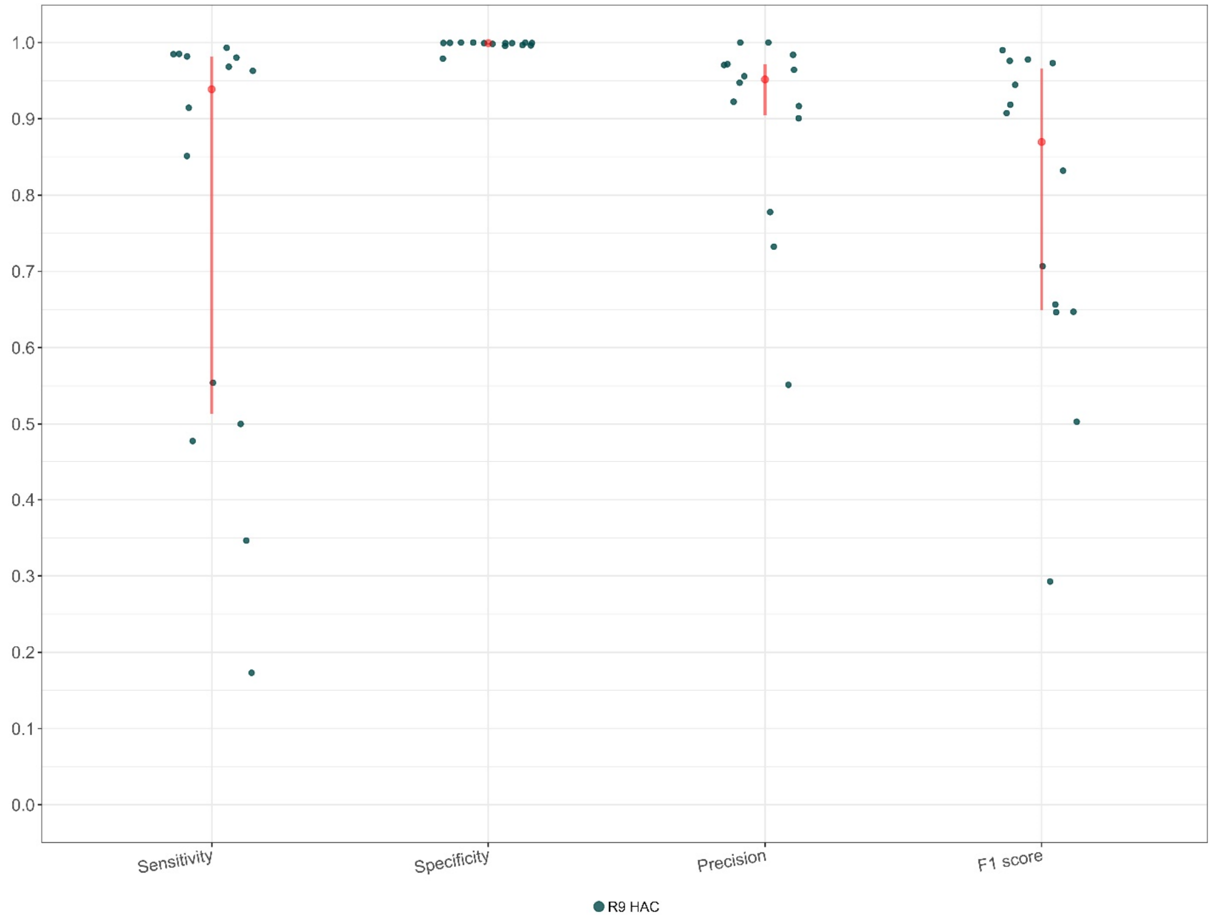

**Supplementary figure 6 Performance measures for the R9 chemistry and HAC basecalling algorithm for 15 human gDNA samples (SAPHIR).**

The red point and line show median values and interquartile range. The blue points are the single samples. Despite having some outliers, the median sensitivity is above 90 %, specificity 100 %, precision above 95 % and F1 score above 85 %.

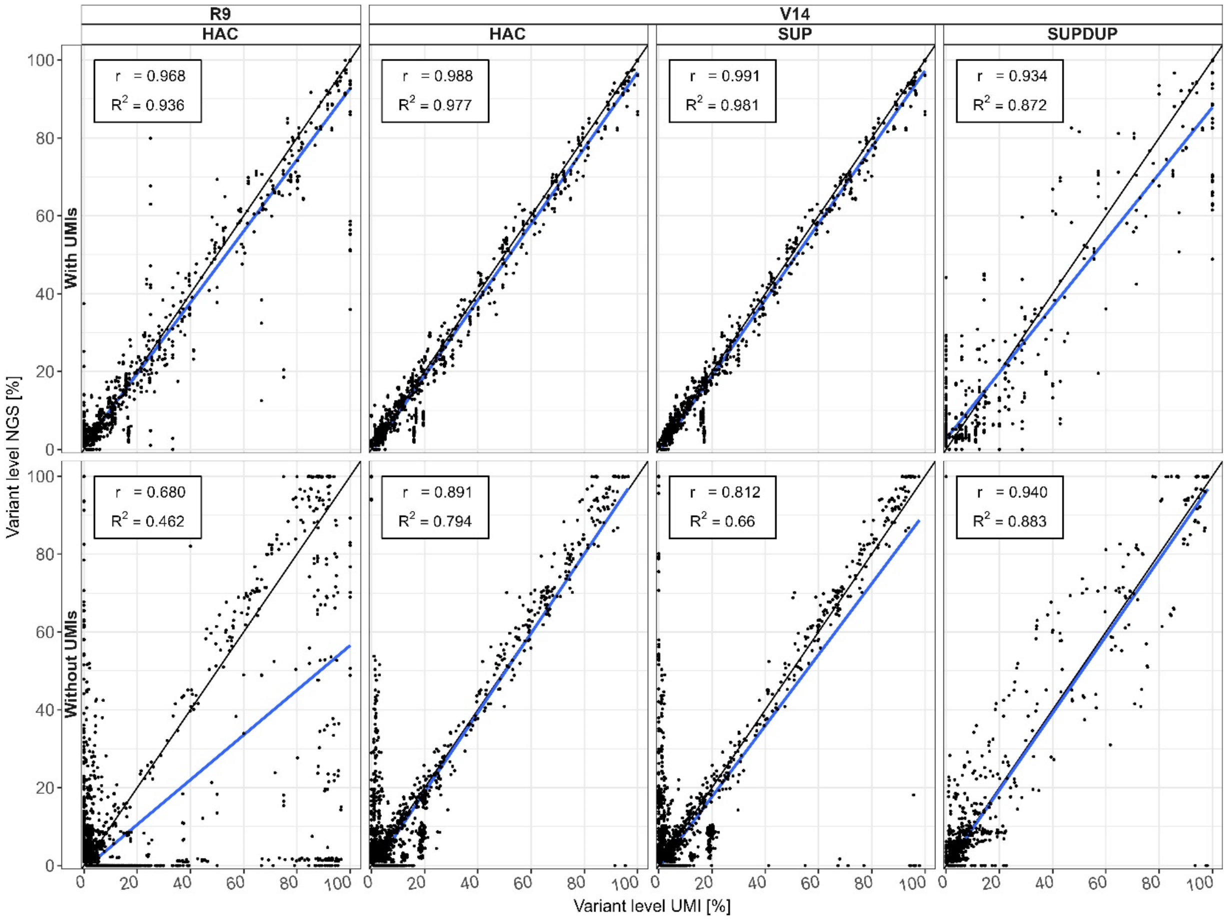

**Supplementary figure 7 Correlation of the mutation levels in standard UMI-free ONT-Seq and in UMI-ONT-Seq with levels determined by deep KIV-2 NGS[4] for all sequencing chemistries and basecalling algorithms.**

Black points are variants found in NGS, ONT-Seq (top panels) or UMI-ONT-Seq (bottom panels). Without UMIs (using only the raw reads) we observed increasing correlation with increasing raw read accuracy (lowest to best: R9 HAC, V14 HAC, V14 SUP, V14 SUPDUP). For UMI-ONT-Seq we had generally high correlation between the variant levels compared to NGS for all conditions. V14 HAC and V14 SUP showed the best correlation ≥97 %, while SUPDUP showed lower performance due to the reasons described in the main manuscript.

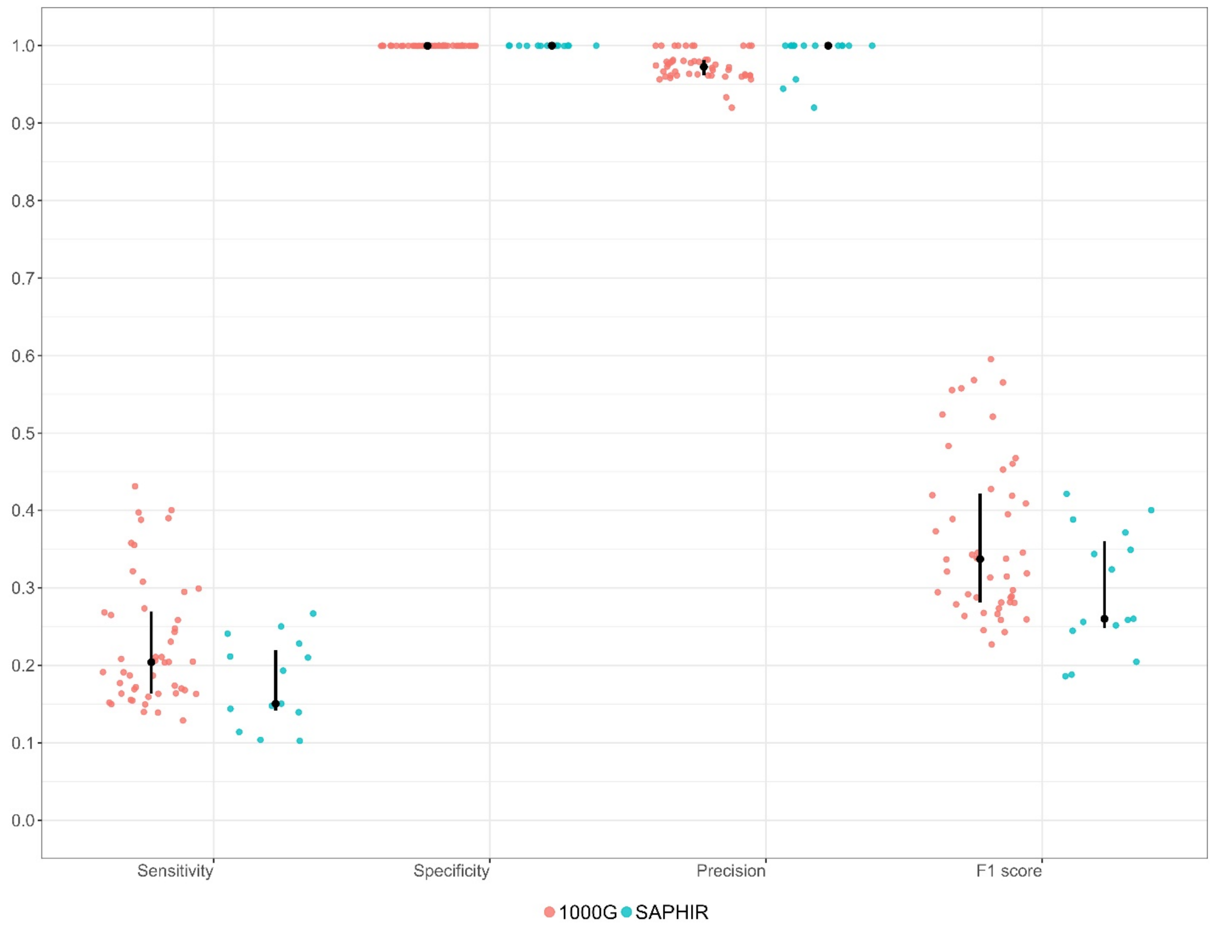

**Supplementary figure 8 Performance of the nanopore-specific low-level variant caller ClairS-TO for 63 genomic DNA samples (15 samples from SAPHIR, 48 samples of 4 populations from 1000G) compared to conventional KIV-2 NGS[4] (SAPHIR, turquoise) and variants derived from publically available WGS data[4, 8] (1000G, red).**

ClairS-TO detected only a fraction of the actual variants of the genomic DNA samples compared, resulting in low median sensitivity (≈21 % for 1000G and ≈15% for SAPHIR). Specificity and precision are high for both sample sets (>95 %), while the low F1 score reflects the low sensitivity values.

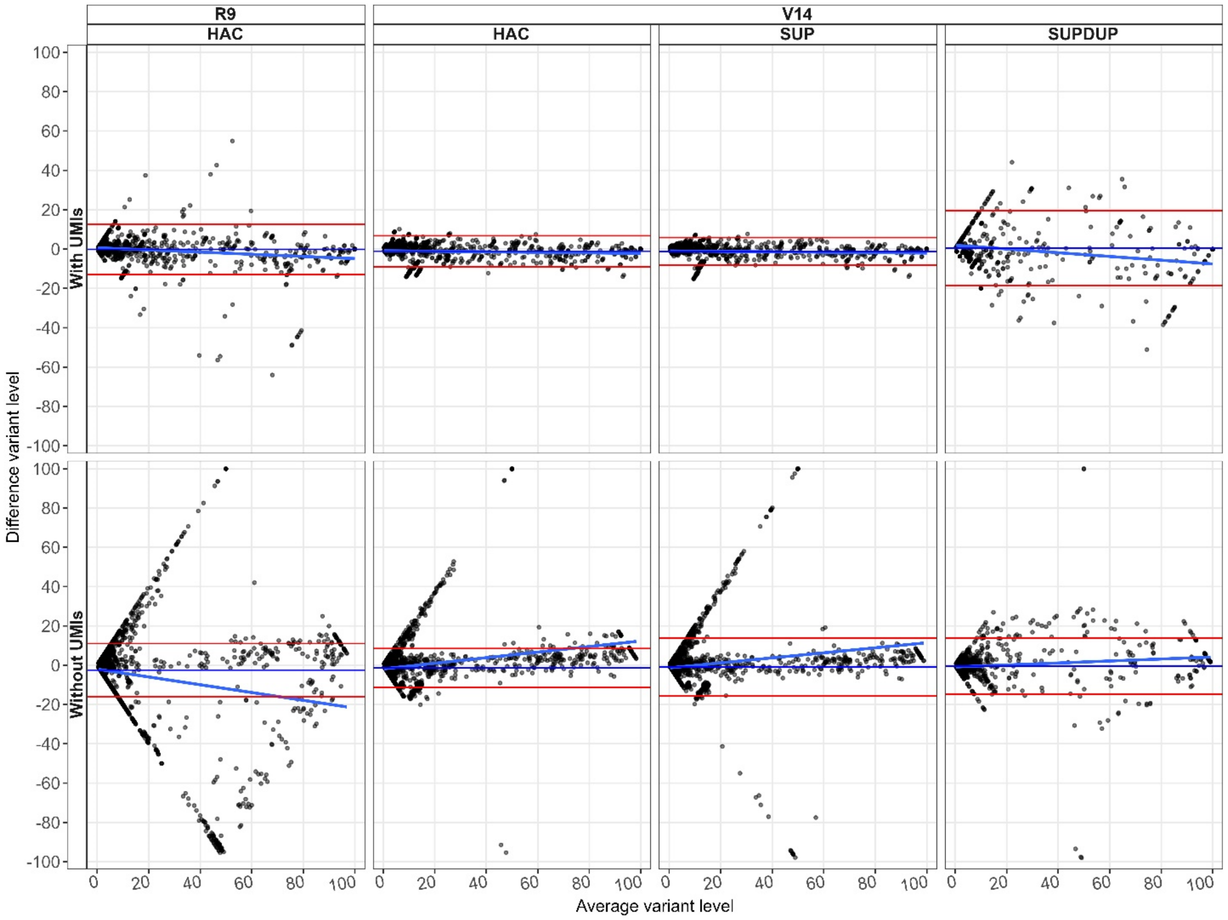

**Supplementary figure 9 Bland-Altman plot for ONT-Seq and UMI-ONT-Seq compared to NGS for all sequencing chemistries and basecalling algorithms used.**

Black points are variants found in NGS, ONT-Seq (top panels) or UMI-ONT-Seq (bottom panels). Without UMIs (using only the raw reads) we observed increasing correlation, decreasing bias for high level variants and narrower lines of agreement (95 % confidence interval) with increasing raw read accuracy (lowest to best: R9 HAC, V14 HAC, V14 SUP, V14 SUPDUP). For UMI-ONT-Seq we had generally high correlation between the variant levels compared to NGS for all conditions, narrower lines of agreement and, for V14 HAC and SUP, also a negligible bias.

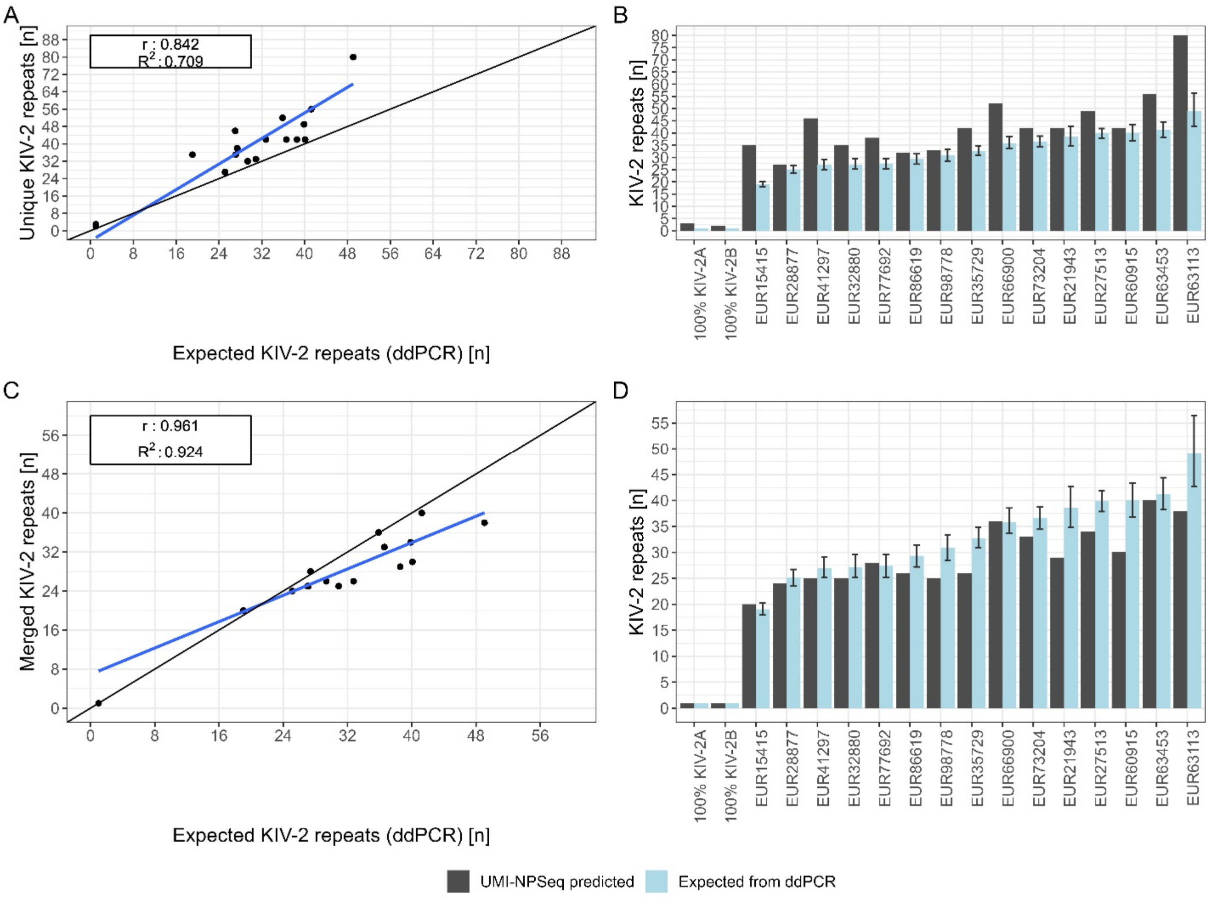

**Supplementary figure 10 Correlation of UMI-ONT-Seq estimated and actual number of KIV-2 repeats measured by ddPCR (expected) and haplotype cluster size distribution of 15 human gDNA samples and two 100% KIV-2A or KIV-2B plasmid.**

**Panels A to D** show the correlation (A, C) and absolute agreement (B, D) of the number of KIV-2 repeats derived from the number of unique haplotypes in UMI-ONT-Seq and as expected from ddPCR. Despite high correlation, for the majority of samples the number of KIV-2 repeats is overestimated. Panels C and D show the same correlation using a merging algorithm that removes and merges highly unlikely haplotype cluster sizes.

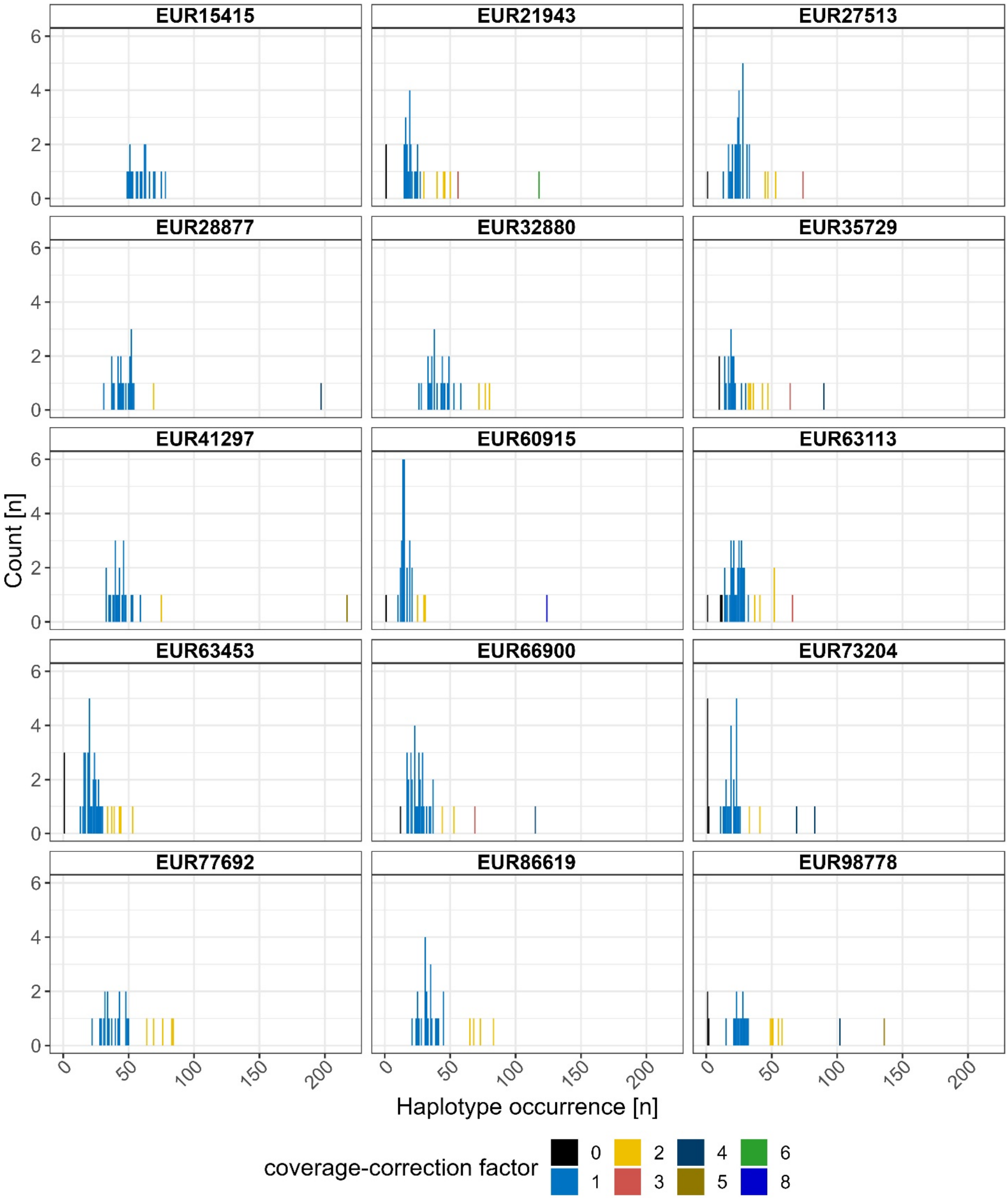

**Supplementary figure 11 Occurrence of the different haplotypes for the 15 SAPHIR samples.**

We observed haplotype occurrences that were not according to the assumed binomial distribution, suggesting the occurrence of identical KIV-2 repeats (e.g. recent repeat expansions). This was leveraged to determine a coverage-correction factor dependent on the occurrence of each haplotype per sample.

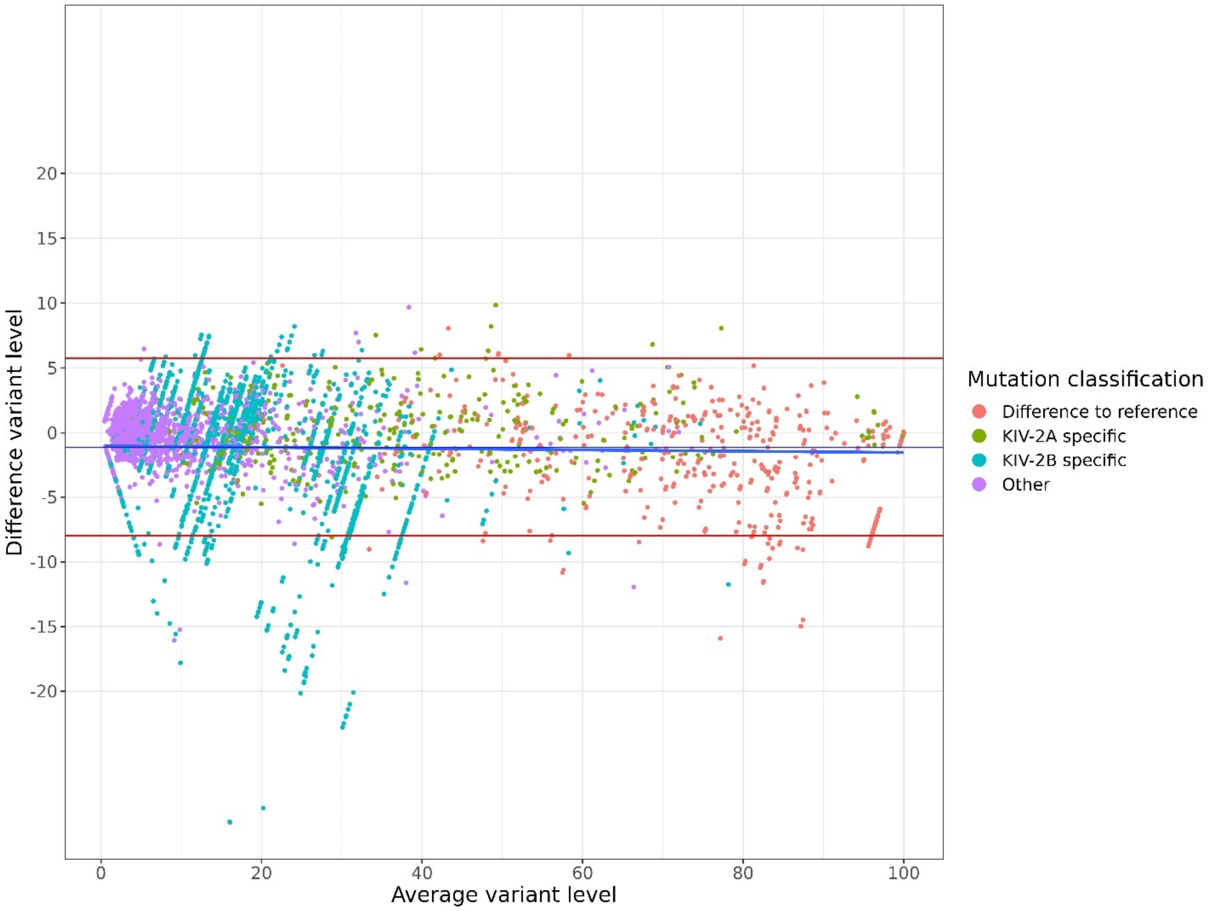

**Supplementary figure 12 Bland Altman plot of the 48 1000G samples analyzed with UMI-ONT-Seq compared variant levels derived from publically available WGS data[8].**

Mutations were classified according to their specificity for certain KIV-2 subtypes according to [4] (red: Difference to the reference sequence [No specific subtype], green: KIV-2A specific, blue: KIV-2B specific, purple: Not classified]. Variant levels were highly correlated, showed no bias, high accordance for all variant levels and low margins of error (< 5 %). A subset of KIV-2B specific mutations was found only with UMI-ONT-Seq and therefore showed higher deviation between both methods (turquoise points).

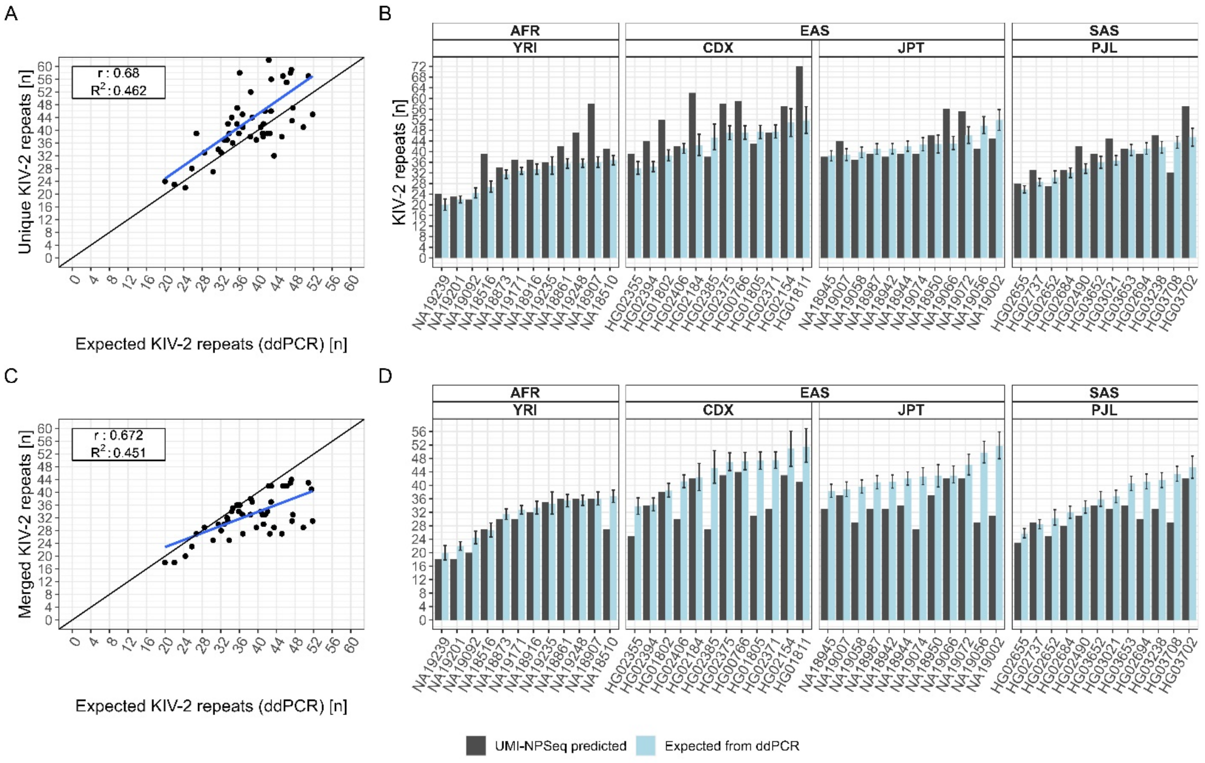

**Supplementary figure 13 Correlation of expected (ddPCR) and UMI-ONT-Seq predicted number of KIV-2 for different steps of the haplotype extraction and number of KIV-2 repeat prediction in 1000G.**

**Panels A and B** show the correlation and absolute concordance when only the uniquely occurring haplotypes are used for prediction. Taking only the unique haplotypes the number of KIV-2 repeats is generally overestimated and low correlation (r: 0.68, $R^2$: 0.462). **Panels C and D** show the correlation and absolute concordance after merging small haplotype clusters (occurrence < 0.85% of all consensus sequences) to cluster within a maximum of 1 edit distance and removing all small clusters that were not able to be merged. After that no sample showed more than the expected number of KIV-2 repeats and correlation increased to 0.732 (r) and 0.536 ($R^2$). For about one third of the samples the number of KIV-2 repeats determined by UMI-ONT-Seq is indeed within the 95% confidence interval of the ddPCR. This is improved further after coverage-correction (see **Fehler! Verweisquelle konnte nicht gefunden werden.**).

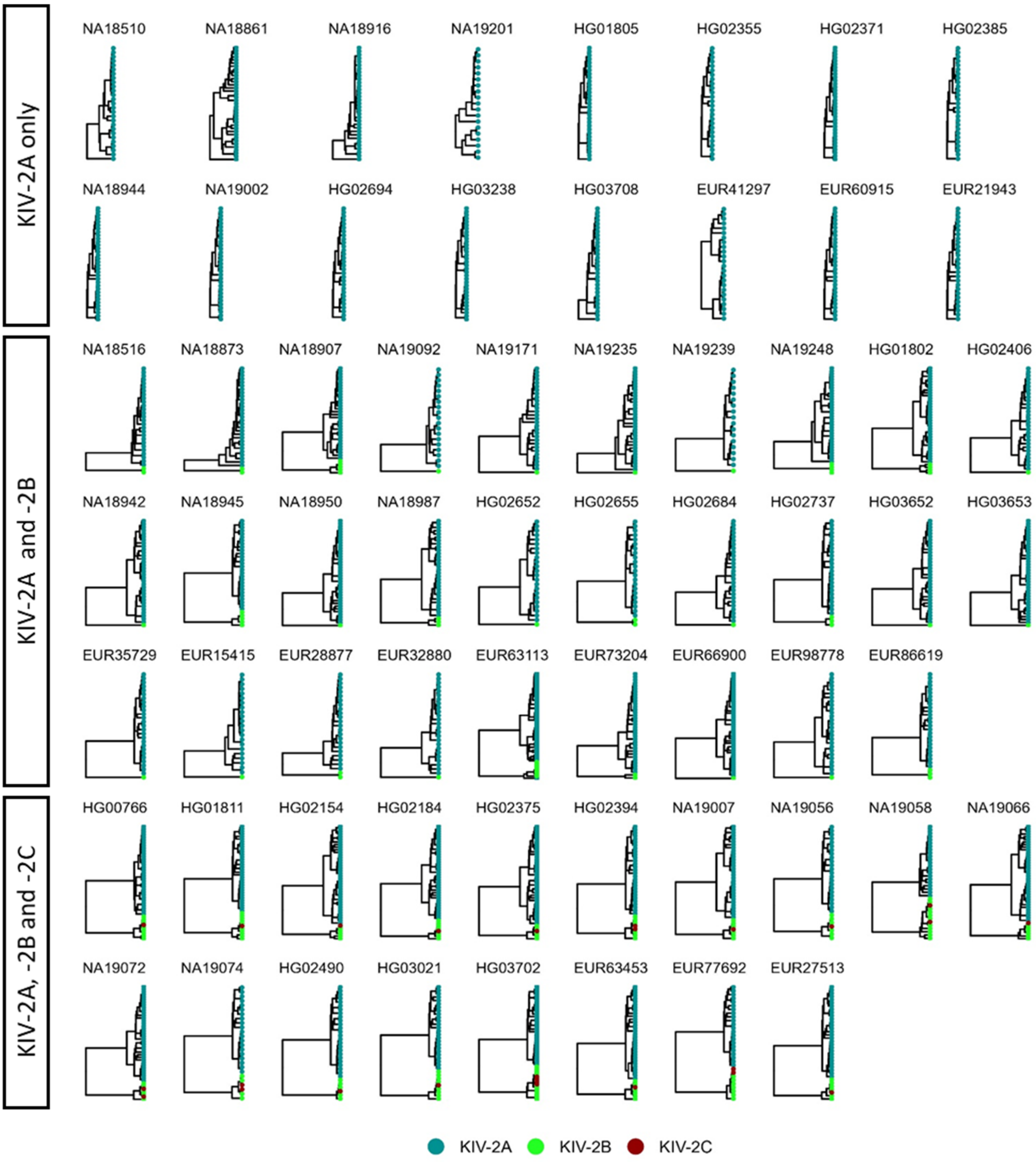

**Supplementary figure 14 KIV-2 subtype specific haplotype diversity of all 63 human gDNA samples in the present study (1000G, SAPHIR).**

KIV-2 subtype-specific haplotype diversity. Splitting the samples by their KIV-2 subtypes, revealed the 2 major clusters representing KIV-2A and the phylogenetically distant KIV-2B and C subtypes. While the KIV-2B and C cluster contain very similar sequences, with low diversity, the KIV-2A cluster has large internal differentiation. EUR: SAPHIR samples. Other samples: 1000G. See

Supplementary table 26 for population assignment of the 1000G sample IDs.

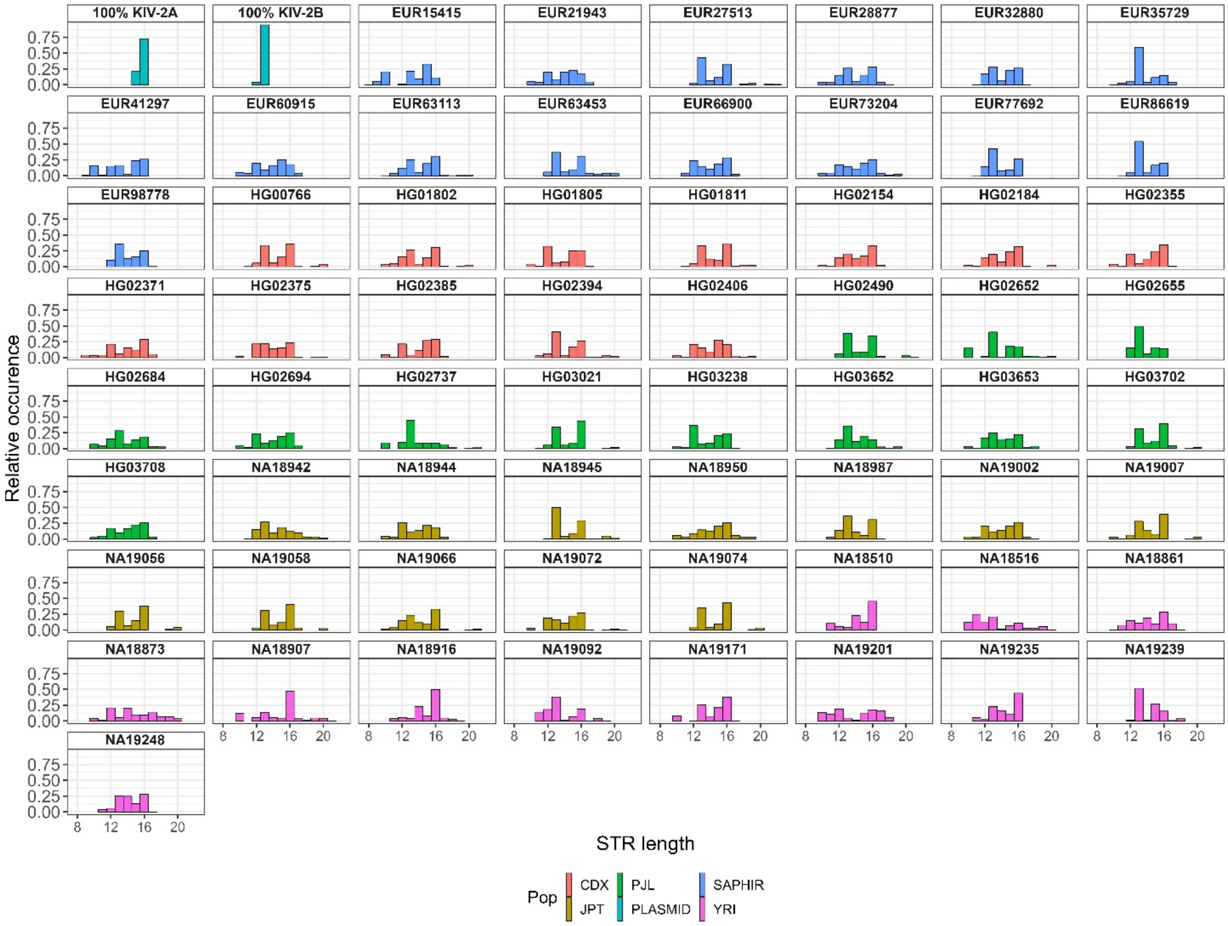

**Supplementary figure 14 Length distribution (relative occurrence) of the extracted STR-region of 63 gDNA samples of 5 different populations (1000G and SAPHIR) and the two unmixed plasmids.**

Using the high-quality UMI-ONT-Seq consensus sequences, we extracted 62,679 STR regions and counted the number of CA-repeats. The STR-region contains 7 to 22 repeats. While the STR-length was homogenous throughout all populations, a degeneration of the STR at position 7 and 15 suggest potential population specific pattern (Supplementary table 30 and Supplementary table 31).

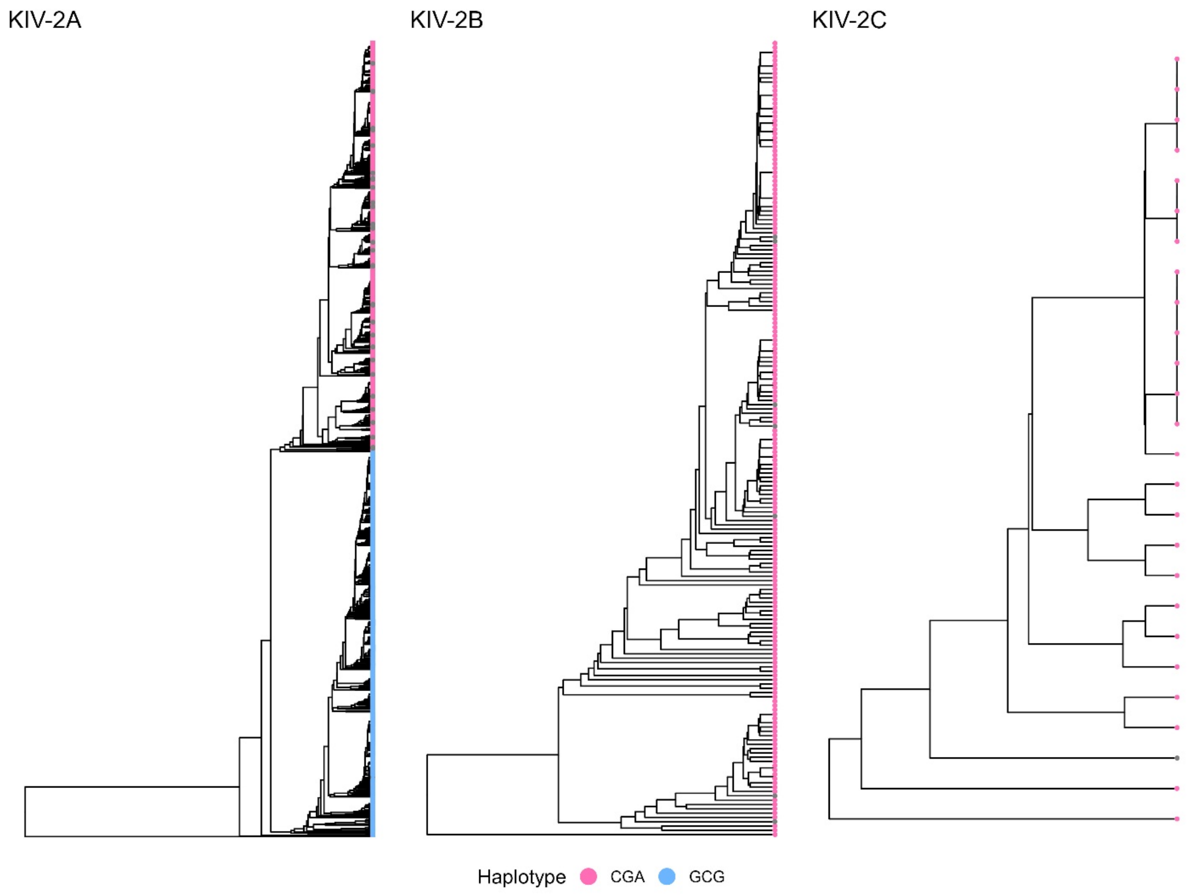

**Supplementary figure 15 Classification of KIV-2A repeat subclusters using 3 positions (35, 3103, 4358) of a previously suggested haplotype in a multi-ancestry sample set (1000G and SAPHIR) of 63 samples.**

Using three positions (35, 3103, 4358) of a previously suggested haplotype we were readily able to divide KIV-2A into the two main subclusters (CGA: pink or GCG: blue) observed in our analysis. All KIV-2B and KIV-2 repeats showed the same haplotype at these positions in all 63 samples.

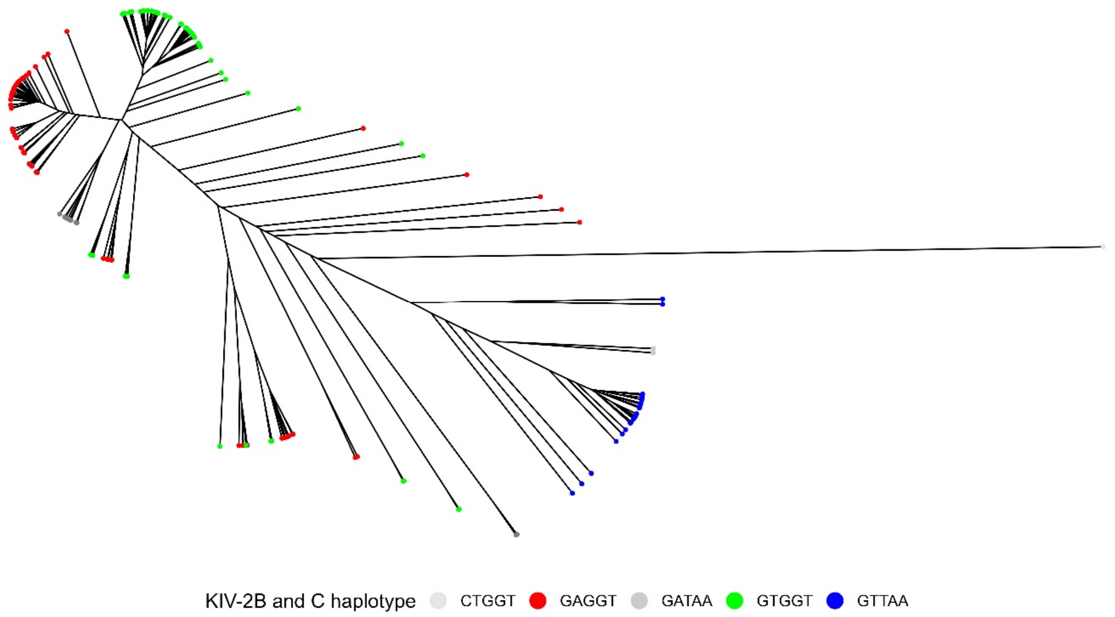

**Supplementary figure 16 Classification of KIV-2B and KIV-2C repeats into three main subclusters using 5 positions (50, 2409, 5037, 5045, 5052) in a multi-ancestry sample set (1000G and SAPHIR) of 63 samples.**

We observed 5 positions that split KIV-2B and KIV-2C repeats into three main subclusters. Interestingly, KIV-2C repeats were not found to build a distinct cluster. Noteworthy, the three haplotypes GAGGT, GTGGT and GTTAA represented 98.5 % of all found haplotypes.

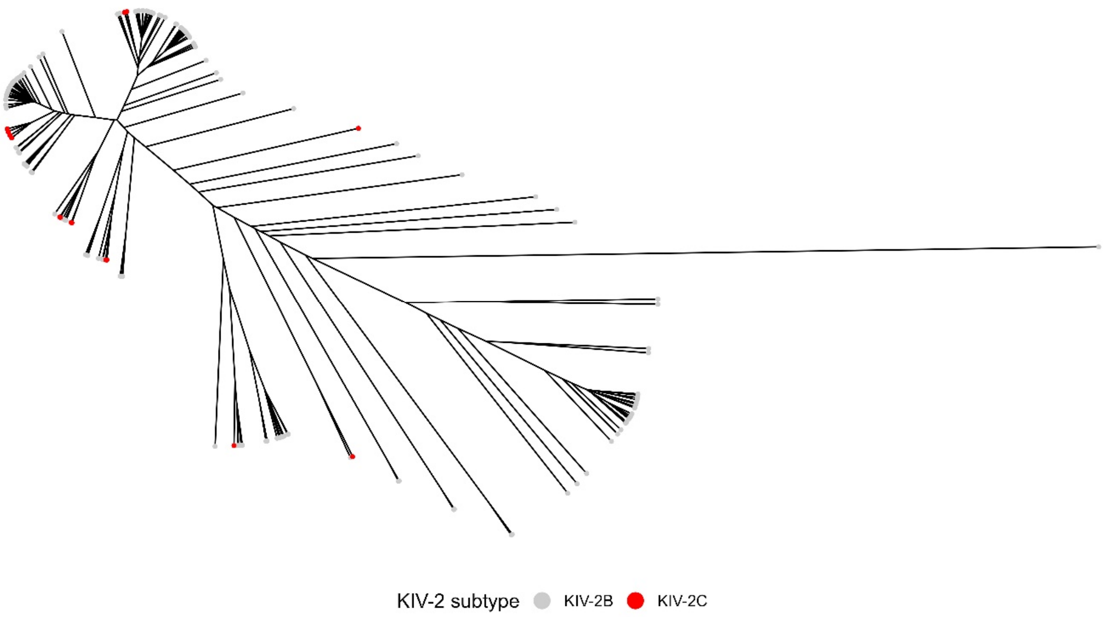

**Supplementary figure 17 Classification of KIV-2C containing repeats with the commonly used KIV-2 subtype defining positions (594, 621, 666) in a multi-ancestry sample set (1000G and SAPHIR) of 63 samples.**

Despite being associated with KIV-2B and KIV-2C, the three positions are not sufficient two define a distinct KIV-2C subcluster in all KIV-2B and KIV-2C containing repeats. KIV-2C can be found in several subclusters.

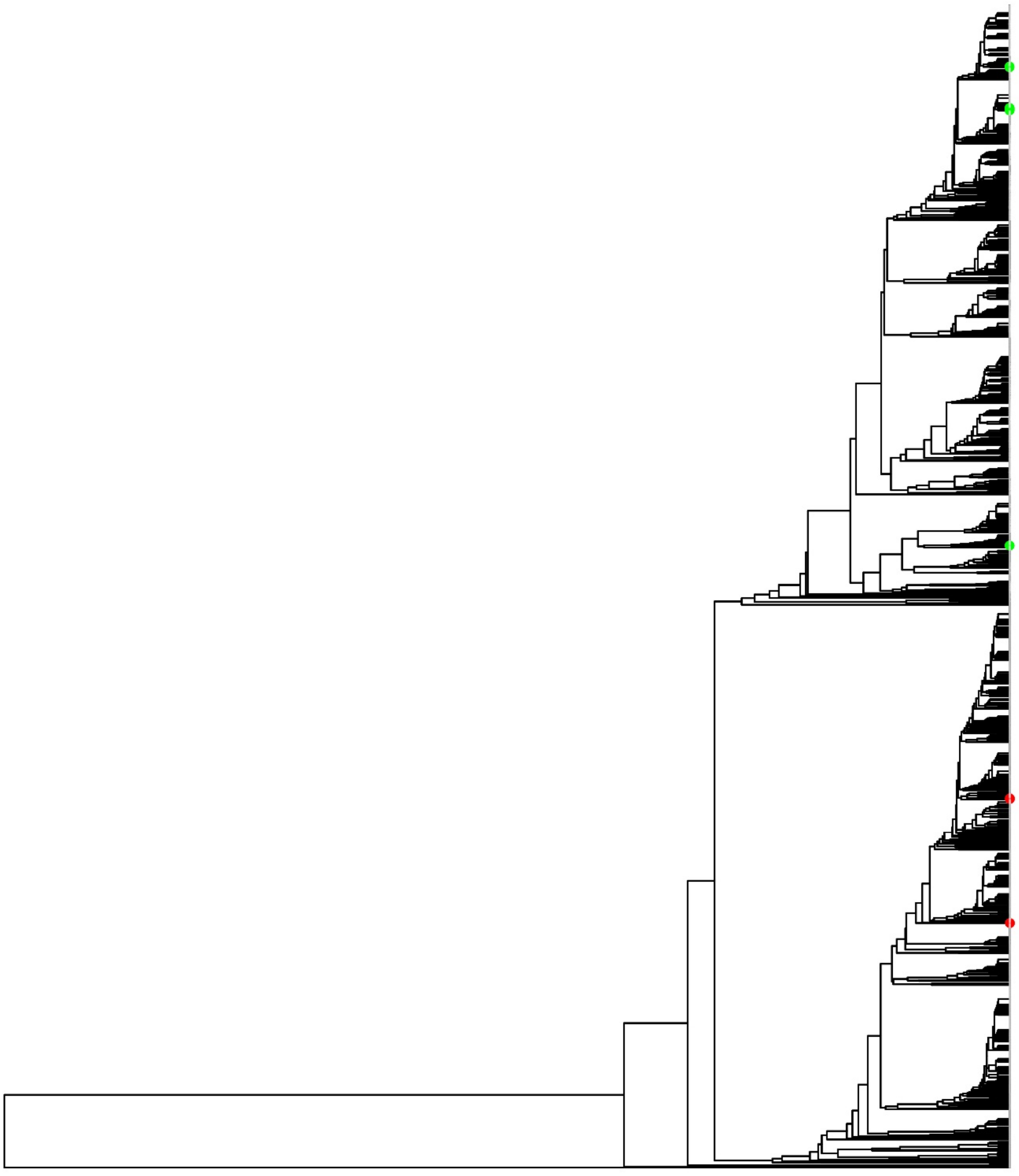

**Supplementary figure 18 Classification of the two most frequent disease relevant variants in the KIV-2 VNTR (4925G>A: red and 4733G>A: green) in a multi-ancestry sample set (1000G and SAPHIR) of 63 samples.**

The majority of the repeats was wild type for both alleles (GG: grey). 7 samples were found the carry either 4925G>A and 4733G>A. The two variants were solely found in KIV-2A repeats and associate with two different KIV-2A subclusters. Of 7 samples carrying either one of the two mutations 6 were European and 1 Punjabi. We observed 100% inter-individual sequence identity of the complete KIV-2 repeat in two (4925G>A) and three (4733G>A) samples.

### Supplementary tables

**Supplementary table 1 Primer sequences.** The tagging primers have a universal amplification sequence, the UMI sequence and the KIV-2 5104 or 2645 fragment-specific primer sequence. The amplification primer are complementary to the universal primer sequence of the tagging primer.

| ID | Target | Orientation | Step | Sequence (5‘-3‘) |
|---|---|---|---|---|
| 1 | PCR2645 | forward | tagging | GTATCGTGTAGAGACTGCGTAGGTTTVVVVTTVVVVTTVVVVTTVVVV TTTAGAAACAAACCTACTAAACCTGACAG |
| 2 | PCR2645 | reverse | tagging | AGTGATCGAGTCAGTGCGAGTGTTTVVVVTTVVVVTTVVVVTTVVVVT TTTTTTTCTGACAATCGGAATATAC |
| 3 | PCR2645 | forward | amplification | GGTGCTGAAGAAAGTTGTCGGTGTCTTTGTGTTAACCGTATCGTGTAG AGACTGCGTAGG |
| 4 | PCR2645 | reverse | amplification | GGTGCTGAAGAAAGTTGTCGGTGTCTTTGTGTTAACCAGTGATCGAGT CAGTGCGAGTG |
| 5 | PCR5104 | forward | tagging | GTATCGTGTAGAGACTGCGTAGGTTTVVVVTTVVVVTTVVVVTTVVVV TTTTCAGGATGCAGGGCATGAG |
| 6 | PCR5104 | reverse | tagging | AGTGATCGAGTCAGTGCGAGTGTTTVVVVTTVVVVTTVVVVTTVVVVT TTCACCAGAAATCACTCCGCTG |
| 7 | PCR5104 | forward | amplification | GGTGCTGAAGAAAGTTGTCGGTGTCTTTGTGTTAACCGTATCGTGTAG AGACTGCGTAGG |
| 8 | PCR5104 | reverse | amplification | GGTGCTGAAGAAAGTTGTCGGTGTCTTTGTGTTAACCAGTGATCGAGT CAGTGCGAGTG |

**Supplementary table 2 PCR conditions for the 2645 and the 5104 fragment.**

| | **PCR2645 - tagging** | **PCR2645 - early PCR** | **PCR2645 - late PCR** | **PCR5104 - tagging** | **PCR5104 - early PCR** | **PCR5104 - late PCR** |
|---|---|---|---|---|---|---|
| Product length [bp] | 2746 | 2820 | 2820 | 5205 | 5279 | 5279 |
| Reaction Volume [µL] | 15 | 50 | 50 | 15 | 50 | 50 |
| Enzyme | Superfi II [a] | Superfi II [a] | Superfi II [a] | Superfi II [a] | Superfi II [a] | Superfi II [a] |
| Initial Denaturation | 98 °C, 180 s, | 98 °C, 180 s, | 98 °C, 180 s, | 98 °C, 180 s, | 98 °C, 180 s, | 98 °C, 180 s, |
| Denaturation | 98 °C, 30 s, | 98 °C, 20 s, | 98 °C, 20 s, | 98 °C, 30 s, | 98 °C, 20 s, | 98 °C, 20 s, |
| Annealing, ramp rate | 66 - 60 °C, 90 s, 0.2 °C/s | 1 s 70°C, 70 - 63 °C, 45 s, 0.4 °C/s 1 s 63°C | - | 66 - 60 °C, 90 s, 0.2 °C/s | 1 s 70°C, 70 - 63 °C, 45 s, 0.4 °C/s 1 s 63°C | - |
| Extension | 72 °C, 90 s, | 72 °C, 90 s, | 72 °C, 120 s, | 72 °C, 180 s, | 72 °C, 180 s, | 72 °C, 240 s, |
| Cycles | 2 | 5 | 28 | 2 | 5 | 28 |
| Denaturation 2 | - | 98 °C, 20 s, | - | - | 98 °C, 20 s, | - |
| Extension 2 | - | 72 °C, 120 s, | - | - | 72 °C, 180 s, | - |
| Cycles | 1 | 5 | | 1 | 5 | |
| Final extension | 72 °C, 300 s, | 72 °C, 300 s, | 72 °C, 300 s, | 72 °C, 300 s, | 72 °C, 300 s, | 72 °C, 300 s, |
| Primer | 1, 2 [b] | 3, 4 [b] | 3, 4 [b] | 5, 6 [b] | 7, 8 [b] | 7, 8 [b] |
| Final primer conc. [nM] | 750 | 830 | 830 | 750 | 830 | 830 |
| Final MgCl2 conc. [mM] | - | 3.3 | 3.3 | - | 3.3 | 3.3 |
| Sample input | 2 ng | 21 µl | 21 µl | 2 ng | 21 µl | 21 µl |
| Targets | 50,000 | - | - | 50,000 | - | - |

[a] Full name of the enzyme: ThermoFisher Scientific Platinum SuperFi II Green PCR Master Mix.

[b] PrimerIDs refer to Supplementary table 1.

**Supplementary table 3 ddPCR validation of the two unmixed and the plasmid mixtures for both fragments. Abundance is the fractional abundance of KIV-2B specific alleles in the plasmid mixtures. CI: Confidence interval**

| Sample | Fragment | Abundance [%] | Abundance CI 95% - Low | Abundance CI 95% - High |
|---|---|---|---|---|
| 100 % KIV-2A | 2645 | 99.75 | 99.94 | 100.44 |
| 100 % KIV-2B | 2645 | 99.88 | 99.63 | 100.12 |
| 5% | 2645 | 5.29 | 4.66 | 5.93 |
| 2.50% | 2645 | 3.21 | 2.28 | 4.04 |
| 1.50% | 2645 | 1.65 | 1.16 | 2.18 |
| 1% | 2645 | 1.33 | 1.02 | 1.64 |
| 0.5% | 2645 | 0.68 | 0.59 | 0.77 |
| 100 % KIV-2A | 5104 | 99.14 | 99.58 | 101.31 |
| 100 % KIV-2B | 5104 | 99.94 | 99.79 | 100.08 |
| 5% | 5104 | 5.43 | 4.69 | 6.16 |
| 2.50% | 5104 | 2.97 | 2.16 | 3.77 |
| 1.50% | 5104 | 1.39 | 0.8 | 1.99 |
| 1% | 5104 | 1.18 | 0.67 | 1.69 |
| 0.5% | 5104 | 0.62 | 0.5 | 0.74 |

**Supplementary table 4 ddPCR and qPCR reaction conditions.** MGBEQ: minor groove binding black hole quencher. The ddPCR for KIV-2A/KIV-2B plasmid mixture validation uses two probes binding in LPA KIV-2 exon 1 at a base that differs between the two subtypes.

| | **KIV-2 copy number ddPCR** | **KIV2B quantification ddPCR** |
|---|---|---|
| Product length [bp] | 94 | 79 |
| Reaction volume [µL] | 20 | 20 |
| Enzyme | Biorad ddPCR 2X Supermix for Probes (No dUTP) | Biorad ddPCR 2X Supermix for Probes (No dUTP) |
| Forward primer (5‘-3‘) [a] | GTGGCAGCTCCTTATTGTTATACG | TTTTCATTTCAGCACCGACTGA |
| Reverse primer (5‘-3‘) [a] | CGACGGCAGTCCCTTCTG | GTGCCTCGATAACTCTGTCCATT |
| Probe (5‘-3‘) [a] | [FAM]-TACTGCAACCTGACGCAATGCTCAGAC-[BHQ1] | [HEX]-AGGAGTGCTACCACGG-[MGBEQ]<br>[FAM]-AGGAGTGCTACCATGGT-[MGBEQ] |
| Diploid reference assay | ThermoFisher RNAse P Copy Number Reference assay<br>(Part.no 4403328) | none |
| Final primer concentration [nM] | 900 nM | 900 nM |
| Final LPA probe concentration [nM] | 300 nM | 300 nM |
| Initial denaturation | 95 °C, 10 min | 95 °C, 10 min |
| Denaturation | 94 °C, 30 s, | 94 °C, 30 s |
| Annealing (ramp rate) | 60 °C, 60 s (2 °C/s) | 57.4 °C, 30 s (2 °C/s) |
| Extension step | None, 2 step PCR | 60.0 °C, 30 sec |
| Cycles | 40 | 40 |
| Final denaturation | 98°C, 10 min | 98°C, 10 min |
| Sample input [ng] | 4 ng EcoRI digested | 4 ng Plasmid |

**Supplementary table 5 Summary statistics of the sequencing results per kit and basecalling algorithm.**

| Kit | Algorithm | Number of reads | Median Q-score | Mean Q-score | Number of bases [Gb] | Number of samples |
|---|---|---|---|---|---|---|
| R9 | HAC | 3509445 | 13.2 | 13.1 | 14.7 | 29 |
| V14 | HAC | 3227736 | 15.7 | 15.3 | 13.8 | 29 |
| V14 | SUP | 3548652 | 18.5 | 17.9 | 14 | 29 |
| V14 | SUPDUP | 277911 | 28.6 | 28 | 1.4 | 12 |
| Total | - | 10563744 | - | - | 43.9 | 99 |

**Supplementary table 6 Sequencing overview R9 HAC.**

| Sample | Fragment | Number of reads | Number of bases [Mb] | Median Q-score | Mean Q-score | Median read length [b] | Mean read length [b] |
|---|---|---|---|---|---|---|---|
| 100% KIV-2B | 2645 | 109843 | 303.5 | 12.1 | 12.1 | 2911 | 2762.7 |
| 5% | 2645 | 126532 | 359.7 | 12 | 12.1 | 2911 | 2842.6 |
| 2.50% | 2645 | 113217 | 323.4 | 12.4 | 12.4 | 2913 | 2856.1 |
| 1.50% | 2645 | 93180 | 264.6 | 12.4 | 12.3 | 2913 | 2839.2 |
| 1% | 2645 | 101180 | 288 | 12.4 | 12.3 | 2912 | 2846.4 |
| 0.50% | 2645 | 137340 | 388.5 | 12 | 12.1 | 2912 | 2828.5 |
| 100% KIV-2A | 2645 | 134277 | 383.1 | 12 | 12.1 | 2913 | 2853.3 |
| 100% KIV-2B | 5104 | 125709 | 639.7 | 13.8 | 13.6 | 5335 | 5088.9 |
| 5% | 5104 | 127595 | 602.2 | 12.7 | 12.6 | 5283 | 4719.3 |
| 2.50% | 5104 | 77479 | 397 | 12.1 | 12.1 | 5333 | 5123.4 |
| 1.50% | 5104 | 106608 | 535.4 | 13.8 | 13.5 | 5327 | 5022.3 |
| 1% | 5104 | 105730 | 497.6 | 12.7 | 12.6 | 5281 | 4705.9 |
| 0.50% | 5104 | 152135 | 770 | 13.8 | 13.5 | 5336 | 5061 |
| 100% KIV-2A | 5104 | 163591 | 819.8 | 13.8 | 13.5 | 5339 | 5011.3 |
| EUR15415 | 5104 | 70038 | 317.1 | 14 | 13.7 | 5308 | 4527.4 |
| EUR21943 | 5104 | 100754 | 492 | 13.4 | 13.2 | 5329 | 4883 |
| EUR27513 | 5104 | 88489 | 389.9 | 13.9 | 13.7 | 5321 | 4406.6 |
| EUR28877 | 5104 | 156027 | 771.2 | 13.5 | 13.3 | 5342 | 4943 |
| EUR32880 | 5104 | 151916 | 739.8 | 13.5 | 13.3 | 5339 | 4869.8 |
| EUR35729 | 5104 | 95958 | 469.7 | 13.5 | 13.3 | 5328 | 4895 |
| EUR41297 | 5104 | 128104 | 628.8 | 13.5 | 13.3 | 5336 | 4908.3 |
| EUR60915 | 5104 | 110100 | 529.6 | 13.5 | 13.2 | 5329 | 4810.1 |
| EUR63113 | 5104 | 157136 | 689.4 | 14 | 13.7 | 5331 | 4387.3 |
| EUR63453 | 5104 | 114891 | 518.8 | 13.5 | 13.3 | 5323 | 4516 |
| EUR66900 | 5104 | 93850 | 397 | 13.9 | 13.7 | 5313 | 4229.7 |
| EUR73204 | 5104 | 208536 | 494.3 | 13.8 | 13.6 | 1500 | 2370.2 |
| EUR77692 | 5104 | 147495 | 640.1 | 13.9 | 13.7 | 5326 | 4339.7 |
| EUR86619 | 5104 | 97587 | 485.7 | 13.9 | 13.7 | 5343 | 4977.6 |
| EUR98778 | 5104 | 114148 | 560.6 | 13.9 | 13.7 | 5343 | 4911 |

**Supplementary table 7 Sequencing overview V14 HAC.**

| Sample | Fragment | Number of reads | Number of bases [Mb] | Median Q-score | Mean Q-score | Median read length [b] | Mean read length [b] |
|---|---|---|---|---|---|---|---|
| 100% KIV-2B | 2645 | 105403 | 307.2 | 14.8 | 14.5 | 2940 | 2914.9 |
| 5% | 2645 | 110703 | 304.1 | 18.9 | 18.4 | 2817 | 2746.6 |
| 2.50% | 2645 | 105147 | 289.5 | 18.9 | 18.4 | 2817 | 2753 |
| 1.50% | 2645 | 94187 | 261.1 | 18.9 | 18.3 | 2817 | 2772.2 |
| 1% | 2645 | 131673 | 368.2 | 14.9 | 14.5 | 2937 | 2796.7 |
| 0.50% | 2645 | 91836 | 243.5 | 18.9 | 18.4 | 2816 | 2652 |
| 100% KIV-2A | 2645 | 96155 | 261.1 | 18.8 | 18.3 | 2816 | 2715.3 |
| 100% KIV-2B | 5104 | 145038 | 576.7 | 14.9 | 14.6 | 5363 | 3976.5 |
| 5% | 5104 | 107300 | 533.4 | 15.3 | 14.9 | 5385 | 4970.8 |
| 2.50% | 5104 | 127751 | 636.9 | 15.4 | 14.9 | 5385 | 4985.1 |
| 1.50% | 5104 | 110409 | 528.8 | 15 | 14.6 | 5381 | 4789.9 |
| 1% | 5104 | 113702 | 568.5 | 15.4 | 15 | 5386 | 5000.3 |
| 0.50% | 5104 | 94354 | 485.7 | 15.4 | 15 | 5387 | 5148.1 |
| 100% KIV-2A | 5104 | 104810 | 513.1 | 15.3 | 14.9 | 5384 | 4895.2 |
| EUR15415 | 5104 | 107148 | 530.3 | 15 | 14.6 | 5378 | 4949.6 |
| EUR21943 | 5104 | 115224 | 572.5 | 15 | 14.7 | 5380 | 4968.4 |
| EUR27513 | 5104 | 90414 | 415.1 | 14.9 | 14.6 | 5375 | 4591.1 |
| EUR28877 | 5104 | 106609 | 532.9 | 15.1 | 14.7 | 5382 | 4998.3 |
| EUR32880 | 5104 | 103952 | 515.9 | 15.1 | 14.7 | 5381 | 4962.6 |
| EUR35729 | 5104 | 100376 | 500.7 | 15.1 | 14.7 | 5381 | 4987.8 |
| EUR41297 | 5104 | 114752 | 573.3 | 15.1 | 14.7 | 5381 | 4996.1 |
| EUR60915 | 5104 | 101977 | 503.7 | 15 | 14.6 | 5380 | 4939.2 |
| EUR63113 | 5104 | 100513 | 457.5 | 15 | 14.6 | 5376 | 4551.5 |
| EUR63453 | 5104 | 99587 | 470.5 | 15 | 14.6 | 5380 | 4724.2 |
| EUR66900 | 5104 | 112271 | 501.4 | 15 | 14.7 | 5374 | 4465.6 |
| EUR73204 | 5104 | 157966 | 441.8 | 14.5 | 14.3 | 1525 | 2796.7 |
| EUR77692 | 5104 | 102688 | 467.9 | 14.9 | 14.6 | 5376 | 4556.9 |
| EUR86619 | 5104 | 116328 | 589.5 | 15 | 14.6 | 5381 | 5067.6 |
| EUR98778 | 5104 | 159463 | 803.3 | 15 | 14.7 | 5381 | 5037.3 |

**Supplementary table 8 Sequencing overview V14 SUP.**

| Sample | Fragment | Number of reads | Number of bases [Mb] | Median Q-score | Mean Q-score | Median read length [b] | Mean read length [b] |
|---|---|---|---|---|---|---|---|
| 100% KIV-2B | 2645 | 110348 | 305.2 | 18.2 | 17.7 | 2819 | 2766.2 |
| 5% | 2645 | 110703 | 304.1 | 18.9 | 18.4 | 2817 | 2746.6 |
| 2.50% | 2645 | 105147 | 289.5 | 18.9 | 18.4 | 2817 | 2753 |
| 1.50% | 2645 | 94187 | 261.1 | 18.9 | 18.3 | 2817 | 2772.2 |
| 1% | 2645 | 133275 | 359.5 | 18.5 | 18 | 2816 | 2697.5 |
| 0.50% | 2645 | 91836 | 243.5 | 18.9 | 18.4 | 2816 | 2652 |
| 100% KIV-2A | 2645 | 96155 | 261.1 | 18.8 | 18.3 | 2816 | 2715.3 |
| 100% KIV-2B | 5104 | 179155 | 610.7 | 18.4 | 17.8 | 4788 | 3408.7 |
| 5% | 5104 | 109517 | 531.2 | 18.8 | 18.1 | 5268 | 4850.2 |
| 2.50% | 5104 | 130071 | 632.5 | 18.8 | 18.1 | 5267 | 4862.9 |
| 1.50% | 5104 | 146157 | 569.6 | 18.2 | 17.7 | 5256 | 3896.9 |
| 1% | 5104 | 115813 | 565.1 | 18.8 | 18.2 | 5267 | 4879.6 |
| 0.50% | 5104 | 96259 | 484.2 | 18.9 | 18.3 | 5269 | 5030 |
| 100% KIV-2A | 5104 | 106702 | 509.7 | 18.8 | 18.2 | 5267 | 4776.4 |
| EUR15415 | 5104 | 109926 | 532.2 | 18.4 | 17.7 | 5263 | 4841.9 |
| EUR21943 | 5104 | 117770 | 572.8 | 18.3 | 17.7 | 5263 | 4863.5 |
| EUR27513 | 5104 | 106423 | 432.8 | 18.1 | 17.5 | 5256 | 4067 |
| EUR28877 | 5104 | 108763 | 532.7 | 18.4 | 17.8 | 5264 | 4897.5 |
| EUR32880 | 5104 | 106256 | 516.1 | 18.4 | 17.8 | 5264 | 4857.5 |
| EUR35729 | 5104 | 102554 | 501 | 18.4 | 17.7 | 5264 | 4885.3 |
| EUR41297 | 5104 | 117428 | 574.3 | 18.5 | 17.8 | 5264 | 4890.7 |
| EUR60915 | 5104 | 104458 | 504.7 | 18.4 | 17.7 | 5263 | 4832.1 |
| EUR63113 | 5104 | 119810 | 478 | 18.3 | 17.6 | 5255 | 3989.6 |
| EUR63453 | 5104 | 101851 | 471.3 | 18.4 | 17.7 | 5264 | 4627.8 |
| EUR66900 | 5104 | 138525 | 528.8 | 18.3 | 17.7 | 5249 | 3817 |
| EUR73204 | 5104 | 261885 | 556.5 | 18.2 | 17.7 | 1389 | 2124.8 |
| EUR77692 | 5104 | 124413 | 491.4 | 18.2 | 17.5 | 5253 | 3949.7 |
| EUR86619 | 5104 | 125810 | 599 | 18.2 | 17.5 | 5262 | 4760.8 |
| EUR98778 | 5104 | 177455 | 822.6 | 18.3 | 17.6 | 5262 | 4635.6 |

**Supplementary table 9 Sequencing overview V14 SUPDUP.**

| Sample | Fragment | Number of reads | Number of bases [Mb] | Median Q-score | Mean Q-score | Median read length [b] | Mean read length [b] |
|---|---|---|---|---|---|---|---|
| EUR15415 | 5104 | 29849 | 155.9 | 28.7 | 28.1 | 5273 | 5223 |
| EUR21943 | 5104 | 29657 | 154.8 | 29.1 | 28.4 | 5272 | 5218.5 |
| EUR28877 | 5104 | 30581 | 160.3 | 29 | 28.3 | 5273 | 5242.2 |
| EUR32880 | 5104 | 28577 | 149.2 | 28.9 | 28.2 | 5273 | 5219.7 |
| EUR35729 | 5104 | 24606 | 128.6 | 28.8 | 28.1 | 5273 | 5228.3 |
| EUR41297 | 5104 | 34236 | 179.2 | 29 | 28.3 | 5273 | 5234.4 |
| EUR60915 | 5104 | 26888 | 140.2 | 29.1 | 28.5 | 5272 | 5214.1 |
| EUR63113 | 5104 | 11125 | 49.7 | 28.4 | 28 | 5270 | 4463.5 |
| EUR63453 | 5104 | 25678 | 133 | 28.2 | 27.5 | 5275 | 5180.4 |
| EUR73204 | 5104 | 6685 | 8.1 | 25.9 | 25.8 | 1158 | 1206.9 |
| EUR86619 | 5104 | 12618 | 64.3 | 28.9 | 28.3 | 5271 | 5097.3 |
| EUR98778 | 5104 | 17411 | 87.4 | 28.9 | 28.4 | 5271 | 5019.7 |

**Supplementary table 10 Performance measures for all plasmid mixtures using UMI-ONT-Seq with the default clustering strategy.**

| Sample | Fragment | Kit | Algorithm | Raw reads | Consensus sequences | True positive | True negative | False positive | False negative | Sensitivity | Specificity | Precision | F1 score |
|---|---|---|---|---|---|---|---|---|---|---|---|---|---|
| 100 % KIV-2A | 2645 | R9 | HAC | 134277 | 600 | 8 | 2637 | 0 | 0 | 1 | 1 | 1 | 1 |
| 100 % KIV-2B | 2645 | R9 | HAC | 109843 | 495 | 112 | 2533 | 0 | 0 | 1 | 1 | 1 | 1 |
| 5% | 2645 | R9 | HAC | 103124 | 578 | 118 | 2527 | 0 | 0 | 1 | 1 | 1 | 1 |
| 2.50% | 2645 | R9 | HAC | 113217 | 618 | 118 | 2527 | 0 | 0 | 1 | 1 | 1 | 1 |
| 1.50% | 2645 | R9 | HAC | 93180 | 390 | 117 | 2527 | 0 | 1 | 0.992 | 1 | 1 | 0.996 |
| 1% | 2645 | R9 | HAC | 101180 | 507 | 118 | 2527 | 0 | 0 | 1 | 1 | 1 | 1 |
| 0.50% | 2645 | R9 | HAC | 137340 | 611 | 68 | 2527 | 0 | 50 | 0.576 | 1 | 1 | 0.731 |
| 100 % KIV-2A | 5104 | R9 | HAC | 163591 | 636 | 27 | 5077 | 0 | 0 | 1 | 1 | 1 | 1 |
| 100 % KIV-2B | 5104 | R9 | HAC | 125709 | 576 | 75 | 5029 | 0 | 0 | 1 | 1 | 1 | 1 |
| 5% | 5104 | R9 | HAC | 127595 | 496 | 88 | 5014 | 2 | 0 | 1 | 1 | 0.978 | 0.989 |
| 2.50% | 5104 | R9 | HAC | 77479 | 416 | 88 | 5015 | 1 | 0 | 1 | 1 | 0.989 | 0.994 |
| 1.50% | 5104 | R9 | HAC | 106608 | 461 | 88 | 5015 | 1 | 0 | 1 | 1 | 0.989 | 0.994 |
| 1% | 5104 | R9 | HAC | 105730 | 493 | 27 | 5015 | 1 | 61 | 0.307 | 1 | 0.964 | 0.466 |
| 0.50% | 5104 | R9 | HAC | 152135 | 555 | 27 | 5015 | 1 | 61 | 0.307 | 1 | 0.964 | 0.466 |
| 100 % KIV-2A | 2645 | V14 | HAC | 96155 | 628 | 8 | 2637 | 0 | 0 | 1 | 1 | 1 | 1 |
| 100 % KIV-2B | 2645 | V14 | HAC | 105403 | 310 | 112 | 2533 | 0 | 0 | 1 | 1 | 1 | 1 |
| 5% | 2645 | V14 | HAC | 110703 | 708 | 118 | 2527 | 0 | 0 | 1 | 1 | 1 | 1 |
| 2.50% | 2645 | V14 | HAC | 105147 | 818 | 118 | 2527 | 0 | 0 | 1 | 1 | 1 | 1 |
| 1.50% | 2645 | V14 | HAC | 94187 | 463 | 118 | 2527 | 0 | 0 | 1 | 1 | 1 | 1 |
| 1% | 2645 | V14 | HAC | 131673 | 387 | 118 | 2527 | 0 | 0 | 1 | 1 | 1 | 1 |
| 0.50% | 2645 | V14 | HAC | 91836 | 283 | 118 | 2526 | 1 | 0 | 1 | 1 | 0.992 | 0.996 |
| 100 % KIV-2A | 5104 | V14 | HAC | 81945 | 417 | 27 | 5077 | 0 | 0 | 1 | 1 | 1 | 1 |
| 100 % KIV-2B | 5104 | V14 | HAC | 145038 | 779 | 75 | 5029 | 0 | 0 | 1 | 1 | 1 | 1 |
| 5% | 5104 | V14 | HAC | 107300 | 435 | 88 | 5016 | 0 | 0 | 1 | 1 | 1 | 1 |
| 2.50% | 5104 | V14 | HAC | 127751 | 833 | 88 | 5015 | 1 | 0 | 1 | 1 | 0.989 | 0.994 |
| 1.50% | 5104 | V14 | HAC | 110409 | 659 | 88 | 5016 | 0 | 0 | 1 | 1 | 1 | 1 |
| 1% | 5104 | V14 | HAC | 113702 | 224 | 87 | 5013 | 3 | 1 | 0.989 | 0.999 | 0.967 | 0.978 |
| 0.50% | 5104 | V14 | HAC | 94354 | 343 | 27 | 5016 | 0 | 61 | 0.307 | 1 | 1 | 0.47 |
| 100 % KIV-2A | 2645 | V14 | SUP | 96155 | 627 | 8 | 2637 | 0 | 0 | 1 | 1 | 1 | 1 |

| | | | | | | | | | | | | | |
|---|---|---|---|---|---|---|---|---|---|---|---|---|---|
| 100 % KIV-2B | 2645 | V14 | SUP | 110348 | 348 | 112 | 2533 | 0 | 0 | 1 | 1 | 1 | 1 |
| 5% | 2645 | V14 | SUP | 110703 | 725 | 118 | 2527 | 0 | 0 | 1 | 1 | 1 | 1 |
| 2.50% | 2645 | V14 | SUP | 105147 | 828 | 118 | 2527 | 0 | 0 | 1 | 1 | 1 | 1 |
| 1.50% | 2645 | V14 | SUP | 94187 | 462 | 118 | 2527 | 0 | 0 | 1 | 1 | 1 | 1 |
| 1% | 2645 | V14 | SUP | 133275 | 399 | 118 | 2527 | 0 | 0 | 1 | 1 | 1 | 1 |
| 0.50% | 2645 | V14 | SUP | 91836 | 289 | 118 | 2526 | 1 | 0 | 1 | 1 | 0.992 | 0.996 |
| 100 % KIV-2A | 5104 | V14 | SUP | 83509 | 395 | 27 | 5077 | 0 | 0 | 1 | 1 | 1 | 1 |
| 100 % KIV-2B | 5104 | V14 | SUP | 179155 | 816 | 75 | 5029 | 0 | 0 | 1 | 1 | 1 | 1 |
| 5% | 5104 | V14 | SUP | 109517 | 451 | 88 | 5016 | 0 | 0 | 1 | 1 | 1 | 1 |
| 2.50% | 5104 | V14 | SUP | 130071 | 944 | 88 | 5016 | 0 | 0 | 1 | 1 | 1 | 1 |
| 1.50% | 5104 | V14 | SUP | 146157 | 732 | 88 | 5016 | 0 | 0 | 1 | 1 | 1 | 1 |
| 1% | 5104 | V14 | SUP | 115813 | 215 | 87 | 5013 | 3 | 1 | 0.989 | 0.999 | 0.967 | 0.978 |
| 0.50% | 5104 | V14 | SUP | 96259 | 346 | 28 | 5016 | 0 | 60 | 0.318 | 1 | 1 | 0.483 |

**Supplementary table 11 Edit distances within UMI sequences in the same UMI cluster using UMI-ONT-Seq with the default clustering strategy.**

| Fragment | Kit | Algorithm | Clusters | Max UMI edit distance in clusters | Mean UMI edit distance in clusters | SD UMI edit distance in clusters |
|---|---|---|---|---|---|---|
| PCR2645 | R9 | HAC | 28127 | 26 | 15.879 | 3.944 |
| PCR2645 | V14 | HAC | 15535 | 25 | 15.453 | 4.145 |
| PCR2645 | V14 | SUP | 14700 | 25 | 15.441 | 4.172 |
| PCR5104 | R9 | HAC | 46503 | 26 | 16.106 | 3.987 |
| PCR5104 | V14 | HAC | 35965 | 26 | 15.41 | 4.189 |
| PCR5104 | V14 | SUP | 32974 | 25 | 15.503 | 4.204 |

**Supplementary table 12 Variant levels and noise of mutation level using UMI-ONT-Seq with the default clustering strategy.**

| Sample | Fragment | Kit | Algorithm | Mean variant level [%] | SD variant level [%] | Absolute variant level range observed |
|---|---|---|---|---|---|---|
| 100 % KIV-2B | PCR2645 | R9 | HAC | 99.763 | 1.127 | 6.869 |
| 5% | PCR2645 | R9 | HAC | 7.8 | 0.291 | 1.73 |
| 2.50% | PCR2645 | R9 | HAC | 3.41 | 0.269 | 0.971 |
| 1.50% | PCR2645 | R9 | HAC | 1.205 | 0.137 | 1.026 |
| 1% | PCR2645 | R9 | HAC | 1.637 | 0.168 | 0.986 |
| 0.50% | PCR2645 | R9 | HAC | 0.873 | 0.132 | 0.327 |
| 100 % KIV-2B | PCR5104 | R9 | HAC | 100 | 0 | 0 |
| 5% | PCR5104 | R9 | HAC | 3.421 | 0.23 | 0.806 |
| 2.50% | PCR5104 | R9 | HAC | 1.446 | 0.031 | 0.24 |
| 1.50% | PCR5104 | R9 | HAC | 0.871 | 0.028 | 0.217 |
| 1% | PCR5104 | R9 | HAC | 0.609 | 0 | 0 |
| 0.50% | PCR5104 | R9 | HAC | 0.18 | 0 | 0 |
| 100 % KIV-2B | PCR2645 | V14 | HAC | 99.645 | 0.315 | 0.968 |
| 5% | PCR2645 | V14 | HAC | 4.756 | 0.277 | 1.412 |
| 2.50% | PCR2645 | V14 | HAC | 2.683 | 0.162 | 0.978 |
| 1.50% | PCR2645 | V14 | HAC | 2.289 | 0.106 | 0.216 |
| 1% | PCR2645 | V14 | HAC | 2.424 | 0.175 | 0.775 |
| 0.50% | PCR2645 | V14 | HAC | 1.502 | 0.151 | 0.355 |
| 100 % KIV-2B | PCR5104 | V14 | HAC | 99.996 | 0.023 | 0.128 |
| 5% | PCR5104 | V14 | HAC | 4.116 | 0.107 | 0.682 |
| 2.50% | PCR5104 | V14 | HAC | 1.624 | 0.068 | 0.24 |
| 1.50% | PCR5104 | V14 | HAC | 1.291 | 0.228 | 0.759 |
| 1% | PCR5104 | V14 | HAC | 0.89 | 0.057 | 0.45 |
| 0.50% | PCR5104 | V14 | HAC | 0.292 | 0 | 0 |
| 100 % KIV-2B | PCR2645 | V14 | SUP | 99.83 | 0.147 | 0.575 |
| 5% | PCR2645 | V14 | SUP | 3.582 | 0.298 | 1.517 |
| 2.50% | PCR2645 | V14 | SUP | 1.994 | 0.155 | 0.725 |
| 1.50% | PCR2645 | V14 | SUP | 2.519 | 0.113 | 0.433 |

| | | | | | | |
|---|---|---|---|---|---|---|
| 1% | PCR2645 | V14 | SUP | 2.424 | 0.17 | 0.752 |
| 0.50% | PCR2645 | V14 | SUP | 1.387 | 0.033 | 0.346 |
| 100 % KIV-2B | PCR5104 | V14 | SUP | 99.996 | 0.022 | 0.123 |
| 5% | PCR5104 | V14 | SUP | 2.89 | 0.185 | 0.443 |
| 2.50% | PCR5104 | V14 | SUP | 1.665 | 0.221 | 0.847 |
| 1.50% | PCR5104 | V14 | SUP | 1.505 | 0.035 | 0.274 |
| 1% | PCR5104 | V14 | SUP | 1.372 | 0.132 | 0.93 |
| 0.50% | PCR5104 | V14 | SUP | 0.578 | 0.053 | 0.578 |

**Supplementary table 13 Performance measures for all plasmid mixtures analyzed with by UMI-ONT-Seq with the cluster splitting strategy.**

| Sample | Fragment | Kit | Algorithm | Raw reads | Consensus sequences | True positive | True negative | False positive | False negative | Sensitivity | Specificity | Precision | F1 score |
|---|---|---|---|---|---|---|---|---|---|---|---|---|---|
| 100 % KIV-2A | PCR2645 | R9 | HAC | 134277 | 265 | 8 | 2637 | 0 | 0 | 1 | 1 | 1 | 1 |
| 100 % KIV-2B | PCR2645 | R9 | HAC | 109843 | 106 | 112 | 2533 | 0 | 0 | 1 | 1 | 1 | 1 |
| 5% | PCR2645 | R9 | HAC | 126532 | 321 | 118 | 2522 | 5 | 0 | 1 | 0.998 | 0.959 | 0.979 |
| 2.50% | PCR2645 | R9 | HAC | 113217 | 247 | 118 | 2526 | 1 | 0 | 1 | 1 | 0.992 | 0.996 |
| 1.50% | PCR2645 | R9 | HAC | 93180 | 71 | 114 | 2504 | 27 | 0 | 0.966 | 0.991 | 0.809 | 0.88 |
| 1% | PCR2645 | R9 | HAC | 101180 | 128 | 118 | 2520 | 7 | 0 | 1 | 0.997 | 0.944 | 0.971 |
| 0.50% | PCR2645 | R9 | HAC | 137340 | 249 | 118 | 2524 | 3 | 0 | 1 | 0.999 | 0.975 | 0.987 |
| 100 % KIV-2A | PCR5104 | R9 | HAC | 163591 | 613 | 27 | 5077 | 0 | 0 | 1 | 1 | 1 | 1 |
| 100 % KIV-2B | PCR5104 | R9 | HAC | 125709 | 514 | 75 | 5029 | 0 | 0 | 1 | 1 | 1 | 1 |
| 5% | PCR5104 | R9 | HAC | 127595 | 453 | 88 | 5008 | 8 | 0 | 1 | 0.998 | 0.917 | 0.957 |
| 2.50% | PCR5104 | R9 | HAC | 77479 | 211 | 88 | 5002 | 14 | 0 | 1 | 0.997 | 0.863 | 0.926 |
| 1.50% | PCR5104 | R9 | HAC | 106608 | 471 | 88 | 5015 | 1 | 0 | 1 | 1 | 0.989 | 0.994 |
| 1% | PCR5104 | R9 | HAC | 105730 | 347 | 88 | 5004 | 12 | 0 | 1 | 0.998 | 0.88 | 0.936 |
| 0.50% | PCR5104 | R9 | HAC | 152135 | 533 | 27 | 5016 | 0 | 61 | 0.307 | 1 | 1 | 0.47 |
| 100 % KIV-2A | PCR2645 | V14 | HAC | 96155 | 760 | 8 | 2637 | 0 | 0 | 1 | 1 | 1 | 1 |
| 100 % KIV-2B | PCR2645 | V14 | HAC | 105403 | 341 | 112 | 2533 | 0 | 0 | 1 | 1 | 1 | 1 |
| 5% | PCR2645 | V14 | HAC | 110703 | 898 | 118 | 2527 | 0 | 0 | 1 | 1 | 1 | 1 |
| 2.50% | PCR2645 | V14 | HAC | 105147 | 1047 | 118 | 2527 | 0 | 0 | 1 | 1 | 1 | 1 |
| 1.50% | PCR2645 | V14 | HAC | 94187 | 548 | 118 | 2527 | 0 | 0 | 1 | 1 | 1 | 1 |
| 1% | PCR2645 | V14 | HAC | 131673 | 432 | 118 | 2527 | 0 | 0 | 1 | 1 | 1 | 1 |
| 0.50% | PCR2645 | V14 | HAC | 91836 | 282 | 118 | 2526 | 1 | 0 | 1 | 1 | 0.992 | 0.996 |
| 100 % KIV-2A | PCR5104 | V14 | HAC | 104810 | 140 | 27 | 5077 | 0 | 0 | 1 | 1 | 1 | 1 |
| 100 % KIV-2B | PCR5104 | V14 | HAC | 145038 | 1051 | 75 | 5029 | 0 | 0 | 1 | 1 | 1 | 1 |
| 5% | PCR5104 | V14 | HAC | 107300 | 441 | 88 | 5016 | 0 | 0 | 1 | 1 | 1 | 1 |
| 2.50% | PCR5104 | V14 | HAC | 127751 | 893 | 88 | 5016 | 0 | 0 | 1 | 1 | 1 | 1 |
| 1.50% | PCR5104 | V14 | HAC | 110409 | 793 | 88 | 5016 | 0 | 0 | 1 | 1 | 1 | 1 |
| 1% | PCR5104 | V14 | HAC | 113702 | 202 | 88 | 5015 | 1 | 0 | 1 | 1 | 0.989 | 0.994 |
| 0.50% | PCR5104 | V14 | HAC | 94354 | 322 | 27 | 5016 | 1 | 60 | 0.307 | 1 | 0.964 | 0.466 |
| 100 % KIV-2A | PCR2645 | V14 | SUP | 96155 | 746 | 8 | 2637 | 0 | 0 | 1 | 1 | 1 | 1 |

| | | | | | | | | | | | | | |
|---|---|---|---|---|---|---|---|---|---|---|---|---|---|
| 100 % KIV-2B | PCR2645 | V14 | SUP | 110348 | 346 | 112 | 2533 | 0 | 0 | 1 | 1 | 1 | 1 |
| 5% | PCR2645 | V14 | SUP | 110703 | 893 | 118 | 2527 | 0 | 0 | 1 | 1 | 1 | 1 |
| 2.50% | PCR2645 | V14 | SUP | 105147 | 1010 | 118 | 2527 | 0 | 0 | 1 | 1 | 1 | 1 |
| 1.50% | PCR2645 | V14 | SUP | 94187 | 541 | 118 | 2527 | 0 | 0 | 1 | 1 | 1 | 1 |
| 1% | PCR2645 | V14 | SUP | 133275 | 436 | 118 | 2527 | 0 | 0 | 1 | 1 | 1 | 1 |
| 0.50% | PCR2645 | V14 | SUP | 91836 | 278 | 118 | 2526 | 1 | 0 | 1 | 1 | 0.992 | 0.996 |
| 100 % KIV-2A | PCR5104 | V14 | SUP | 106702 | 185 | 27 | 5077 | 0 | 0 | 1 | 1 | 1 | 1 |
| 100 % KIV-2B | PCR5104 | V14 | SUP | 179155 | 1206 | 75 | 5029 | 0 | 0 | 1 | 1 | 1 | 1 |
| 5% | PCR5104 | V14 | SUP | 109517 | 462 | 88 | 5016 | 0 | 0 | 1 | 1 | 1 | 1 |
| 2.50% | PCR5104 | V14 | SUP | 130071 | 1100 | 88 | 5016 | 0 | 0 | 1 | 1 | 1 | 1 |
| 1.50% | PCR5104 | V14 | SUP | 146157 | 932 | 88 | 5016 | 0 | 0 | 1 | 1 | 1 | 1 |
| 1% | PCR5104 | V14 | SUP | 115813 | 222 | 88 | 5015 | 1 | 0 | 1 | 1 | 0.989 | 0.994 |
| 0.50% | PCR5104 | V14 | SUP | 96259 | 332 | 27 | 5016 | 0 | 61 | 0.307 | 1 | 1 | 0.47 |

**Supplementary table 14 Noise reduction of detected variant levels of the UMI-ONT-Seq analysis pipeline using either the default clustering or the cluster splitting strategy.**

| Kit | Algorithm | Mean noise reduction [%] | SD noise reduction [%] | Median noise reduction [%] | Samples with no noise with default clustering strategy | Samples with no noise with cluster splitting strategy |
|---|---|---|---|---|---|---|
| R9 | HAC | 82.7 | 96.4 | 72.8 | 5 | 10 |
| V14 | HAC | 314.6 | 310.9 | 257.5 | 4 | 12 |
| V14 | SUP | 232.8 | 108.8 | 204.9 | 4 | 12 |

**Supplementary table 15 Variant levels and noise of mutation level of the UMI-ONT-Seq analysis pipeline using the cluster splitting strategy.**

| Sample | Fragment | Kit | Algorithm | Mean variant level [%] | SD variant level [%] | Absolute variant level range observed [%] |
|---|---|---|---|---|---|---|
| 100 % KIV-2B | PCR2645 | R9 | HAC | 99.691 | 1.395 | 8.491 |
| 5% | PCR2645 | R9 | HAC | 8.403 | 0.099 | 0.935 |
| 2.50% | PCR2645 | R9 | HAC | 4.844 | 0.076 | 0.405 |
| 1.50% | PCR2645 | R9 | HAC | 1.434 | 0.189 | 1.408 |
| 1% | PCR2645 | R9 | HAC | 3.125 | 0 | 0 |
| 0.50% | PCR2645 | R9 | HAC | 1.205 | 0 | 0 |
| 100 % KIV-2B | PCR5104 | R9 | HAC | 99.987 | 0.049 | 0.195 |
| 5% | PCR5104 | R9 | HAC | 4.415 | 0 | 0 |
| 2.50% | PCR5104 | R9 | HAC | 0.948 | 0 | 0 |
| 1.50% | PCR5104 | R9 | HAC | 1.274 | 0 | 0 |
| 1% | PCR5104 | R9 | HAC | 1.153 | 0 | 0 |
| 0.50% | PCR5104 | R9 | HAC | 0.188 | 0 | 0 |
| 100 % KIV-2B | PCR2645 | V14 | HAC | 99.979 | 0.077 | 0.293 |
| 5% | PCR2645 | V14 | HAC | 6.679 | 0.03 | 0.334 |
| 2.50% | PCR2645 | V14 | HAC | 3.059 | 0.016 | 0.096 |
| 1.50% | PCR2645 | V14 | HAC | 2.739 | 0.017 | 0.182 |
| 1% | PCR2645 | V14 | HAC | 3.009 | 0 | 0 |
| 0.50% | PCR2645 | V14 | HAC | 1.422 | 0.034 | 0.355 |
| 100 % KIV-2B | PCR5104 | V14 | HAC | 99.989 | 0.052 | 0.381 |
| 5% | PCR5104 | V14 | HAC | 4.085 | 0.029 | 0.227 |
| 2.50% | PCR5104 | V14 | HAC | 2.467 | 0.02 | 0.112 |
| 1.50% | PCR5104 | V14 | HAC | 1.515 | 0.016 | 0.126 |
| 1% | PCR5104 | V14 | HAC | 0.99 | 0 | 0 |
| 0.50% | PCR5104 | V14 | HAC | 0.326 | 0.088 | 0.621 |
| 100 % KIV-2B | PCR2645 | V14 | SUP | 99.982 | 0.071 | 0.289 |
| 5% | PCR2645 | V14 | SUP | 6.717 | 0.03 | 0.336 |
| 2.50% | PCR2645 | V14 | SUP | 2.875 | 0.023 | 0.198 |
| 1.50% | PCR2645 | V14 | SUP | 2.774 | 0.018 | 0.185 |
| 1% | PCR2645 | V14 | SUP | 2.752 | 0 | 0 |
| 0.50% | PCR2645 | V14 | SUP | 1.442 | 0.034 | 0.36 |
| 100 % KIV-2B | PCR5104 | V14 | SUP | 99.996 | 0.018 | 0.083 |

| | | | | | | |
|---|---|---|---|---|---|---|
| 5% | PCR5104 | V14 | SUP | 3.9 | 0.028 | 0.216 |
| 2.50% | PCR5104 | V14 | SUP | 2.37 | 0.037 | 0.273 |
| 1.50% | PCR5104 | V14 | SUP | 1.289 | 0.014 | 0.107 |
| 1% | PCR5104 | V14 | SUP | 1.802 | 0.082 | 0.901 |
| 0.50% | PCR5104 | V14 | SUP | 0.306 | 0.039 | 0.301 |

**Supplementary table 16 Threshold values for reaching a dataset Q-score of Q40. SD: Standard deviation**

| Kit | Algorithm | Result type | Minimal number of reads per cluster | Consensus sequences | Consensus sequences without error | Consensus sequences without error [%] | Mean Q-score per consensus | SD Q-score per consensus |
|---|---|---|---|---|---|---|---|---|
| R9 | HAC | Raw | 14 | 1220 | 747 | 61.23 | 38.336 | 2.41 |
| V14 | HAC | Raw | 10 | 2488 | 1552 | 62.379 | 38.094 | 2.76 |
| V14 | SUP | Raw | 6 | 3483 | 2174 | 62.417 | 38.168 | 2.681 |
| R9 | HAC | Transversion error corrected | 12 | 1470 | 908 | 61.769 | 38.347 | 2.427 |
| V14 | HAC | Transversion error corrected | 6 | 3271 | 2939 | 89.85 | 39.517 | 1.582 |
| V14 | SUP | Transversion error corrected | 6 | 3483 | 3327 | 95.521 | 39.795 | 1.02 |

**Supplementary table 17 Consensus sequence summary statistics for a minimal cluster size threshold of 6 reads per UMI cluster.**

| Kit | Algorithm | Total number of reads | Maximal number of errors | Reads with n or less errors | Percent of total reads | Number of bases[Gb] | Number of errors |
|---|---|---|---|---|---|---|---|
| R9 | HAC | 2997 | 0 | 1004 | 33.5 | 4.588 | 0 |
| V14 | HAC | 3271 | 0 | 1901 | 58.117 | 7.487 | 0 |
| V14 | SUP | 3483 | 0 | 2174 | 62.417 | 8.954 | 0 |
| R9 | HAC | 2997 | 2 | 2111 | 70.437 | 9.132 | 1522 |
| V14 | HAC | 3271 | 2 | 3168 | 96.851 | 12.619 | 1553 |
| V14 | SUP | 3483 | 2 | 3425 | 98.335 | 14.061 | 1487 |

**Supplementary table 18 Summary of the error profiles for all cluster thresholds greater than 10. ":" stands for insertion. "D" Deletion**

| Kit | Algorithm | Base change | Mean occurrence of error [%] | SD occurrence of error [%] |
|---|---|---|---|---|
| R9 | HAC | :A | 57.15% | 26.58% |
| R9 | HAC | AD | 15.18% | 20.32% |
| R9 | HAC | CA | 22.71% | 20.02% |
| R9 | HAC | GT | 15.70% | 21.24% |

| | | | | |
|---|---|---|---|---|
| V14 | HAC | CA | 62.37% | 5.69% |
| V14 | HAC | GT | 31.80% | 8.81% |
| V14 | SUP | CA | 63.64% | 5.27% |
| V14 | SUP | GT | 31.31% | 7.22% |

**Supplementary table 19 Dataset Q-scores per sequencing chemistry and basecalling algorithm after error-profile adjustment.**

| Kit | Algorithm | Minimal number of reads per cluster | Cluster | Nucleotides | Errors | Dataset Q-score |
|---|---|---|---|---|---|---|
| R9 | HAC | 2 | 8639 | 35221384 | 724239 | 16.869 |
| R9 | HAC | 4 | 4102 | 16375163 | 24585 | 28.235 |
| R9 | HAC | 6 | 2997 | 12459002 | 5693 | 33.401 |
| R9 | HAC | 8 | 2294 | 9974981 | 2315 | 36.344 |
| R9 | HAC | 10 | 1827 | 8247966 | 1272 | 38.119 |
| R9 | HAC | 12 | 1470 | 6902884 | 772 | 39.514 |
| R9 | HAC | 14 | 1220 | 5833440 | 541 | 40.327 |
| R9 | HAC | 16 | 1039 | 5049779 | 410 | 40.905 |
| R9 | HAC | 18 | 882 | 4356647 | 321 | 41.326 |
| R9 | HAC | 20 | 755 | 3769914 | 253 | 41.732 |
| V14 | HAC | 2 | 11806 | 46875946 | 502112 | 19.701 |
| V14 | HAC | 4 | 4415 | 18346483 | 5428 | 35.289 |
| V14 | HAC | 6 | 3271 | 13082913 | 501 | 44.169 |
| V14 | HAC | 8 | 2806 | 11019387 | 163 | 48.3 |
| V14 | HAC | 10 | 2488 | 9679100 | 92 | 50.22 |
| V14 | HAC | 12 | 2238 | 8636705 | 72 | 50.79 |
| V14 | HAC | 14 | 2023 | 7743442 | 57 | 51.331 |
| V14 | HAC | 16 | 1837 | 6995736 | 52 | 51.288 |
| V14 | HAC | 18 | 1649 | 6232904 | 47 | 51.226 |
| V14 | HAC | 20 | 1483 | 5587278 | 42 | 51.24 |
| V14 | SUP | 2 | 13316 | 52615786 | 454575 | 20.635 |
| V14 | SUP | 4 | 4825 | 20493221 | 2649 | 38.885 |
| V14 | SUP | 6 | 3483 | 14295288 | 192 | 48.719 |
| V14 | SUP | 8 | 2947 | 11852165 | 94 | 51.007 |
| V14 | SUP | 10 | 2625 | 10464413 | 76 | 51.389 |
| V14 | SUP | 12 | 2389 | 9468884 | 70 | 51.312 |
| V14 | SUP | 14 | 2203 | 8708883 | 64 | 51.338 |
| V14 | SUP | 16 | 2047 | 8082330 | 63 | 51.082 |
| V14 | SUP | 18 | 1873 | 7388495 | 50 | 51.696 |
| V14 | SUP | 20 | 1684 | 6618100 | 46 | 51.58 |

**Supplementary table 20 Performance measures for all human gDNA samples for all sequencing chemistries and basecalling conditions.**

| Sample | Fragment | Kit | Algorithm | Raw reads | Consensus sequences | True positive | True negative | False positive | False negative | Sensitivity | Specificity | Precision | F1 score |
|---|---|---|---|---|---|---|---|---|---|---|---|---|---|
| EUR15415 | 5104 | R9 | HAC | 70038 | 17 | 44 | 4973 | 4 | 83 | 0.346 | 0.999 | 0.917 | 0.503 |
| EUR27513 | 5104 | R9 | HAC | 88489 | 142 | 107 | 4978 | 9 | 10 | 0.915 | 0.998 | 0.922 | 0.918 |
| EUR28877 | 5104 | R9 | HAC | 156027 | 487 | 132 | 4966 | 4 | 2 | 0.985 | 0.999 | 0.971 | 0.978 |
| EUR32880 | 5104 | R9 | HAC | 151916 | 251 | 65 | 4971 | 3 | 65 | 0.5 | 0.999 | 0.956 | 0.657 |
| EUR35729 | 5104 | R9 | HAC | 95958 | 3 | 18 | 4999 | 1 | 86 | 0.173 | 1 | 0.947 | 0.293 |
| EUR41297 | 5104 | R9 | HAC | 128104 | 81 | 52 | 5031 | 19 | 2 | 0.963 | 0.996 | 0.732 | 0.832 |
| EUR60915 | 5104 | R9 | HAC | 110100 | 4 | 32 | 5037 | 0 | 35 | 0.478 | 1 | 1 | 0.646 |
| EUR63113 | 5104 | R9 | HAC | 157136 | 692 | 145 | 4942 | 16 | 1 | 0.993 | 0.997 | 0.901 | 0.945 |
| EUR63453 | 5104 | R9 | HAC | 114891 | 34 | 103 | 4980 | 3 | 18 | 0.851 | 0.999 | 0.972 | 0.907 |
| EUR66900 | 5104 | R9 | HAC | 93850 | 253 | 77 | 4943 | 22 | 62 | 0.554 | 0.996 | 0.778 | 0.647 |
| EUR73204 | 5104 | R9 | HAC | 208536 | 106 | 129 | 4868 | 105 | 2 | 0.985 | 0.979 | 0.551 | 0.707 |
| EUR77692 | 5104 | R9 | HAC | 147495 | 537 | 108 | 4990 | 4 | 2 | 0.982 | 0.999 | 0.964 | 0.973 |
| EUR86619 | 5104 | R9 | HAC | 97587 | 331 | 99 | 5003 | 0 | 2 | 0.98 | 1 | 1 | 0.99 |
| EUR98778 | 5104 | R9 | HAC | 114148 | 529 | 122 | 4976 | 2 | 4 | 0.968 | 1 | 0.984 | 0.976 |
| EUR15415 | 5104 | V14 | HAC | 107148 | 450 | 126 | 4975 | 1 | 2 | 0.984 | 1 | 0.992 | 0.988 |
| EUR21943 | 5104 | V14 | HAC | 115224 | 148 | 61 | 5039 | 1 | 3 | 0.953 | 1 | 0.984 | 0.968 |
| EUR27513 | 5104 | V14 | HAC | 90414 | 423 | 113 | 4985 | 2 | 4 | 0.966 | 1 | 0.983 | 0.974 |
| EUR28877 | 5104 | V14 | HAC | 106609 | 740 | 131 | 4970 | 0 | 3 | 0.978 | 1 | 1 | 0.989 |
| EUR32880 | 5104 | V14 | HAC | 103952 | 508 | 124 | 4974 | 0 | 6 | 0.954 | 1 | 1 | 0.976 |
| EUR35729 | 5104 | V14 | HAC | 100376 | 142 | 100 | 5000 | 0 | 4 | 0.962 | 1 | 1 | 0.98 |
| EUR41297 | 5104 | V14 | HAC | 114752 | 477 | 53 | 5048 | 1 | 2 | 0.964 | 1 | 0.981 | 0.972 |
| EUR60915 | 5104 | V14 | HAC | 101977 | 81 | 51 | 5029 | 8 | 16 | 0.761 | 0.998 | 0.864 | 0.81 |
| EUR63113 | 5104 | V14 | HAC | 100513 | 680 | 145 | 4943 | 14 | 2 | 0.986 | 0.997 | 0.912 | 0.948 |
| EUR63453 | 5104 | V14 | HAC | 99587 | 266 | 117 | 4982 | 1 | 4 | 0.967 | 1 | 0.992 | 0.979 |
| EUR66900 | 5104 | V14 | HAC | 112271 | 717 | 137 | 4962 | 2 | 3 | 0.979 | 1 | 0.986 | 0.982 |
| EUR73204 | 5104 | V14 | HAC | 157966 | 290 | 128 | 4967 | 6 | 3 | 0.977 | 0.999 | 0.955 | 0.966 |
| EUR77692 | 5104 | V14 | HAC | 102688 | 632 | 108 | 4994 | 0 | 2 | 0.982 | 1 | 1 | 0.991 |
| EUR86619 | 5104 | V14 | HAC | 116328 | 743 | 98 | 5002 | 1 | 3 | 0.97 | 1 | 0.99 | 0.98 |

| | | | | | | | | | | | | | |
|---|---|---|---|---|---|---|---|---|---|---|---|---|---|
| EUR98778 | 5104 | V14 | HAC | 159463 | 733 | 122 | 4978 | 0 | 4 | 0.968 | 1 | 1 | 0.984 |
| EUR15415 | 5104 | V14 | SUP | 109926 | 784 | 126 | 4976 | 0 | 2 | 0.984 | 1 | 1 | 0.992 |
| EUR21943 | 5104 | V14 | SUP | 117770 | 316 | 61 | 5040 | 0 | 3 | 0.953 | 1 | 1 | 0.976 |
| EUR27513 | 5104 | V14 | SUP | 106423 | 653 | 113 | 4987 | 0 | 4 | 0.966 | 1 | 1 | 0.983 |
| EUR28877 | 5104 | V14 | SUP | 108763 | 959 | 131 | 4970 | 0 | 3 | 0.978 | 1 | 1 | 0.989 |
| EUR32880 | 5104 | V14 | SUP | 106256 | 773 | 124 | 4974 | 0 | 6 | 0.954 | 1 | 1 | 0.976 |
| EUR35729 | 5104 | V14 | SUP | 102554 | 317 | 100 | 5000 | 0 | 4 | 0.962 | 1 | 1 | 0.98 |
| EUR41297 | 5104 | V14 | SUP | 117428 | 807 | 53 | 5049 | 0 | 2 | 0.964 | 1 | 1 | 0.981 |
| EUR60915 | 5104 | V14 | SUP | 104458 | 220 | 63 | 5036 | 1 | 4 | 0.94 | 1 | 0.984 | 0.962 |
| EUR63113 | 5104 | V14 | SUP | 119810 | 809 | 145 | 4945 | 12 | 2 | 0.986 | 0.998 | 0.924 | 0.954 |
| EUR63453 | 5104 | V14 | SUP | 101851 | 532 | 116 | 4983 | 0 | 5 | 0.959 | 1 | 1 | 0.979 |
| EUR66900 | 5104 | V14 | SUP | 138525 | 887 | 137 | 4964 | 0 | 3 | 0.979 | 1 | 1 | 0.989 |
| EUR73204 | 5104 | V14 | SUP | 261885 | 469 | 128 | 4966 | 7 | 3 | 0.977 | 0.999 | 0.948 | 0.962 |
| EUR77692 | 5104 | V14 | SUP | 124413 | 816 | 108 | 4994 | 0 | 2 | 0.982 | 1 | 1 | 0.991 |
| EUR86619 | 5104 | V14 | SUP | 125810 | 860 | 98 | 5003 | 0 | 3 | 0.97 | 1 | 1 | 0.985 |
| EUR98778 | 5104 | V14 | SUP | 177455 | 786 | 122 | 4978 | 0 | 4 | 0.968 | 1 | 1 | 0.984 |
| EUR15415 | 5104 | V14 | SUPDUP | 29849 | 7 | 30 | 4976 | 0 | 98 | 0.234 | 1 | 1 | 0.38 |
| EUR28877 | 5104 | V14 | SUPDUP | 30581 | 17 | 54 | 4969 | 1 | 80 | 0.403 | 1 | 0.982 | 0.571 |
| EUR32880 | 5104 | V14 | SUPDUP | 28577 | 8 | 31 | 4973 | 2 | 98 | 0.24 | 1 | 0.939 | 0.383 |
| EUR41297 | 5104 | V14 | SUPDUP | 34236 | 7 | 26 | 5048 | 1 | 29 | 0.473 | 1 | 0.963 | 0.634 |
| EUR63113 | 5104 | V14 | SUPDUP | 11125 | 27 | 124 | 4918 | 40 | 22 | 0.849 | 0.992 | 0.756 | 0.8 |
| EUR86619 | 5104 | V14 | SUPDUP | 12618 | 5 | 23 | 4999 | 4 | 78 | 0.228 | 0.999 | 0.852 | 0.359 |
| EUR98778 | 5104 | V14 | SUPDUP | 17411 | 81 | 118 | 4965 | 13 | 8 | 0.937 | 0.997 | 0.901 | 0.918 |

**Supplementary table 21 Summary performance measures for UMI-ONT-Seq.**

| Kit | Algorithm | Samples | Sensitivity Mean | Sensitivity SD | Specificity Mean | Specificity SD | Precision Mean | Precision SD | F1 score Mean | F1 score SD |
|---|---|---|---|---|---|---|---|---|---|---|
| R9 | HAC | 14 | 0.762 | 0.288 | 0.997 | 0.005 | 0.9 | 0.128 | 0.784 | 0.214 |
| V14 | HAC | 15 | 0.957 | 0.055 | 1 | 0.001 | 0.976 | 0.039 | 0.966 | 0.045 |
| V14 | SUP | 15 | 0.968 | 0.013 | 1 | 0.001 | 0.99 | 0.023 | 0.979 | 0.011 |
| V14 | SUPDUP | 7 | 0.481 | 0.298 | 0.998 | 0.003 | 0.913 | 0.086 | 0.578 | 0.221 |

**Supplementary table 22 Summary performance measures for ONT-Seq without UMIs.**

| Kit | Algorithm | Samples | Sensitivity Mean | Sensitivity SD | Specificity Mean | Specificity SD | Precision Mean | Precision SD | F1 score Mean | F1 score SD |
|---|---|---|---|---|---|---|---|---|---|---|
| R9 | HAC | 14 | 0.614 | 0.3 | 0.595 | 0.191 | 0.106 | 0.221 | 0.073 | 0.058 |
| V14 | HAC | 15 | 0.89 | 0.049 | 0.837 | 0.007 | 0.111 | 0.027 | 0.196 | 0.043 |
| V14 | SUP | 15 | 0.799 | 0.317 | 0.889 | 0.021 | 0.132 | 0.06 | 0.225 | 0.1 |
| V14 | SUPDUP | 8 | 0.95 | 0.031 | 0.971 | 0.008 | 0.399 | 0.122 | 0.554 | 0.123 |

**Supplementary table 23 Differences and correlation values between ddPCR measured and UMI-ONT-Seq predicted number of KIV-2 repeats for different prediction strategies. CI: Confidence interval**

| Strategy | Mean difference | SD difference | r | $R^2$ | Samples within ddPCR CI [n] |
|---|---|---|---|---|---|
| All unique haplotypes | +9.059 | 8.15 | 0.842 | 0.709 | 3 |
| Merged unique haplotypes | -3.588 | 3.94 | 0.961 | 0.924 | 5 |
| Coverage-corrected | +0.412 | 2.891 | 0.975 | 0.950 | 10 |

**Supplementary table 24 KIV-2 subtype occurrence in the SAPHIR and 1000G dataset.**

| Population | Super Population | Samples | KIV-2 subtype | Subtype occurrence | Relative subtype occurrence |
|---|---|---|---|---|---|
| CDX | EAS | 12 | KIV-2B | 2 | 0.167 |
| CDX | EAS | 12 | KIV-2C | 6 | 0.5 |
| CDX | EAS | 12 | only KIV-2A | 4 | 0.333 |
| JPT | EAS | 12 | KIV-2B | 4 | 0.333 |
| JPT | EAS | 12 | KIV-2C | 6 | 0.5 |
| JPT | EAS | 12 | only KIV-2A | 2 | 0.167 |
| PJL | SAS | 12 | KIV-2B | 6 | 0.5 |
| PJL | SAS | 12 | KIV-2C | 3 | 0.25 |
| PJL | SAS | 12 | only KIV-2A | 3 | 0.25 |
| SAPHIR | EUR | 15 | KIV-2B | 8 | 0.533 |
| SAPHIR | EUR | 15 | KIV-2C | 4 | 0.267 |
| SAPHIR | EUR | 15 | only KIV-2A | 3 | 0.2 |
| YRI | AFR | 12 | KIV-2B | 8 | 0.667 |
| YRI | AFR | 12 | only KIV-2A | 4 | 0.333 |

**Supplementary table 25 KIV-2 subtype specific summary performance measures for UMI-ONT-Seq analysis of the 48 1000G samples.**

| Samples | KIV-2 subtype | Sensitivity Mean | Sensitivity SD | Specificity Mean | Specificity SD | Precision Mean | Precision SD | F1 score Mean | F1 score SD |
|---|---|---|---|---|---|---|---|---|---|
| 35 | contains KIV-2B and/or KIV-2C | 0.995 | 0.007 | 1 | 0.001 | 0.983 | 0.022 | 0.989 | 0.013 |
| 13 | only KIV-2A | 0.592 | 0.047 | 1 | 0 | 0.988 | 0.009 | 0.74 | 0.037 |

**Supplementary table 26 Per-sample performance measures for UMI-ONT-Seq analysis of the 48 1000G samples for V14 SUP.**

| Sample | Population | KIV-2 subtype | Raw reads | Consensus sequences | True positive | True negative | False positive | False negative | Sensitivity | Specificity | Precision | F1 score |
|---|---|---|---|---|---|---|---|---|---|---|---|---|
| HG00766 | CDX | contains KIV-2B and/or KIV-2C | 180551 | 937 | 114 | 4990 | 0 | 0 | 1 | 1 | 1 | 1 |
| HG01802 | CDX | contains KIV-2B and/or KIV-2C | 212541 | 1272 | 143 | 4958 | 2 | 1 | 0.993 | 1 | 0.986 | 0.99 |
| HG01811 | CDX | contains KIV-2B and/or KIV-2C | 310581 | 1802 | 114 | 4989 | 1 | 0 | 1 | 1 | 0.991 | 0.996 |
| HG02154 | CDX | contains KIV-2B and/or KIV-2C | 203792 | 1269 | 143 | 4960 | 1 | 0 | 1 | 1 | 0.993 | 0.997 |
| HG02184 | CDX | contains KIV-2B and/or KIV-2C | 200983 | 1190 | 145 | 4958 | 1 | 0 | 1 | 1 | 0.993 | 0.997 |
| HG02375 | CDX | contains KIV-2B and/or KIV-2C | 149627 | 963 | 149 | 4954 | 1 | 0 | 1 | 1 | 0.993 | 0.997 |
| HG02394 | CDX | contains KIV-2B and/or KIV-2C | 146497 | 684 | 108 | 4993 | 3 | 0 | 1 | 0.999 | 0.973 | 0.986 |
| HG02406 | CDX | contains KIV-2B and/or KIV-2C | 231893 | 1410 | 136 | 4966 | 1 | 1 | 0.993 | 1 | 0.993 | 0.993 |
| HG01805 | CDX | only KIV-2A | 218346 | 1313 | 58 | 5004 | 1 | 41 | 0.586 | 1 | 0.983 | 0.734 |
| HG02355 | CDX | only KIV-2A | 172259 | 919 | 56 | 5003 | 1 | 44 | 0.56 | 1 | 0.982 | 0.713 |
| HG02371 | CDX | only KIV-2A | 132332 | 883 | 63 | 4997 | 1 | 43 | 0.594 | 1 | 0.984 | 0.741 |
| HG02385 | CDX | only KIV-2A | 202281 | 1191 | 60 | 4997 | 0 | 47 | 0.561 | 1 | 1 | 0.719 |
| NA18942 | JPT | contains KIV-2B and/or KIV-2C | 191416 | 1052 | 131 | 4972 | 1 | 0 | 1 | 1 | 0.992 | 0.996 |
| NA18945 | JPT | contains KIV-2B and/or KIV-2C | 185226 | 920 | 110 | 4994 | 0 | 0 | 1 | 1 | 1 | 1 |

| | | | | | | | | | | | | |
|---|---|---|---|---|---|---|---|---|---|---|---|---|
| NA18950 | JPT | contains KIV-2B and/or KIV-2C | 192303 | 1153 | 137 | 4965 | 2 | 0 | 1 | 1 | 0.986 | 0.993 |
| NA18987 | JPT | contains KIV-2B and/or KIV-2C | 182461 | 839 | 146 | 4956 | 2 | 0 | 1 | 1 | 0.986 | 0.993 |
| NA19007 | JPT | contains KIV-2B and/or KIV-2C | 182972 | 822 | 140 | 4960 | 0 | 4 | 0.972 | 1 | 1 | 0.986 |
| NA19056 | JPT | contains KIV-2B and/or KIV-2C | 165790 | 904 | 112 | 4992 | 0 | 0 | 1 | 1 | 1 | 1 |
| NA19058 | JPT | contains KIV-2B and/or KIV-2C | 75303 | 94 | 112 | 4983 | 8 | 1 | 0.991 | 0.998 | 0.933 | 0.961 |
| NA19066 | JPT | contains KIV-2B and/or KIV-2C | 117345 | 471 | 149 | 4952 | 3 | 0 | 1 | 0.999 | 0.98 | 0.99 |
| NA19072 | JPT | contains KIV-2B and/or KIV-2C | 115109 | 686 | 142 | 4960 | 2 | 0 | 1 | 1 | 0.986 | 0.993 |
| NA19074 | JPT | contains KIV-2B and/or KIV-2C | 99583 | 556 | 102 | 5001 | 1 | 0 | 1 | 1 | 0.99 | 0.995 |
| NA18944 | JPT | only KIV-2A | 132767 | 863 | 63 | 4993 | 2 | 46 | 0.578 | 1 | 0.969 | 0.724 |
| NA19002 | JPT | only KIV-2A | 230404 | 1371 | 64 | 4997 | 1 | 42 | 0.604 | 1 | 0.985 | 0.749 |
| HG02490 | PJL | contains KIV-2B and/or KIV-2C | 134239 | 754 | 107 | 4996 | 1 | 0 | 1 | 1 | 0.991 | 0.995 |
| HG02652 | PJL | contains KIV-2B and/or KIV-2C | 94659 | 510 | 115 | 4986 | 2 | 1 | 0.991 | 1 | 0.983 | 0.987 |
| HG02655 | PJL | contains KIV-2B and/or KIV-2C | 144664 | 632 | 95 | 5008 | 1 | 0 | 1 | 1 | 0.99 | 0.995 |
| HG02684 | PJL | contains KIV-2B and/or KIV-2C | 139851 | 720 | 124 | 4969 | 10 | 1 | 0.992 | 0.998 | 0.925 | 0.958 |
| HG02737 | PJL | contains KIV-2B and/or KIV-2C | 148915 | 931 | 115 | 4989 | 0 | 0 | 1 | 1 | 1 | 1 |
| HG03021 | PJL | contains KIV-2B and/or KIV-2C | 131982 | 736 | 114 | 4989 | 1 | 0 | 1 | 1 | 0.991 | 0.996 |
| HG03652 | PJL | contains KIV-2B and/or KIV-2C | 188154 | 837 | 127 | 4974 | 3 | 0 | 1 | 0.999 | 0.977 | 0.988 |
| HG03653 | PJL | contains KIV-2B and/or KIV-2C | 158362 | 948 | 136 | 4967 | 1 | 0 | 1 | 1 | 0.993 | 0.996 |
| HG03702 | PJL | contains KIV-2B and/or KIV-2C | 201185 | 1171 | 110 | 4993 | 1 | 0 | 1 | 1 | 0.991 | 0.995 |
| HG02694 | PJL | only KIV-2A | 158195 | 1065 | 63 | 5000 | 1 | 40 | 0.612 | 1 | 0.984 | 0.754 |
| HG03238 | PJL | only KIV-2A | 180456 | 1158 | 70 | 4982 | 1 | 51 | 0.579 | 1 | 0.986 | 0.729 |

| | | | | | | | | | | | | |
|---|---|---|---|---|---|---|---|---|---|---|---|---|
| HG03708 | PJL | only KIV-2A | 310158 | 1251 | 60 | 4997 | 1 | 46 | 0.566 | 1 | 0.984 | 0.719 |
| NA18516 | YRI | contains KIV-2B and/or KIV-2C | 181110 | 1222 | 113 | 4990 | 1 | 0 | 1 | 1 | 0.991 | 0.996 |
| NA18873 | YRI | contains KIV-2B and/or KIV-2C | 153257 | 774 | 142 | 4950 | 10 | 2 | 0.986 | 0.998 | 0.934 | 0.959 |
| NA18907 | YRI | contains KIV-2B and/or KIV-2C | 347909 | 1941 | 160 | 4937 | 4 | 3 | 0.982 | 0.999 | 0.976 | 0.979 |
| NA19092 | YRI | contains KIV-2B and/or KIV-2C | 264116 | 769 | 107 | 4995 | 1 | 1 | 0.991 | 1 | 0.991 | 0.991 |
| NA19171 | YRI | contains KIV-2B and/or KIV-2C | 386363 | 1120 | 130 | 4960 | 12 | 2 | 0.985 | 0.998 | 0.915 | 0.949 |
| NA19235 | YRI | contains KIV-2B and/or KIV-2C | 170053 | 706 | 144 | 4956 | 2 | 2 | 0.986 | 1 | 0.986 | 0.986 |
| NA19239 | YRI | contains KIV-2B and/or KIV-2C | 181384 | 517 | 96 | 5006 | 0 | 2 | 0.98 | 1 | 1 | 0.99 |
| NA19248 | YRI | contains KIV-2B and/or KIV-2C | 207469 | 1197 | 139 | 4962 | 1 | 2 | 0.986 | 1 | 0.993 | 0.989 |
| NA18510 | YRI | only KIV-2A | 215364 | 1250 | 58 | 4989 | 0 | 57 | 0.504 | 1 | 1 | 0.671 |
| NA18861 | YRI | only KIV-2A | 229313 | 1095 | 98 | 4964 | 0 | 42 | 0.7 | 1 | 1 | 0.824 |
| NA18916 | YRI | only KIV-2A | 270688 | 1135 | 77 | 4984 | 1 | 42 | 0.647 | 1 | 0.987 | 0.782 |
| NA19201 | YRI | only KIV-2A | 189399 | 922 | 73 | 4984 | 0 | 47 | 0.608 | 1 | 1 | 0.756 |

**Supplementary table 27 Summary performance measures for UMI-ONT-Seq analysis of the 48 1000G samples without KIV-2B specific variants.**

| Samples | Sensitivity Mean | Sensitivity SD | Specificity Mean | Specificity SD | Precision Mean | Precision SD | F1 score Mean | F1 score SD |
|---|---|---|---|---|---|---|---|---|
| 48 | 0.987 | 0.018 | 1 | 0.001 | 0.979 | 0.033 | 0.983 | 0.02 |

**Supplementary table 28 Correlation between the number of KIV-2 repeats quantified by ddPCR and by UMI-ONT-Seq for each step of the UMI-ONT-Seq KIV-2 quantification algorithm in the 1000G dataset. The confidence interval (CI) is set to 95 %. CI: Confidence interval**

| Strategy | Mean difference | SD difference | r | $R^2$ | Within ddPCR CI [n] |
|---|---|---|---|---|---|
| Unique | +3.833 | 7.413 | 0.68 | 0.462 | 10 |
| Merged | -6.125 | 5.895 | 0.672 | 0.451 | 16 |
| Coverage corrected | +0.292 | 4.371 | 0.851 | 0.724 | 32 |

**Supplementary table 29 extracted STR sequences of all 63 human gDNA samples and two unmixed KIV-2A, respectively KIV-2B plasmids.**

| STR sequence | STR length | Occurrence (KIV-2 repeats) [n] |
|---|---|---|
| CACACACACACACACACACACACACACACACA | 16 | 15504 |
| CACACACACACACACACACACACACA | 13 | 15367 |
| CACACACACACACACACACACACACACACA | 15 | 9701 |
| CACACACACACACACACACACACACACA | 14 | 6005 |
| CACACACACACACACACACACACA | 12 | 4056 |
| CACACAGACACACACACACACACA | 12 | 3844 |
| CACACACACACACACACACA | 10 | 2018 |
| CACACACACACACAGACACACACACACACACA | 16 | 1546 |
| CACACACACACACACACACACACACACACACACA | 17 | 1433 |
| CACACACACACACACACACACA | 11 | 1411 |
| CACACACACACACACACACACACACACACACACACACA | 19 | 500 |
| CACACACACACACACACACACACACACACACACACACACA | 20 | 429 |
| CACACACACACACACACACACACACACACACACACA | 18 | 414 |
| CACACACACACACACACA | 9 | 97 |
| CACACAGACACACACACACACA | 11 | 67 |
| CACACACACACACACACACACACACACACACACACACACACA | 21 | 50 |
| CACACACAAACACACACACACACACACACACACACACA | 18 | 33 |
| CACACACAAACACACACACACACACACACACACACACACA | 19 | 33 |
| CACACAAACACACACACACACACA | 12 | 30 |
| CACACAGACACACACACACACACACA | 13 | 30 |
| CACACAGACACACACACACACACACACA | 14 | 27 |
| CACACACAAACACACACACACACACACACACACACA | 17 | 24 |
| CACACACACACACAGACACACACACACACA | 15 | 22 |
| CACACAGCACACACACACACACA | 11.5 | 20 |
| CACACACACACACACACACACACACACACACACACACACACACA | 22 | 4 |
| CACACACACACACAGCACACACACACACACA | 15.5 | 4 |
| CACACACACACAGACACACACACACACACA | 15 | 3 |
| CACACAGACACACACACACA | 10 | 3 |
| CACACACAACACACACACACACACACACACACACA | 17.5 | 2 |
| CACACACACACACACA | 8 | 2 |

**Supplementary table 30 Population-specific degeneration of the STR region.**

| Population | Super population | Mutation | Mutation is at STR position | Occurrence [n samples] | Occurrence [n sequences] |
|---|---|---|---|---|---|
| YRI | AFR | none | - | 12 | 12529 |
| YRI | AFR | AA | 8 | 1 | 90 |
| YRI | AFR | A | 2 | 1 | 3 |
| CDX | EAS | none | - | 12 | 12032 |
| CDX | EAS | GA | 7 | 9 | 1362 |
| CDX | EAS | GA | 15 | 9 | 417 |
| CDX | EAS | G | 7 | 4 | 13 |
| JPT | EAS | none | - | 12 | 8541 |
| JPT | EAS | GA | 7 | 8 | 729 |
| JPT | EAS | GA | 15 | 7 | 444 |
| JPT | EAS | G | 7 | 1 | 2 |
| SAPHIR | EUR | none | - | 15 | 12926 |
| SAPHIR | EUR | GA | 7 | 8 | 886 |
| SAPHIR | EUR | GA | 15 | 7 | 435 |
| SAPHIR | EUR | AA | 6 | 2 | 31 |
| SAPHIR | EUR | G | 7 | 2 | 5 |
| SAPHIR | EUR | G | 15 | 1 | 2 |
| PJL | SAS | none | - | 12 | 9426 |
| PJL | SAS | GA | 7 | 7 | 994 |
| PJL | SAS | GA | 15 | 6 | 272 |
| PLASMID | EUR | none | - | 2 | 1537 |

**Supplementary table 31 Diversity of the extracted STR sequences per population.**

| Population | Super population | Unique STR sequences |
|---|---|---|
| PLASMID | EUR | 4 |
| CDX | EAS | 18 |
| YRI | AFR | 18 |
| PJL | SAS | 20 |
| JPT | EAS | 22 |
| SAPHIR | EUR | 32 |

**Supplementary table 32 Predicted number of KIV-2 repeats for 48 samples of the 1000G project using either UMI-ONT-Seq, ddPCR or the results from Behera et al. [9]**

| Sample | Population | Super population | ddPCR predicted number of KIV-2 repeats | UMI-ONT-Seq predicted number of KIV-2 repeats | Behera et al predicted number of KIV-2 repeats |
|---|---|---|---|---|---|
| NA18510 | YRI | AFR | 37 | 35 | 38 |
| NA18873 | YRI | AFR | 31 | 29 | 31 |
| NA18861 | YRI | AFR | 35 | 35 | 37 |
| NA19092 | YRI | AFR | 24 | 26 | 27 |
| NA19235 | YRI | AFR | 35 | 36 | 37 |
| NA19201 | YRI | AFR | 22 | 22 | 23 |
| NA18916 | YRI | AFR | 33 | 35 | 35 |
| NA19171 | YRI | AFR | 33 | 34 | 34 |
| NA19248 | YRI | AFR | 36 | 38 | 38 |
| NA19239 | YRI | AFR | 20 | 23 | 23 |
| NA18907 | YRI | AFR | 36 | 38 | 37 |
| NA18516 | YRI | AFR | 27 | 32 | 31 |
| HG02371 | CDX | EAS | 48 | 42 | 50 |
| HG02385 | CDX | EAS | 45 | 42 | 48 |
| HG02355 | CDX | EAS | 34 | 32 | 35 |
| HG00766 | CDX | EAS | 47 | 49 | 52 |
| HG02375 | CDX | EAS | 47 | 48 | 50 |
| HG02406 | CDX | EAS | 41 | 38 | 40 |
| HG02184 | CDX | EAS | 42 | 47 | 49 |
| HG01811 | CDX | EAS | 52 | 53 | 54 |
| HG01805 | CDX | EAS | 47 | 48 | 49 |
| HG02394 | CDX | EAS | 34 | 36 | 36 |
| HG01802 | CDX | EAS | 38 | 41 | 41 |
| HG02154 | CDX | EAS | 51 | 52 | 52 |
| NA19074 | JPT | EAS | 43 | 23 | 45 |
| NA19058 | JPT | EAS | 40 | 30 | 48 |
| NA19056 | JPT | EAS | 50 | 41 | 48 |
| NA18950 | JPT | EAS | 43 | 44 | 48 |
| NA19002 | JPT | EAS | 52 | 50 | 53 |
| NA19007 | JPT | EAS | 39 | 37 | 40 |
| NA18987 | JPT | EAS | 41 | 40 | 42 |
| NA19066 | JPT | EAS | 43 | 44 | 45 |
| NA18945 | JPT | EAS | 38 | 42 | 42 |
| NA18942 | JPT | EAS | 41 | 43 | 43 |
| NA18944 | JPT | EAS | 42 | 46 | 46 |
| NA19072 | JPT | EAS | 46 | 50 | 47 |
| HG03021 | PJL | SAS | 37 | 38 | 40 |
| HG03708 | PJL | SAS | 43 | 44 | 46 |
| HG03653 | PJL | SAS | 41 | 41 | 42 |

| | | | | | |
|---|---|---|---|---|---|
| HG02490 | PJL | SAS | 34 | 34 | 35 |
| HG03652 | PJL | SAS | 36 | 41 | 41 |
| HG02694 | PJL | SAS | 41 | 43 | 43 |
| HG02655 | PJL | SAS | 26 | 28 | 28 |
| HG02684 | PJL | SAS | 32 | 33 | 33 |
| HG02737 | PJL | SAS | 28 | 31 | 30 |
| HG03238 | PJL | SAS | 42 | 44 | 43 |
| HG03702 | PJL | SAS | 45 | 51 | 50 |
| HG02652 | PJL | SAS | 30 | 33 | 32 |
